# SPICEMIX: Integrative single-cell spatial modeling of cell identity

**DOI:** 10.1101/2020.11.29.383067

**Authors:** Benjamin Chidester, Tianming Zhou, Shahul Alam, Jian Ma

## Abstract

Spatial transcriptomics technologies promise to reveal spatial relationships of cell-type composition in complex tissues. However, the development of computational methods that can utilize the unique properties of spatial transcriptome data to unveil cell identities remains a challenge. Here, we introduce SpiceMix, a new interpretable method based on probabilistic, latent variable modeling for effective joint analysis of spatial information and gene expression from spatial transcriptome data. Both simulation and real data evaluations demonstrate that SpiceMix markedly improves upon the inference of cell types and their spatial patterns compared with existing approaches. By applying to spatial transcriptome data of brain regions in human and mouse acquired by seqFISH+, STARmap, and Visium, we show that SpiceMix can enhance the inference of complex cell identities, reveal interpretable spatial metagenes, and uncover differentiation trajectories. SpiceMix is a generalizable framework for analyzing spatial transcriptome data to provide critical insights into the cell type composition and spatial organization of cells in complex tissues.

## Introduction

The compositions of different cell types in mammalian tissues, such as brain, remain poorly understood due to the complex interplay among intrinsic, spatial, and temporal factors that collectively contribute to the identify of a cell [1–3]. Single-cell RNA-seq (scRNA-seq) has greatly advanced our understanding of complex cell types in different tissues [4–6], but its utility in disentangling spatial factors in particular is inherently limited by the dissociation of cells from their spatial context. The emerging spatial transcriptomics technologies based on multiplexed imaging and sequencing [7–18] are able to reveal spatial information of gene expression of dozens to tens of thousands of genes in individual cells *in situ* within the tissue context. However, the development of computational methods that can incorporate the unique properties of spatially resolved transcriptome data to unveil cell identities and spatially variable features remains a challenge [19, 20].

Several methods have been developed for the analysis of spatial transcriptome data to reveal spatial domains and programs of cell types in tissues [21–25], to explore the spatial variance of genes [26–29], and to align scRNA-seq with spatial transcriptome data [30–33]. Both probabilistic graphical models, such as methods using hidden Markov random field (HMRF), and graph-based neural network architectures have been proposed [20]. The conventional HMRF, which is commonly used to model spatial dependencies and is the underlying approach of Zhu et al. [22] as well as the more recent BayesSpace [25] have two major limitations for modeling cell identity: (1) it assumes that cell types or spatial domains are discrete, and therefore it cannot model the interplay of intrinsic and extrinsic factors that give rise to cell identity; and (2) it assumes that cell types or domains exhibit smooth spatial patterns, which is often not true for many cell types, such as the spatial pattern of inhibitory neurons that sparsely populate tissue. More recently, graph convolution neural networks have also been used for learning spatial domains, such as SpaGCN [23]. The drawbacks of such methods for modeling cell identity are that their learned latent representations are not easily interpretable and they are more susceptible to overfitting, in comparison to effective linear latent variable models for scRNA-seq data, such as non-negative matrix factorization (NMF). In addition, the existing methods typically do not integrate the modeling of the spatial variability of genes with their contribution to cell identity [23, 26, 29]. Therefore, there is an urgent need for robust, interpretable methods that can jointly model both the spatial and intrinsic factors of cell identity, which is of vital importance to fully utilize spatial transcriptome data.

Here, we introduce SpiceMix (Spatial Identification of Cells using Matrix Factorization), a new interpretable and integrative framework to model spatial transcriptome data. SpiceMix uses latent variable modeling to express the interplay of spatial and intrinsic factors that collectively contribute to cell identity. Crucially, SpiceMix enhances the NMF [34] model of gene expression with a novel integration of a graphical representation of the spatial relationship of cells. Thus, the spatial patterns learned by SpiceMix can elucidate the relationship between intrinsic and spatial factors, leading to more meaningful representations of cell identity. Applications to the spatial transcriptome datasets of brain regions in human and mouse acquired by seqFISH+ [12], STARmap [13], and Visium [18] demonstrate that SpiceMix can enhance the inference of complex cell identities, uncover spatially variable features, and reveal important biological processes. Notably, the interpretable metagenes from SpiceMix uniquely unveil spatially variable gene expression patterns of important cell types. SpiceMix has the potential to provide critical new insights into the spatial composition of cells based on spatial transcriptome data.

## Results

### Overview of SpiceMix

SpiceMix is an integrative solution for spatial transciptomic analysis to provide key insights into the spatially variable features of cell composition in complex tissues. Specifically, SpiceMix models spatial transcriptome data by a new probabilistic graphical model NMF-HMRF (**Fig.** 1). Our model has a natural interpretation for single-cell spatial transcriptome data, where each node in the graph represents a cell and edges capture cell-to-cell relationships of nearby cells, but it can also be applied to *in situ* sequencing-based methods (e.g., Visium [10]), where each node represents a spatially barcoded spot.

**Figure 1:**
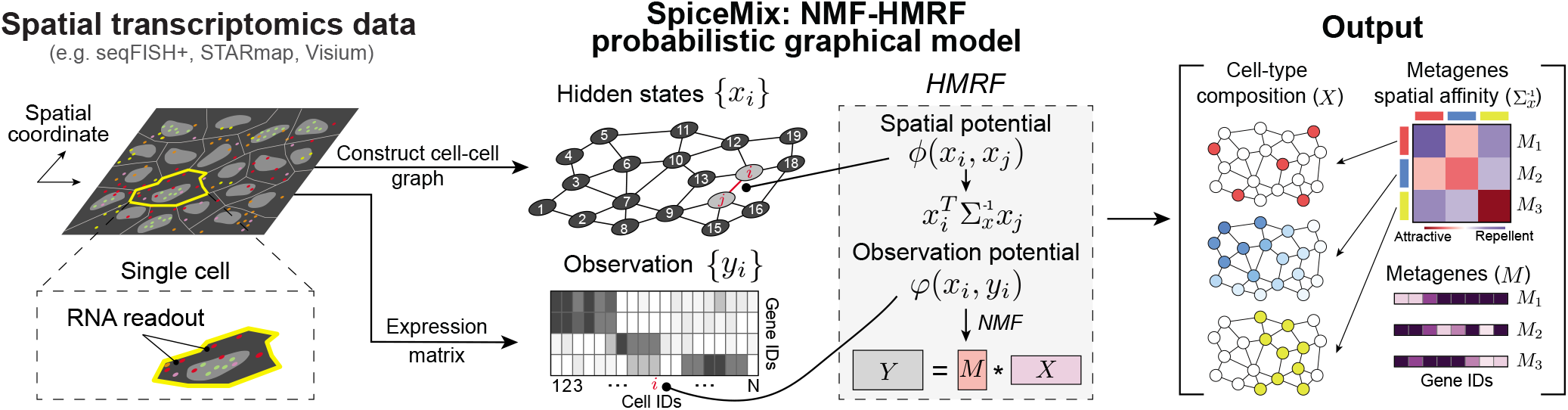
Overview of SpiceMix. Gene expression measurements and a neighbor graph are extracted from spatial trancriptome data and fed into the SpiceMix framework. SpiceMix decomposes the expression *y_i_* in cell (or spot) *i* into a mixture of metagenes weighted by the hidden state *x_i_*. Spatial interaction between neighboring cells (or spots) *i* and *j* is modeled by an inner product of their hidden states, weighted by 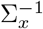, the inferred spatial affinities between metagenes. The hidden mixture weights *X*, the metagene spatial affinity 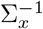, and *K* metagenes *M*, all inferred by SpiceMix, provide unique insight into the spatially variable features that collectively constitute the identity of each cell.

For each node *i* in the graphical model NMF-HMRF, a latent state vector *x_i_* represents the mixture of weights for *K* different intrinsic or extrinsic factors (**Fig.** 1), which collectively constitute the identity of the cell. In order to capture the continuous nature of cell state, our model extends the standard HMRF model by allowing these latent states to be continuous. Importantly, different types of correlations of latent states in nearby cells are captured by the matrix 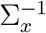 (**Fig.** 1), which, unlike a conventional HMRF and many other spatial models, does not assume smooth spatial patterns only, but has the flexibility to represent both the smooth and sparse spatial patterns that compose real tissue. Each element of the *K* × *K* matrix 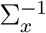 represents the pair-wise affinity between two factors, providing an intuitive interpretation of the spatial patterns of cells in tissue. For each factor, a “metagene” in the *G* × *K* matrix *M* captures the expression of its associated genes (**Fig.** 1), where *G* denotes the number of total genes. The observed expression from spatial transcriptome data, *y_i_* = *M_xi_* for node *i*, follows a robust linear mixing model that affords a natural interpretation of its relationship to the various factors of cell identity from the associated genes. Thus, the novel NMF-HMRF model in SpiceMix is able to uniquely integrate the spatial modeling of the HMRF with the NMF formulation for gene expression into a single model for spatial transcriptome data.

Given an input spatial transcriptome dataset, SpiceMix simultaneously learns the intuitive metagenes *M* of latent factors, the latent states *X* for all nodes, and their spatial affinity 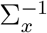. This is achieved by a novel alternating *maximum-a-posteriori* optimization algorithm. Importantly, in SpiceMix, metagenes are an integral part of the model outcome, which presents a methodological advance in comparison to the calculation of spatially variable genes as a post-processing step in other recent methods (such as SpaGCN [23]). A regularizing parameter allows users to control the weight given to the spatial information during optimization to suit the input data. The detailed description of the NMF-HMRF model is provided in the **Methods** section with additional steps of optimization in **Supplementary Methods**.

### Performance evaluation using simulated spatial transcriptome data

We first evaluated SpiceMix using simulation designed to model the mouse cortex, a region which has served as a prominent case study for many spatial transcriptomic methods (**Fig.** 2a-b; see **Methods** for the simulation method details). We devised two methods of generating expression based on the position and type of each cell: Approach I: We followed a metagene-based simulation; Approach II: We used scDesign2 [35] trained on real scRNA-seq data [36]. For Approach II, we introduced two forms of spatial influence over expression: leakage, which randomly swaps some reads of neighboring cells, to mimic challenges of processing real spatial transcriptomics data; and additive noise that follows random, spatially-smooth patterns. We compared the results from SpiceMix to that of NMF, HMRF, Seurat [37], and the very recent SpaGCN [23]. We evaluate the performance of different methods by comparing the inferred cell types with the true cell types using the adjusted Rand index (ARI) metric. For SpiceMix and NMF, we further applied Louvain clustering to the learned latent representations. The approaches for preprocessing the data and for choosing other hyperparameters for each method are provided in **Supplementary Methods** A.2.3.

**Figure 2:**
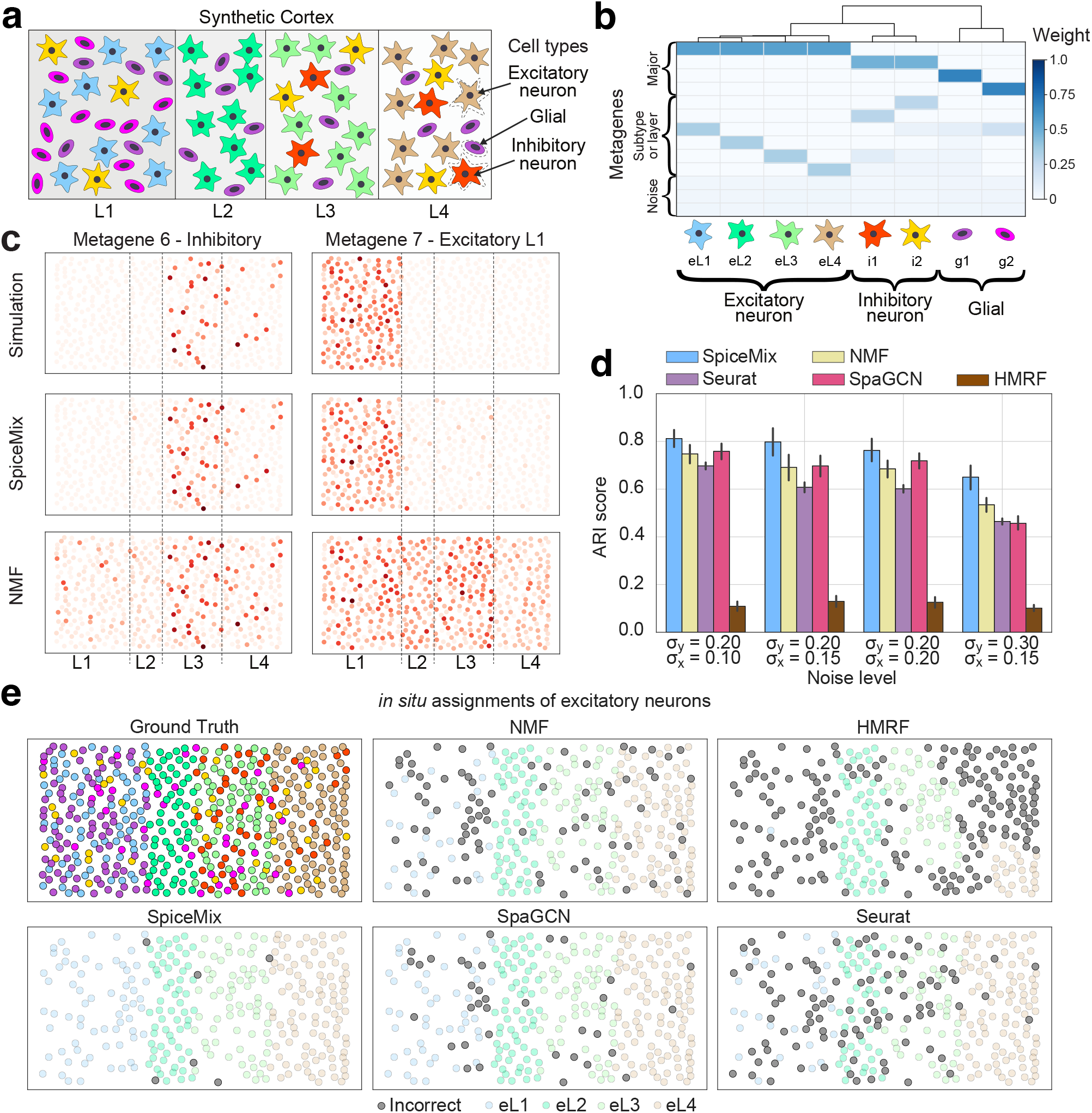
Performance evaluation based on simulated spatial transcriptome data. **a.** Illustration of the simulated spatial transcriptome data of the mouse cortex, including 3 major cell types distributed in 4 layers. Excitatory and inhibitory neurons are star-shaped and glial cells are ovals. Subtypes are distinguished by their colors. **b.** Dendrogram showing the similarity of the expression profiles of the 8 subtypes (top), their metagene profiles (middle), and their colors and shapes (bottom) used in panel **(a)**. The top 4 rows correspond to metagenes that determine major type, the next 6 rows correspond to metagenes that determine subtypes or are layer-specific, and the bottom 3 rows correspond to noise metagenes. **c.** Simulated expression of metagenes 6 and 7, from a single sample generated with *σ_y_* = 0.2 and *σ_x_* = 0.15, in their spatial context (top) and the inferred expression of those metagenes by SpiceMix and NMF. Visualizations in panel **(e)** are of the same simulated sample. **d.** Performance comparison of SpiceMix, NMF, HMRF, Seurat, and SpaGCN. Bar plots of the adjusted Rand index (ARI) score, that measures the matching between the identified cell types and the true cell types, are shown. Results are reported across four simulation scenarios with varying degrees of randomness. Error bars show +/- one standard deviation. **e.** Imputed cell-type labels of each method for the excitatory neurons, shown in their spatial context. Neurons that were correctly identified are colored faintly. Neurons that were incorrectly identified are colored dark gray. The upper left panel is the ground truth cell type of all cells in the simulated sample. The colors match those of panels **(a)** and **(b)**.

For both simulation approaches, we found that SpiceMix consistently outperformed other methods. For Approach I, SpiceMix achieved the highest average ARI scores (0.65-0.82) across all scenarios. For lower noise scenarios (*σ_y_* = 0.2), the ARI of SpiceMix was 9-18% higher than that of SpaGCN or NMF, which performed comparably (**Fig.** 2d). SpiceMix, SpaGCN, and NMF all outperformed Seurat and HMRF. For the higher noise setting (*σ_y_* = 0.3), SpiceMix significantly outperformed all methods (**Fig.** 2d). We found that SpiceMix was able to recover both the layer-specific and sparse metagenes that underlie the identity of cells. For example, SpiceMix successfully recovered metagene 7, which is specific to layer L1 (**Fig.** 2c) and is enriched in eL1 excitatory neurons (blue in **Fig.** 2a). Notably, SpiceMix was able to reveal nearly all excitatory neurons (**Fig.** 2e). SpiceMix also recovered metagene 6 (**Fig.** 2c), which captures intrinsic factors of the sparse inhibitory neuron subtype i1 (red in **Fig.** 2a). In contrast, the equivalent of metagene 7 for NMF is strongly expressed across layers L1-L3 (**Fig.** 2c), and NMF confused some eL3 excitatory neurons (light green) with eL1 excitatory neurons (**Fig.** 2e). The equivalent of metagene 6 for NMF shows a more diffuse pattern (**Fig.** 2c). Additional evaluation by varying the parameter λ_x_ or zero-thresholding to reflect different sparsity of the latent variables of NMF further demonstrated the robust advantage of SpiceMix (**Fig.** S1). In addition, SpaGCN, Seurat, and HMRF all incorrectly assigned the spatial patterns for many more excitatory neurons (**Fig.** 2e).

For simulation Approach II (using scDesign2), SpiceMix performed the best for all but one scenario, for which it tied with NMF, and the advantage of SpiceMix became more significant as the influence of noise and leakage on spatial expression patterns became more prevalent (see **Supplementary Results** B.1 and **Fig.** S2a). We found that the spatial metagenes from SpiceMix reliably reflect both cell type composition and spatial noise (**Fig.** S2b). Overall, SpiceMix achieved much more accurate spatial assignments of cells than all other methods (**Fig.** S2c).

Taken together, we showed that the novel integration of matrix factorization and spatial modeling in SpiceMix yields better inference of the underlying spatially variable features and cell identities as compared to existing methods. This improvement was found for cell types with either sparse or layer-specific spatial patterns, both of which are prevalent in real data from complex tissues (e.g., the brain region data used in this work). Our evaluation also confirmed the effectiveness and robustness of our new optimization scheme for applying the SpiceMix model to spatial transcriptome data.

### SpiceMix refines cell identity modeling of seqFISH+ data

We applied SpiceMix to a recent single-cell spatial transcriptomic dataset from mouse brain acquired by seqFISH+ [12]. Here, we used the data of five separate samples of the mouse primary visual cortex, all from the same mouse but from contiguous layers, each from a distinct image or field-of-view (FOV), with single-cell expression of 2,470 genes in 523 cells [12]. We compared the spatial patterns revealed by SpiceMix to those produced by NMF (given its relatively strong performance based on our simulation evaluation) with various levels of sparsity via *λ_x_* and zero-thresholding. We also compared to the results from Louvain clustering (**Supplementary Methods** A.3.3) and the HMRF-based method of Zhu et al. [22] as originally reported in Eng et al. [12]. In addition, SpiceMix revealed spatially-informed metagenes capturing biological processes in the cortex (see **Supplementary Table**).

To interpret the learned latent representation from SpiceMix, we first clustered the cells based on the latent representation using hierarchical clustering, which led to five excitatory neural subtypes, two inhibitory neural subtypes, and eight glial types (**Fig.** 3a). These cell type assignments were supported by known marker genes from a scRNA-seq study [36] (**Fig.** 3c (left) and **Supplementary Method** A.3.4). The assignment of major types is generally consistent among SpiceMix, NMF, and the Louvain clustering result from Eng et al. [12] (**Fig.** 3c (middle), **Fig.** S3a, and **Fig.** S4a (middle)). However, SpiceMix uncovered more refined cell states compared to any other method. Specifically, SpiceMix identified three distinct stages of oligodendrocyte maturation, from oligodendrocyte precurser cells (OPCs) to mature oligodendrocytes, throughout the five FOVs, as reflected by the expression of spatially-informed metagenes. Oligodendrocytes follow a maturation process starting with OPCs and progressing to myelin-sheath forming cells [38]. As annotated in **Fig.** 3c (right), metagene 7 is expressed at a high proportion among oligodendrocytes, distinguishing them from OPCs, while the expression of metagene 8, which is also present in OPCs, separates a cluster of early-stage oligodendrocytes (Oligo-E) from intermediate or late-stage oligodendrocytes (Oligo-L). The separation of these types is supported by the expression patterns of the OPC marker gene *Cspg4*, the differentiating oligodendrocyte marker gene *Tcf7l2* [39], and the mature oligodendrocyte marker gene *Mog* [40] (**Fig.** 3c (left)), as well as the analysis of 40 marker genes for oligodendrocyte stages from [38] (**Fig.** S6). This result suggests that metagenes 7 and 8 collectively capture the progression of oligodendrocytes along their maturation trajectory. In addition, metagene 7 has a strong spatial affinity with metagenes 3 and 4 (highlighted by black arrows in **Fig.** 3b; different from metagene 8), which are expressed primarily by the excitatory neurons of deeper tissue layers (eL5, eL6a, and eL6b) (**Fig.** 3c (right)). None of the other methods, including NMF, the Louvain clustering reported by [12], and the HMRF-based method of Zhu et al. [22] as reported in [12], could clearly distinguish these spatially-distinct cells (**Fig.** S4a,c, **Fig.** 3c (middle), **Fig.** S3b). Note that different sparsity constraints on NMF did not yield these oligodendrocyte stages either (**Supplementary Results** B.2 and **Fig.** S7).

**Figure 3:**
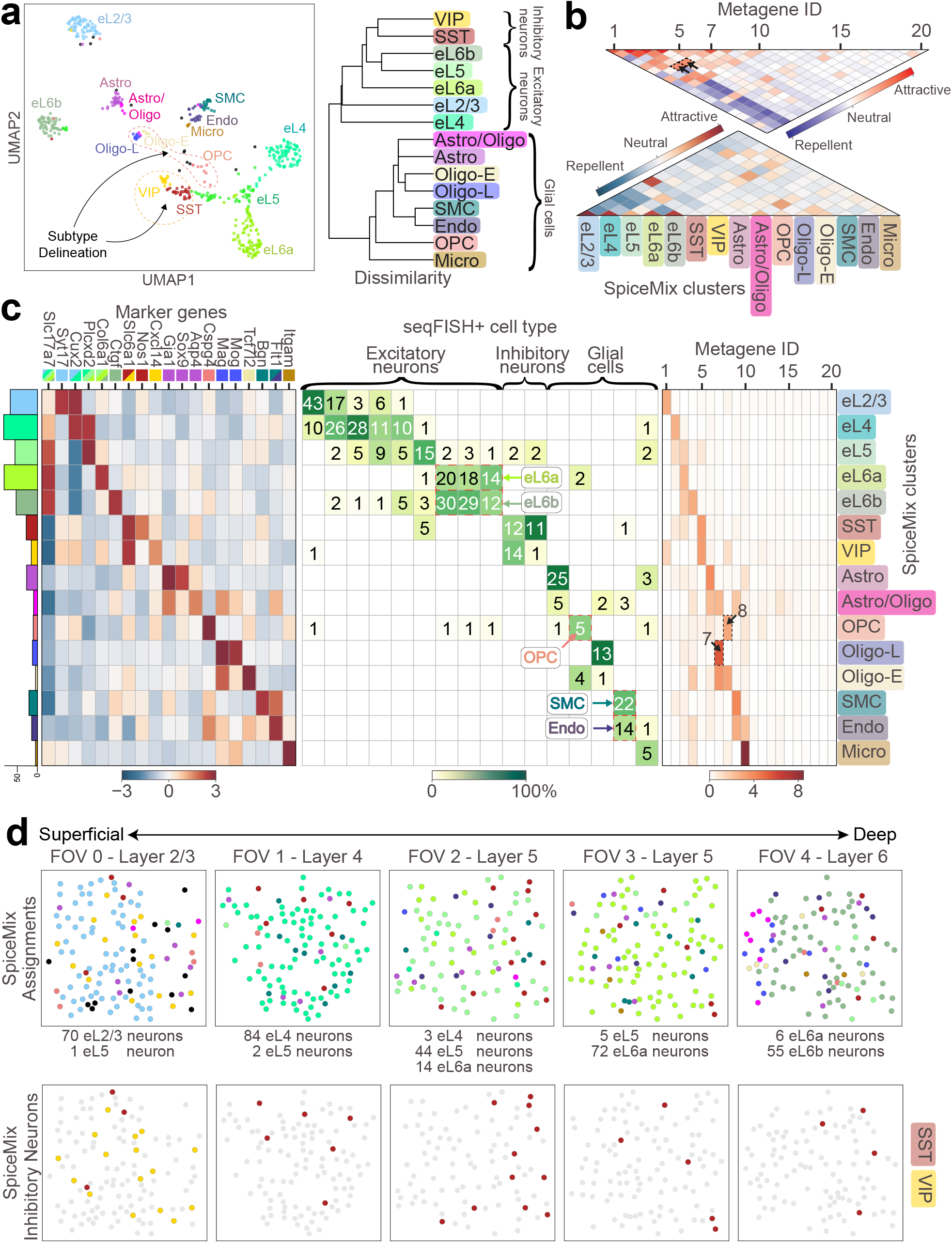
Application of SpiceMix to the seqFISH+ data from the mouse primary visual cortex [12]. Note that colors throughout the figure of cells and labels correspond to the cell-type assignments of SpiceMix. **a.** UMAP plot of the latent states of SpiceMix (left) and the dendrogram of the arithmetic average of the expression for each cell type of SpiceMix (right). It is highlighted in **(a)** (left) that SpiceMix further delineated inhibitory neurons into VIPs (yellow) and SSTs (red-brown) enclosed by the orange dashed cycle and refined oligodendrocytes and OPCs into separate subtypes: Astro/Oligo (magenta), Oligo-1 (beige), Oligo-2 (blue), and OPC (coral), enclosed within the red dashed cycle. **b.** (Top) The inferred pairwise spatial affinity of metagenes, or 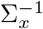. (Bottom) The inferred pairwise spatial affinity of SpiceMix cell types. **c.** (Left) Average expression of known marker genes within SpiceMix cell types, along with the number of cells belonging to each type (colored bar plot). The colored boxes on the top following the name of each marker gene correspond to their known associated cell type. (Middle) Agreement of SpiceMix cell-type assignments with those of the original analysis in [12]. (Right) Average expression of inferred metagenes within SpiceMix cell types. **d.** *In situ* SpiceMix cell-type assignments for all cells in each of the five FOVs (upper panel). *In situ* maps showing spatial enrichment of SpiceMix inhibitory neuron types (lower panel). Samples from superficial layers are on the left and samples from deep tissue layers are on the right. Colors of cell types are the same as in above panels.

SpiceMix also discovered spatially variable features that matched those reported from scRNA-seq studies [36, 41] (**Fig.** 3d). The assignment of excitatory neurons by SpiceMix showed strong layerenrichment patterns (**Fig.** 3d (top)), and the expression of known marker genes within these cell types matched that of a prior scRNA-seq study [36] (**Fig.** S5). SpiceMix also revealed a separation of VIP and SST neurons, with VIP neuron subtypes residing primarily in layers 1-4 and SST neurons diffusely populating all layers (**Fig.** 3d (bottom); VIP in yellow and SST in red-brown. This spatial pattern follows the proportion of VIP and SST neurons in brain layers as observed in a large-scale scRNA-seq study (see Fig. 1 in [41]), whereas the results of Louvain clustering did not exhibit such a pattern as reported in [12] (Fig. 3h in [12]). In addition, the expression of marker genes such as *Nos1* and *Cxcl14* [36] (**Fig.** 3c (left)) further supports this VIP/SST separation made by SpiceMix. NMF could not distinguish these inhibitory subtypes (**Fig.** S3a). Furthermore, the split of eL6 neurons by SpiceMix, which was not revealed by [12], is also supported by marker gene expression (**Supplementary Results** B.2 and **Fig.** S5).

Together, our analysis of seqFISH+ data of the mouse cortex with SpiceMix revealed spatially variable features and more refined cell states. Our results strongly demonstrate the advantages and unique capabilities of SpiceMix.

### SpiceMix reveals spatially variable metagenes and cell types from STARmap data

Next, we applied SpiceMix to a single-cell spatial transcriptome dataset of the mouse cortex acquired by STARmap [13]. We analyzed a sample of the mouse V1 neocortex consisting of 930 cells passing quality control, all from a single image or FOV, with expression measurements for 1020 genes. We mainly compared the results of SpiceMix, NMF, and Wang et al. [13]. To distinguish cell-type labels between methods, we append an asterisk to the end of the cell labels of Wang et al. [13] when referenced. In addition, SpiceMix generated spatially variable metagenes (see **Supplementary Table**).

We found that SpiceMix produced more accurate cell labels than [13] and revealed cell types missed by other methods (**Fig.** 4a, **Fig.** S8). Specifically, SpiceMix revealed two eL6 excitatory neuron subtypes, clearer layer-wise patterns of excitatory neurons, two spatially-distinct astrocyte types (Astro-1 and Astro-2), a maturation process of oligodendrocytes from Opcs to the mature Oligo-1 type, and a separate Oligo-2 type (**Fig.** 4a, c; supported by known marker genes [13, 36]).

**Figure 4:**
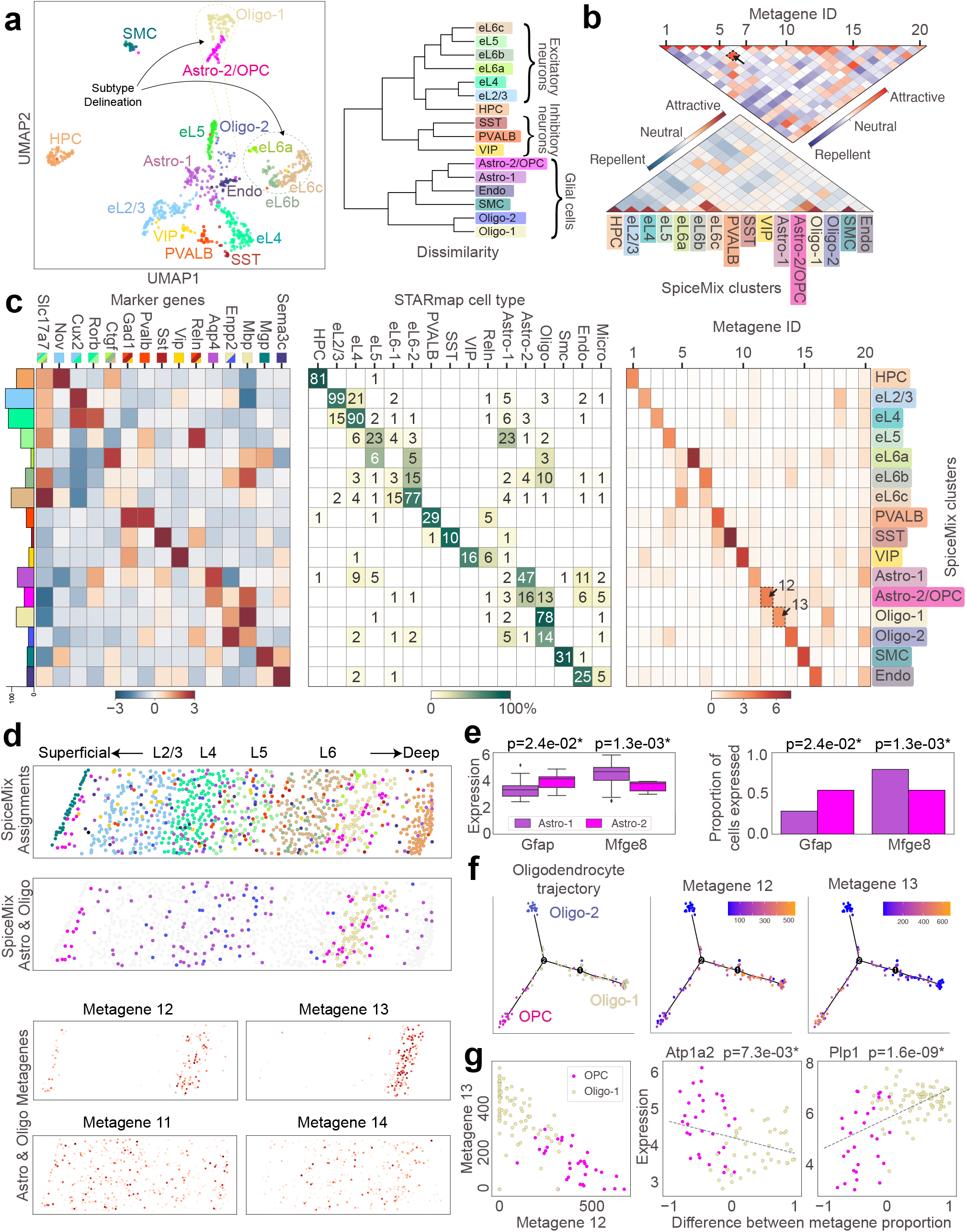
Application of SpiceMix to the STARmap data from mouse primary visual cortex [13]. Note that colors throughout the figure of cells and labels correspond to the cell-type assignments of SpiceMix. **a.** UMAP plots of the latent states of SpiceMix and the dendrogram of the arithmetic average of the expression for each cell type of SpiceMix (right). It is highlighted in **a** (left) that SpiceMix delineated eL6 neurons into three subtypes enclosed in the dark green cycle and delineated oligodendrocytes and Opcs into three separate subtypes: Oligo-1 (light yellow), Oligo-2 (silver), and Astro-2/OPC (magenta), enclosed within the red dashed cycle. **b.** (Top) The inferred pairwise spatial affinity of metagenes, or 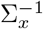. (Bottom) The inferred pairwise spatial affinity of cell types. **c.** (Left) Average expression of known marker genes within SpiceMix cell types, along with the number of cells belonging to each type (colored bar plot). The colored boxes on the top following the name of each marker gene correspond to their known associated cell types. (Middle) Agreement of SpiceMix cell-type assignments with those of the original analysis in [13]. (Right) Average expression of inferred metagenes within SpiceMix cell types. **d.** (Top) *In situ* map of SpiceMix cell-type assignments for all cells (upper) and astrocyte and oligodendrocyte cells (lower) in the sample. (Bottom) *In situ* maps of expression of both layer-specific and ubiquitous metagenes learned by SpiceMix that are relevant to astrocytes and oligodendrocytes. **e.** The expression of astrocyte subtype marker genes in Astro-1 and Astro-2 types of SpiceMix. *: ***P***<0.05. **f.** Trajectory analysis of SpiceMix oligodendrocyte types using Monocle2, showing the expression of metagenes 12 and 13 along the trajectory from OPC to Oligo-1. **g.** (Left) The expression of metagene 13 plotted against the expression of metagene 12 for oligodendrocytes of the SpiceMix Oligo-1 and OPC types. (Right) The expression of important marker genes for myelin-sheath formation in oligodendrocytes plotted against the relative expression of metagenes 12 and 13 of the same cells. The title of each plot consists of the gene symbol and the corrected *P*-value of having a nonzero slope, respectively. *: ***P***<0.05.

SpiceMix achieved a refined, spatially-informed separation of eL6 subtypes. SpiceMix identified metagenes 5 and 7 that collectively separate the excitatory subtypes eL6b and eL6c (**Fig.** 4c (right)). Metagenes 5 and 7 exhibit strong pairwise affinity (highlighted by a black arrow in **Fig.** 4b). Importantly, the patterns of SpiceMix excitatory neurons delineated clear layer boundaries (**Fig.** 4d), which matched enrichment analysis from scRNA-seq studies (see Fig. 4b in [41]). In contrast, the assignments reported in Fig. 5d in [13] showed significant mixing of excitatory types across layer boundaries. Comparison of marker genes from [36] for eL2/3 and eL4 showed that, among eL2/3 and eL4 neurons that were differently assigned between SpiceMix and [13], their expression levels in SpiceMix assignments more closely followed that of [36] (**Supplementary Results** B.3 and **Fig.** S9). SpiceMix also refined the assignment of a large set of cells from the Astro-1* type of [13] to eL5 (**Fig.** 4c (middle), d), which was confirmed by the expression of known excitatory marker genes (**Supplementary Results** B.3 and **Fig.** S10). Note that NMF corrected some assignments of excitatory neurons and also changed the assignment of this group labeled astrocytes to eL5; however, the resulting layers were not as clear as those of SpiceMix (**Fig.** S8). We also found that HMRF missed sparse cell types and smoothed across layers, missing even the layer-wise structure of excitatory neurons (**Fig.** S11).

**Figure 5:**
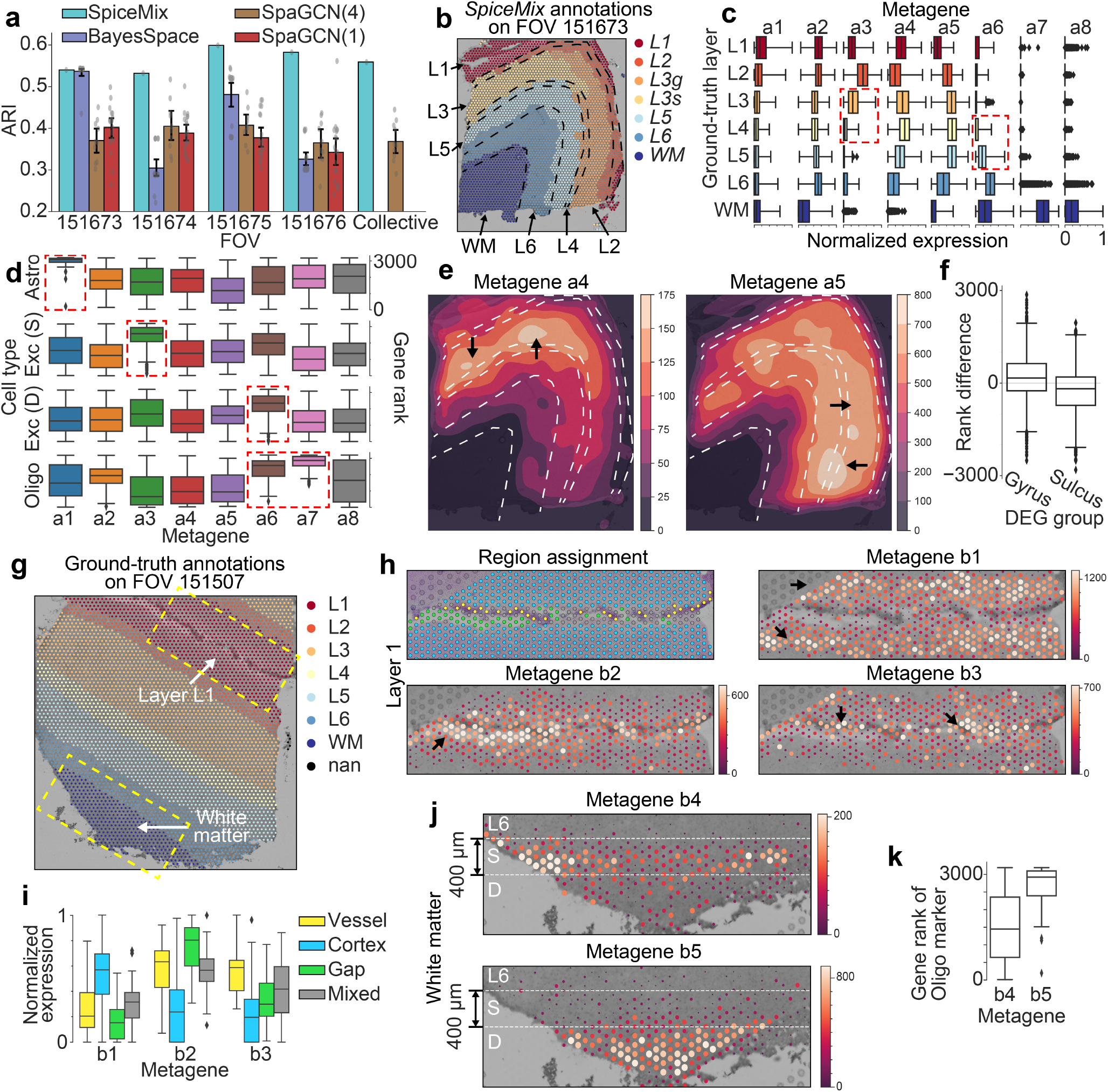
Application of SpiceMix to the Visium dataset on human dorsolateral prefrontal cortex [18]. SpiceMix was trained on the 4 FOVs 151673-151676 from sample Br8100 at the same time (**a-f**), and on FOV 151507 from sample Br5292 (**g-k**). Metagenes from the two runs are labeled with prefix ‘a’ and ‘b’, respectively. **a.** Comparison of the performance of SpiceMix, BayesSpace, and SpaGCN, on the 4 FOVs from sample Br8100 (against the manual annotation ground truth [18]). SpiceMix and SpaGCN(4) were trained on 4 FOVs at the same time, and were evaluated both on single FOVs separately and on 4 FOVs altogether. BayesSpace and SpaGCN(1) were trained on each FOV separately, and were evaluated only on single FOVs. For SpaGCN and BayesSpace, each gray dot represents the score of one of 10 runs with a different random seed and error bars of +/- s.d. across random seeds are shown. **b.** The *in situ* layer assignments of SpiceMix for FOV 151673. The boundaries between ground-truth layers (from [18]) are illustrated by dashed black lines. **c.** The normalized expression of 8 metagenes (from SpiceMix) across the 7 ground-truth layers. Note that metagenes a3 and a6 collectively distinguish layer 4 from layers 3 and 5 (***P*** < 10^−300^; red rectangles). For better visualization, the raw expression levels were divided by the maximum expression level across all spots in the 4 FOVs per metagene. **d.** The rank distribution of known marker genes [47] of 4 cell types in the 8 metagenes. ‘Exc (S)’: markers of excitatory neurons of superficial layers; ‘Exc (D)’: markers of excitatory neurons of deep layers. Metagenes with greater ranks are highlighted by red rectangles for each row (the pairwise ***P*** of Wilcoxon and paired t-test ***P*** are ≤ 10^−7^ for Astro (n=53 genes), ≤ 10^−30^ for Exc (S) (n=406 genes), ≤ 10^−9^ for Exc (D) (n=188 genes), and ≤ 10^−4^ for Oligo (n=67 genes)). **e.** The kernel-smoothed *in situ* expressions of metagenes a4 and a5, demonstrating their differential expressions between the gyric side (the right side, highlighted by arrows in the right panel) and the sulcal side (the upper side, highlighted by arrows in the left panel). **f.** The distribution of the rank difference of gyro-sulcal DEGs between metagenes a4 and a5. DEGs were divided into two groups according to the sign of the logarithm of the fold change. DEGs enriched in the gyric side have greater ranks in metagene a5 than in metagene a4 (Wilcoxon ***P***> and paired t-test ***P*** < 10^−24^), and DEGs enriched in the sulcal side exhibit the opposite trend (Wilcoxon ***P*** and paired t-test ***P*** < 10^−24^). **g.** The *in situ* layer annotations of the ground truth on FOV 151507. **h.** The finer structure annotations of spots (top left) and the *in situ* expressions of metagenes b1-b3 on FOV 151507 (the other three panels). The color legend of the top left panel is in panel **(i)**. Based on the intensity on the histological image, a spot was assigned to a dark stripe (green), a bright gap (blue), or the peripheral region (orange). Spots close to the dark stripe and the bright gap were considered to be a mixture of both structures (grey). As highlighted by black arrows, metagene b1 is enriched in the peripheral region, whereas metagenes b2 and b3 are enriched in the bright gap and the dark stripe, respectively. **i.** The differential expressions of the metagenes b1-b3 across the finer structures. The normalization is the same as in **(c)**. **j.** The *in situ* expression of metagenes b4 and b5 on FOV 151507, implying the delineation of the superficial part (denoted by S) and the deep region (denoted by D) in white matter. **k.** The distribution of the rank difference of the marker genes of Oligo between metagenes b4 and b5.

SpiceMix further refined the Astro-2* and Oligo* clusters of [13] into an astrocyte cluster (Astro-1), two oligodendrocyte clusters (Oligo-1 and Oligo-2), and a cluster that is a mixture of astrocytes and OPCs (Astro-2/OPC), all of which were supported by expression of known marker genes (**Fig.** 4c,e and **Fig.** S12). The oligodendrocyte and OPC types were distinguished by their relative expression of metagenes 12, 13, and 14 from SpiceMix (**Fig.** 4c (right)), which had distinct spatially variable patterns (**Fig.** 4d (bottom)). Metagenes 12 and 13 were highly enriched in layer L6 and exhibited a strong affinity toward each other (**Fig.** 4b,d). Their proportional expression by oligodendrocytes within L6 captured a maturation trajectory from OPCs to Oligo-1 cells that could not be revealed by other methods (see later section). SpiceMix identified metagene 14 that has distinct oligodendrocyte markers (**Fig.** 4c) and scatters from layers L2/3 to L6 (**Fig.** 4d), leading to a spatially distinct Oligo-2 type, clearly separated in the SpiceMix latent space from neighboring excitatory neurons (**Fig.** S13). The expression of oligodendrocyte marker genes from [38] showed that the Oligo-2 cells are late-stage oligodendrocytes, either myelin-forming or fully mature (**Fig.** S12b). In addition, SpiceMix distinguished astrocytes into two types (Astro-1 and Astro-2) based on metagenes 11 and 12. Although the Astro-2 cells shared metagene 12 with OPCs, both their spatial location in the superficial layer and the expression of astrocyte marker genes defined them as astrocytes (**Fig.** S14a). In contrast, the Astro-1 cells expressed metagene 11 with a scattered spatial pattern throughout all layers (**Fig.** 4d). This Astro-1/Astro-2 separation was supported by the expression of known marker genes [42], including *Gfap* (**P**=0.024), a marker gene for astrocytes in the glia limitans, and *Mfge8* (**P**=0.0013), a marker gene for a separate, diffuse astrocyte type (**Fig.** 4e). We found that NMF did not reveal these refined subtypes (**Fig.** S15) and the NMF metagenes typically exhibited unspecific spatial patterns and pairwise affinity (**Fig.** S16, **Fig.** S8d).

These results suggest that SpiceMix is able to refine cell identity and metagene inference with distinct spatial patterns from STARmap data, further demonstrating its unique advantage.

### SpiceMix identifies continuous myelination stages in oligodendrocytes

The expression of metagenes learned by SpiceMix from seqFISH+ and STARmap suggested the existence of continuous factors of cell identity, in this case pertaining to oligodendrocytes, that cannot be described merely by discrete clusters. Since the STARmap dataset included a significant number of oligodendrocyte cells, we performed a more extensive analysis. Applying Monocle2 [43] to the raw expression of cells labelled by SpiceMix as oligodendrocytes showed a clear trajectory from the OPCs of the Astro-2/OPC class to the mature Oligo-1 class (**Fig.** 4f and **Supplementary Methods** A.4.3). The Oligo-2 class is also a mature type of oligodendrocyte (**Fig.** S12), but is likely a distinct type compared to Oligo-1. Importantly, the expression of metagenes 12 and 13, which were highly expressed in OPC and Oligo-1 cells, respectively, strongly correlated with the inferred trajectory (**Fig.** 4f). Along the trajectory, these oligodendrocyte cells exhibited a gradual transition of expression between metagene 12 and 13. Note that Astro-2/OPC (colored magenta in **Fig.** 4) included a distinct group of astrocytes as well, which we removed from consideration (see **Supplementary Results** B.3.1 for details).

To further assess that the differential expression of metagenes 12 and 13 captures the gradually activated myelination process of oligodendrocytes, we used linear regression models to describe the relationship between the expression levels of myelin sheath-related genes and the differences in the proportions of metagenes 12 and 13 in individual cells of the OPC or Oligo-1 cluster. The eleven genes that we tested were those from the STARmap panel attributing to myelin sheath formation, according to Gene Ontology (GO) (**Supplementary Methods** A.4.4 and **Supplementary Results** B.3.1), and were expressed in at least 30% of the cells. We found that the correlations of seven of the eleven genes are significant (*P* < 0.05, after a two-step FDR correction for multiple testing) (**Fig.** 4g and **Fig.** S14b), supporting our hypothesis. One of these genes is *Atp1a2*, recently confirmed by scRNA-seq studies to be suppressed as myelination progresses [44, 45], further demonstrating the robustness of our analysis. Furthermore, we found that the more recent latent variable model scHPF [46], a hierarchical Poisson factorization model for scRNA-seq data, cannot reveal this continuous process of oligodendrocytes, further confirming the importance of spatial information to reveal such processes (**Fig.** S17 and **Supplementary Results** B.3).

This result further demonstrates that the latent representation of SpiceMix is uniquely able to elucidate important biological processes underlying cell states.

### SpiceMix identifies spatial patterns of the human brain from Visium data

We next sought to demonstrate the effectiveness and interpretability of SpiceMix on a dataset of the human dorsolateral prefrontal cortex (DLPFC) acquired by the 10x Genomics Visium platform [18]. We made a direct comparison of SpiceMix to two very recent methods on this dataset: SpaGCN [23] and BayesSpace [25], which was designed for Visium data. We also analyzed the learned spatial metagenes from SpiceMix to demonstrate its unique capability.

SpiceMix achieved consistent advantages over SpaGCN and BayesSpace in identifying the layer structures of DLPFC, which consisted of six cortical layers (layer L1 to layer L6) and white matter. We used the manual annotations provided by the original study [18] as the ground truth and evaluated the performance of different methods using the ARI scores (**Fig.** 5a). For SpiceMix, here the continuous latent states of spots were clustered by Louvain clustering for subsequent analysis (**Supplementary Methods** A.5.3). Since the FOVs in samples Br5595 and Br5292 either contain ambiguous spots or additional anatomic structures (**Supplementary Methods** A.5.2), we focused on the 4 FOVs from sample Br8100 for this analysis. The clusters from SpiceMix produced an ARI score between 0.54 and 0.61 (average 0.575) with consistent advantage over SpaGCN and BayesSpace for each FOV (**Fig.** 5a). We found that SpiceMix was able to learn spatially variable metagenes that clearly manifest the layer structure of DLPFC (**Fig.** S20) (see **Supplementary Table**). We observed that although SpaGCN and BayesSpace were able to produce a layer-like patterns, they typically did not closely match the boundaries of the ground truth annotations (**Fig.** S18 and S19). In contrast, SpiceMix produced contiguous layers for all FOVs and identified clearer boundaries. Across all FOVs, layer L4 could not be reliably identified by any method. However, the combinatorial expression of the metagenes learned by SpiceMix, in particular metagenes a3 and a6, showed differential expression among L3, L4, and L5 (*P* < 10^−300^, highlighted in **Fig.** 5c). Using all four FOVs as input did not significantly affect the ARI score from SpaGCN (**Fig.** 5a), and we were unable to run BayesSpace effectively on all four FOVs simultaneously. Overall, SpiceMix showed clear advantage for the identification of tissue layer structures of DLPFC based on Visium data.

The interpretability of metagenes from SpiceMix helped unveil spatially variable expression and spatial patterns of important cell types of DLPFC. Here, we used differentially expressed genes (DEGs) identified from [47] (**Supplementary Methods** A.5.4). We found that the marker genes of astrocytes had higher ranks in metagene a1 than any other metagene (**Fig.** 5d), and metagene a1 was spatially present in all 7 layers at similar levels (**Fig.** 5c), consistent with the spatial distribution of astrocytes observed in a very recent work [48]. The marker genes of oligodendrocytes were enriched in metagenes a6 and a7 (**Fig.** 5d), and both metagenes were enriched in deep layers or the white matter (**Fig.** 5c), consistent with the spatial distributions of oligodendrocytes [49]. Notably, metagene a7 was nearly restricted to the white matter, whereas metagene a6 was enriched in deep cortical layers (**Fig.** 5c), suggesting the potential separation of cortical oligodendrocytes and white matter oligodendrocytes. Moreover, the marker genes of excitatory neurons located in superficial layers and deep layers were enriched with metagenes a3 and a6, respectively, and metagenes a3 and a6 were present mostly in layers L1-L3 and layer L6, accordingly, consistent with the layer-like spatial patterns of excitatory neural types (**Fig.** 5c-d). These findings confirm the unique ability of SpiceMix to unveil spatially variable features and cell type composition.

### SpiceMix delineates finer anatomic structures of the human brain from Visium data

We then aimed to further demonstrate the ability of SpiceMix for identifying detailed spatial variable metagenes and cell composition using the DLPFC Visium data [18] by comparing to finer anatomic structures of the brain. Specifically, on the four FOVs from sample Br8100, the metagenes produced by SpiceMix captured the gyro-sulcal variability (**Fig.** 5e-f and **Fig.** S21). We found that more than 50% of the genes used for SpiceMix were differentially expressed across the two regions (**Supplementary Methods** A.5.4), strongly supporting this separation. Metagenes a4 and a5 captured the gradual transition between these two regions (**Fig.** 5e), and the relative ranking of DEGs within each metagene, according to the gene’s weight in the metagene, was significantly associated with the respective region (*P* < 10^−24^) (**Fig.** 5f). This demonstrates the distinct ability of SpiceMix to represent gradual changes in spatial expression by metagenes.

Next, we sought to show that SpiceMix can identify subtle anatomical structures from spatial transcriptome data alone. As a proof-of-principle, we applied SpiceMix on FOV 151507 from sample Br5292 (**Fig.** 5g) and discovered spatially variable metagenes (see **Supplementary Table** for the list). We found that metagenes b1-b3 defined 3 finer anatomical structures within layer L1 annotated in [18] (**Fig.** 5h-i). According to the brightness of the staining in the histology image, we classified each spot into one of four types (**Supplementary Methods** A.5.5 and **Fig.** 5h (top left)): the dark stripe (yellow), the bright gap (green), the flanking cortex (blue), and ambiguous mixtures of these three regions (grey). All 7 marker genes of mural cells, which constitute the wall of blood vessels, from [42] that passed quality control (**Supplementary Methods** A.5.5) were expressed at higher levels in the dark stripe, and the enrichment of 5 out of the 7 genes was significant (***P***≤ 0.002), suggesting that the dark stripe is potentially a blood vessel. Aside from the brightness, spots had other varying phenotypes across the three regions, such as cell density, UMI count, and mitochondrial RNA ratio (**Fig.** S22a), indicating that these three regions are biologically different. We found that metagenes b1, b2, and b3 were enriched in the flanking cortex, the white gap, and the blood vessel, respectively (***P***≤ 10^−10^) (**Fig.** 5h-i), supporting the delineation of the three anatomical structures by SpiceMix.

Another example of interpretable delineation of finer structures was that metagenes b4 and b5 defined two finer anatomical structures in the white matter region (**Fig.** 5j). Specifically, metagene b4 was mainly present in a 400*μ*m-wide superficial layer (**Fig.** 5j (S)), whereas metagene b5 was nearly restricted to the deep part (**Fig.** 5j (D)). Spots also exhibited different phenotypes, such as cell density and UMI count, across the two finer structures (**Fig.** S22b). We identified 751 DEGs between the two structures (**Supplementary Methods** A.5.5) and found that 62% of marker genes of oligodendrocytes were enriched in the deep part, whereas only one marker gene was enriched in the superficial layer (**Fig.** S22c), supporting that the two structures are two biologically meaningful anatomical structures rather than artifacts. Consistent with this finding, marker genes of oligodendrocytes had a higher rank in metagene b2, which was enriched in the deep part (**Fig.** 5k).

Together, these results further demonstrated the ability of SpiceMix to capture subtle but biologically important anatomical structures from spatial transcriptome data alone.

## Discussion

In this work, we developed SpiceMix, an unsupervised method for modeling the diverse factors that collectively contribute to cell identity in complex tissues based on various types of spatial transcriptome data. The novel integration of NMF and HMRF in the underlying model of SpiceMix combines the expressive power of NMF for modeling gene expression with the HMRF for modeling spatial relationships, advancing current state-of-the-art modeling for spatial transcriptomics. We evaluated the performance of SpiceMix on simulated data that approximates the mouse cortex spatial transcriptome, showing a clear advantage over existing approaches. The application of SpiceMix to single-cell spatial transcriptome data of the mouse primary visual cortex from seqFISH+ and STARmap demonstrated its effectiveness in producing reliable spatially variable metagenes and and informative latent representations of cell identity, which yielded more accurate and finer cell type identification than prior approaches and uncovered important biological processes underlying cell states. When applied to the human DLPFC data acquired by Visium, SpiceMix improved the identification of annotated layers compared to existing methods and revealed finer anatomical structures.

As future work, SpiceMix could be further enhanced by incorporating additional modalities such as scRNA-seq data. Other recent computational methods have been used to study scRNA-seq and spatial transcriptomic data jointly [30, 31, 33, 50], but unlike SpiceMix, they do not attempt to comprehensively model spatial cell-to-cell relationships. With proper normalization and preprocessing, data from scRNA-seq could easily be integrated into the SpiceMix framework. This additional data may improve the inference of the latent variables and parameters of the model, which could further improve the modeling of cellular heterogeneity. In addition, further enhancements could be made to the probabilistic model of SpiceMix, such as including additional priors, to tailor toward particular application contexts.

A significant feature of SpiceMix for cellular identity discovery is the spatially variable metagene formulation. These metagenes are able to model the interplay of spatial and intrinsic composition of the transcriptome and not merely the spatial patterns of individual genes [26, 29]. Crucially, as part of the model formulation, SpiceMix considers how these spatially variable metagenes are integrally related to continuous states of cell identity, which represents a major distinction compared to other approaches [22, 23]. We have shown in several contexts how metagenes can reveal spatially important patterns, such as continuous maturation stages of oligodendrocytes and genes associated with anatomical structures like the gyrus and sulcus regions. The precision with which SpiceMix can identify metagenes and separate the interplay of different factors is tied to the scale of the dataset. Increasing the scale of spatial transcriptomic studies will potentially enable SpiceMix to discover even finer resolution patterns. In particular, adding the new ability to explicitly account for temporal variation of spatially variable metagenes over time will further enhance the applicability of SpiceMix.

As the area of spatial transcriptomics continues to thrive and data become more widely available, SpiceMix will be a uniquely useful tool to facilitate new discoveries. In particular, refined cell identity with SpiceMix has the potential to improve future studies of cell-cell interactions [51]. Additionally, SpiceMix is not limited to transcriptomic data, and its methodology may also be well-suited for other recent multi-omic spatial data, e.g., DBiT-seq [52]. As more datasets become available, SpiceMix could be integrated with pipeines for managing spatial transcriptomic data, such as Squidpy [53], to streamline analysis. Overall, SpiceMix is a powerful framework that can serve as an essential tool for the analysis of diverse types of spatial transcriptome and multi-omic data, with the distinct advantage that it can unravel the complex mixing of latent intrinsic and spatial factors of heterogeneous cell identity in complex tissues.

## Methods

### The new probabilistic graphical model NMF-HMRF in SpiceMix

#### Gene expression as matrix factorization

We consider the expression of individual cells 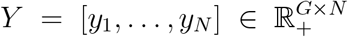, where constants *G* and *N* denote the number of genes and cells, respectively, to be the product of *K* underlying factors (i.e., metagenes), 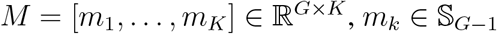, and weights, 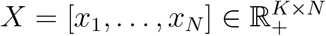, i.e.,

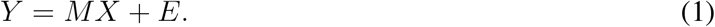

This follows the non-negative matrix factorization (NMF) formulation of expression of prior work [54]. The term 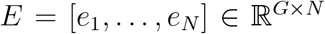 captures unexplained variation or noise, which we model as i.i.d. Gaussian, i.e., 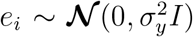. To resolve the scaling ambiguity between *M* and *X*, we constrain the columns of *M* to sum to one, so as to lie in the (*G* – 1)-dimensional simplex, 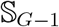. For notational consistency, we use capital letters to denote matrices and use lowercase letters denote their column vectors.

#### Graphical model formulation

Our formulation for the new probabilistic graphical model NMF-HMRF in SpiceMix enhances standard NMF by modeling the spatial correlations among samples (i.e., cells or spots in this context) via the HMRF [55]. This novel integration aids inference of the latent *M* and *X* by enforcing spatial consistency. The spatial relationship between cells in tissue is represented as a graph 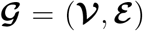 of nodes 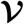 and edges 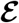, where each cell is a node and edges are determined from the spatial locations. Any graph construction algorithm, such as distance thresholding or Delaunay triangulation, can be used for determining edges. For each node *i* in the graph, the measured gene expression vector, *y_i_*, is the set of observed variables and the weights, *x_i_*, describing the mixture of metagenes are the hidden states. The observations are related to the hidden variables via the potential function *ϕ*, which captures the NMF formulation. The spatial affinity between the metagene proportions of neighboring cells is captured by the potential function *φ*. Together, these elements constitute the HMRF.

More specifically, the potential function *ϕ* measures the squared reconstruction error of the observed expression of cell *i* according to the estimated *x_i_* and *M*,

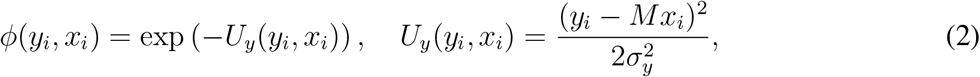

where 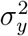 represents the variation of expression, or noise, of the NMF. The spatial potential function *φ* measures the inner-product between the metagene proportions of neighboring cells *i* and *j*, weighted by the learned, pairwise correlation matrix 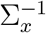, which captures the spatial affinity of metagenes, i.e.,

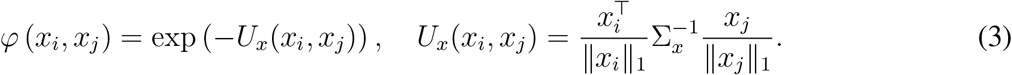

This form for *φ* has several motivations. The weighted inner-product allows the affinity between two cells to be decomposed simply as the weighted sum of affinities between metagenes and for the metagenes to have different and learnable affinities between each other. It also allows the model to capture both positive and negative affinities between metagenes. By normalizing the weights *x_i_* of each cell, any scaling effects, such as cell size, are removed. In this way, the similarity that is measured is purely a function of the relative proportions of metagenes. This form also affords a straightforward interpretation for the affinity matrix 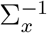. Lastly, it is more convenient for optimization.

Given an observed dataset, the model can be learned by maximizing the likelihood of the data. By the Hammersley-Clifford theorem [56], the likelihood of the data for the pairwise HMRF can be formulated as the product of pairwise dependencies between nodes,

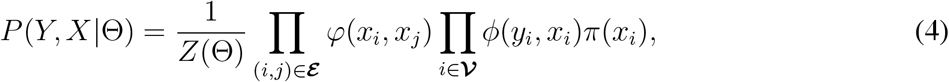

where Θ = {Δ, *M*} is the set of model parameters and metagenes and *Z*(Θ) is the normalizing partition function that ensures *P* is a proper probability distribution. The potential function *π* is added to capture an exponential prior on the hidden states *X*,

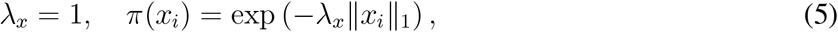

with scale parameter 1. We normalize the average of the total normalized expression levels in individual cells to *K* correspondingly.

#### Parameter priors

We introduce a regularization hyperparameter *λ*_∑_ on the spatial affinities, which allows the users to control the importance of the spatial relationships during inference to suit the sample of interest. As the parameter decreases, the influence of spatial affinities during inference diminishes and the model becomes more similar to standard NMF. If we represent *λ*_∑_ in the form 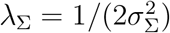, we can treat it as a Gaussian prior, with zero mean and 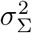 variance, on the elements of the spatial affinity matrix 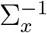,

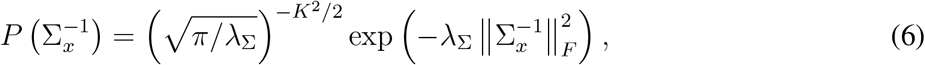

where *F* denotes the Frobenius norm. Note that the matrix 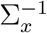 is forced to be transpose symmetric.

### Alternating estimation of hidden states and model parameters

To infer the hidden states and model parameters of the NMF-HMRF model in SpiceMix, we optimize the data likelihood via coordinate ascent, alternating between optimizing hidden states and model parameters. This new optimization scheme is summarized in Algorithm 1. First, to make inference tractable, we approximate the joint probability of the hidden states by the pseudo-likelihood [56], which is the product of conditional probabilities of the hidden state of individual nodes given that of their neighbors,

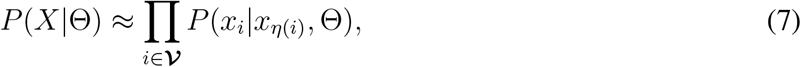

where *η*(*i*) is the set of neighbors to node *i*.

#### Algorithm 1 NMF-HMRF model-fitting and hidden state estimation.

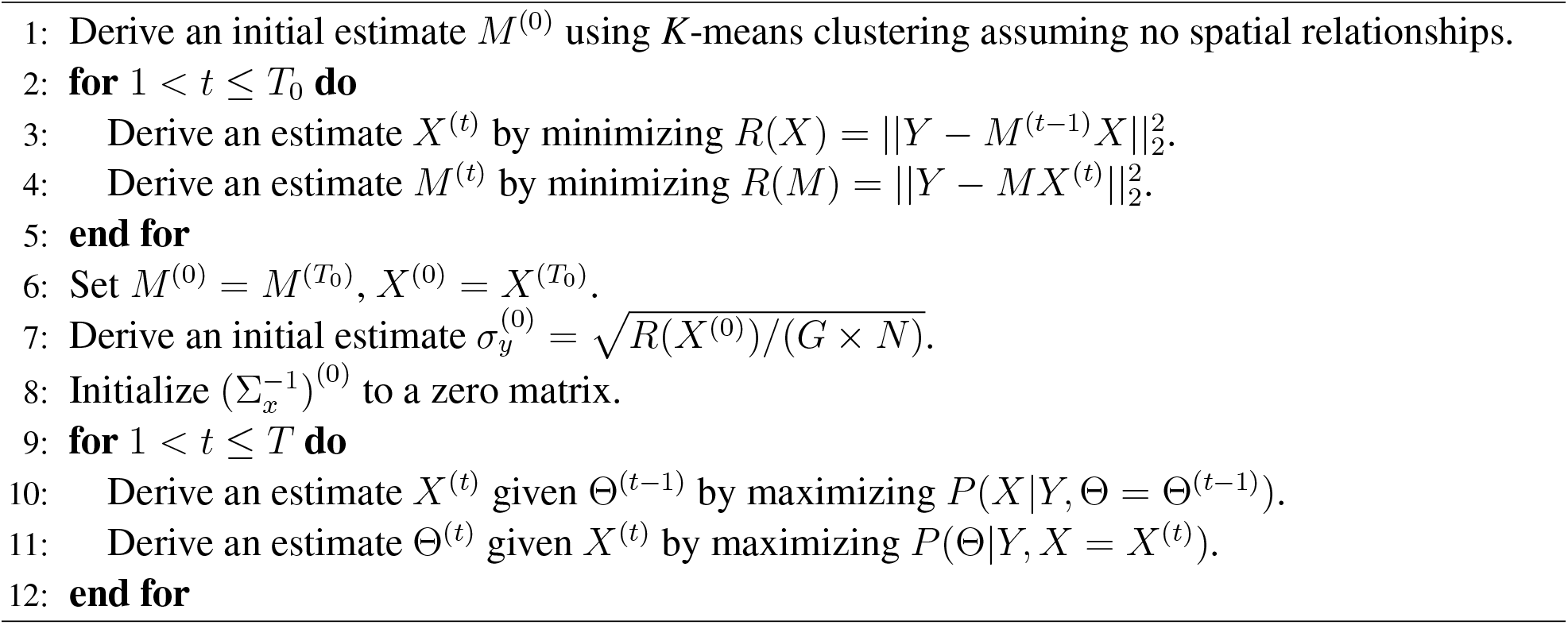

#### Estimation of hidden states

Given parameters Θ of the model, we estimate the factorizations *X* by maximizing their posterior distribution. The maximum a posteriori (MAP) estimate of *X* is given by:

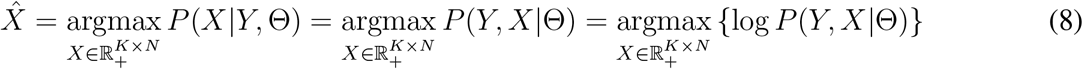

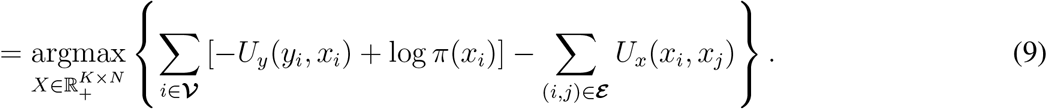

This is a quadratic program and can be solved efficiently via the iterated conditional model (ICM) [57] using the software package Gurobi [58] (see **Supplementary Methods** A.1.1 for more details of the optimization for hidden states).

#### Estimation of model parameters

Given an estimate of the hidden states *X*, we can likewise solve for the unknown model parameters Θ by maximizing their posterior distribution. The MAP estimate of the parameters Θ is given by:

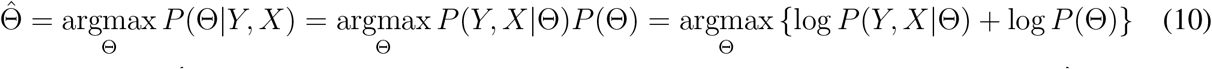

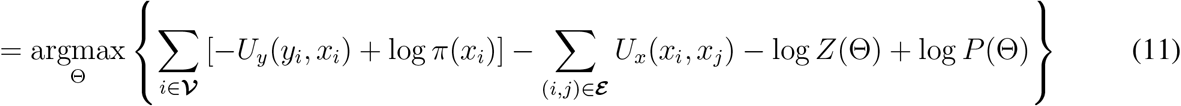

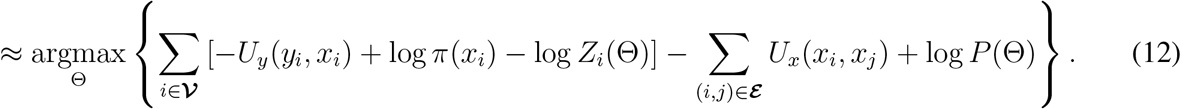

Eqn. 12 is an approximation by the mean-field assumption [56], which is used, in addition to the pseudolikelihood assumption, to make the inference of model parameters tractable. We note that we can estimate metagenes, spatial affinity, and the noise level independently. The MAp estimate of the metagenes *M* is a quadratic program, which is efficient to solve. The MAP estimate of 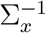 is convex and is solved by the optimizer Adam [59]. Due to the complexity of the partition function *Z_i_*(Θ) of the likelihood, which includes integration over X, it is approximated by Taylor’s expansion. Since it is a function of Θ, this computation must be performed at each optimization iteration. See **Supplementary Methods** A.1.2 for details of the optimization method for model parameters.

#### Initialization

To produce the initial estimates of the model parameters and hidden states, we do the following. First, we use a common strategy for initializing NMF, which is to cluster the data using *K*-means clustering, with *K* equal to the number of metagenes, and use the means of the clusters as an estimate of the metagenes. We then alternate for *T*_0_ iterations between solving the NMF objective for *X* and *M*. This produces, in only a few quick iterations, an appropriate initial estimate for the algorithm, which will be subsequently refined. We observed that if *T*_0_ is too large, it can cause the algorithm to prematurely reach a local minimum before spatial relationships are considered. However, this value can be easily tuned by experimentation, and in our analysis, we found that just 5 iterations were necessary.

### Empirical running time

On a machine with eight 3.6 GHz CPUs and one GeForce 1080 Ti GPU, SpiceMix takes 0.5-2 hours to run on a typical spatial transcriptome dataset with 2,000 genes and 1,000 cells. The GPU is used for the first 5 iterations, or around that number, only, when the spatial affinity matrix 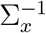 is changed significantly. Later on, most time is spent solving quadratic programs. Since the algorithm uses a few iterations of NMF to provide an initial estimate, which is a reasonable starting point, it is expected to find a good initial estimate of metagenes and latent states efficiently.

### Generation and analysis of simulated data

We generated simulated spatial transcriptomic data following expression and spatial patterns of cells of the mouse primary visual cortex. Cells in the mouse cortex are classified into three primary categories: inhibitory neurons, excitatory neurons, and non-neurons or glial cells [36, 60]. Excitatory neurons in the cortex exhibit dense, concentrated, layer-wise specificity, whereas inhibitory neurons are sparse and can be spread across several layers. Non-neuronal cells can be either layer-specific or dispersed across layers. We simulated data from an imaging-based method applied to a slice of tissue, which consists of four distinct vertical layers and eight cell types: four excitatory, two inhibitory, and two glial (**Fig.** 2a). Each layer was densely populated by one layer-specific excitatory neuron type. The two inhibitory neuron types were scattered sparsely throughout several layers. One non-neuronal type was restricted to the first layer and the other was scattered sparsely throughout several layers. For each simulated image, or tissue sample, 500 cells were created with locations generated randomly in such a way so as to maintain a minimum distance between any two cells, so that the density of cells across the sample was roughly constant. With this spatial layout of cells, we devised two methodologies for generating gene expression data for individual cells. The first uses a metagene-based formulation (see **Supplementary Methods** A.2.1) and the second uses a recent method, scDesign2 [35], which we fit to real scRNA-seq data of the mouse cortex [36] (see **Supplementary Methods** A.2.2). See **Supplementary Methods** A.2.3 for details of the methodology of the analysis of the two simulation datasets.

### Data processing for the spatial transcriptome data used in this work

We applied SpiceMix on a seqFISH+ dataset that profiled the mouse primary visual cortex [12]. For details of the methodology of our analysis, see **Supplementary Methods** A.3. Specifically, for details of data preprocessing, see **Supplementary Methods** A.3.1. For details of the selection of hyperparameters for each algorithm, see **Supplementary Methods** A.3.2. For details of our selection of Louvain clusters from [12] used in our comparative analysis with SpiceMix, see **Supplementary Methods** A.3.3. For additional details of the method to justify the excitatory neuron clusters of SpiceMix, see **Supplementary Methods** A.3.4.

We also applied SpiceMix on a STARmap dataset that profiled the mouse primary visual cortex [13]. For details of the methodology of our analysis, see **Supplementary Methods** A.4. Specifically, for details of data preprocessing, see **Supplementary Methods** A.4.1. For details of the selection of hyperparameters for each algorithm, see **Supplementary Methods** A.4.2. For details of our trajectory analysis of oligodendrocytes using Monocle2, see **Supplementary Methods** A.4.3. For additional details of GO enrichment analysis of myelin sheath formation in oligodendrocytes, see **Supplementary Methods** A.4.4.

Lastly, we applied SpiceMix to a dataset acquired from the 10x Genomics Visium platform that profiled spatial transcriptome of the human DLPFC [18]. Similar to our procedure for other platforms, we applied a preprocessing pipeline; see **Supplementary Methods** A.5.1 for details. For details on the selection of the four FOVs from the Br8100 sample to use for the ARI score comparison, see **Supplementary Methods** A.5.2. For details on the ARI score comparison between SpiceMix, SpaGCN, and BayesSpace on these FOVs, see **Supplementary Methods** A.5.3. For details of the subsequent analysis of SpiceMix metagenes on these FOVs, see **Supplementary Methods** A.5.4. For details of the analysis of SpiceMix metagenes on sample Br5292, see **Supplementary Methods** A.5.5.

Doublet detection [61] was performed on the seqFISH+ and STARmap datasets to confirm that none of the cells in either dataset were doublets; see **Supplementary Methods** A.6.1 for details. For the explanation of our method for constructing the cell-type affinity matrix for the seqFISH+ and STARmap datasets, see **Supplementary Methods** A.6.2.

## Code Availability

The source code of SpiceMix can be accessed at: https://github.com/ma-compbio/SpiceMix.

## Acknowledgements

This work was supported in part by the National Institutes of Health Common Fund 4D Nucleome Program grant UM1HG011593 (J.M.), National Institutes of Health Common Fund Cellular Senescence Network Program grant UG3CA268202 (J.M.), National Institutes of Health grant R01HG007352 (J.M.), and National Science Foundation grant 1717205 (J.M.). J.M. is additionally supported by a Guggenheim Fellowship from the John Simon Guggenheim Memorial Foundation.

## Author Contributions

Conceptualization, B.C. and J.M.; Methodology, B.C., T.Z., and J.M.; Software, T.Z. and B.C.; Investigation, B.C., T.Z., S.A., and J.M.; Writing – Original Draft, B.C., T.Z., and J.M.; Writing – Review & Editing, B.C., T.Z., S.A., and J.M.; Funding Acquisition, J.M.

## References

[1] Arendt D, Musser JM, Baker CV, Bergman A, Cepko C, Erwin DH, et al. The origin and evolution of cell types. Nature Reviews Genetics. 2016;17(12):744.

[2] Chen X, Teichmann SA, Meyer KB. From Tissues to Cell Types and Back: Single-Cell Gene Expression Analysis of Tissue Architecture. Annual Review of Biomedical Data Science. 2018;1:29–51.

[3] Consortium H, et al. The human body at cellular resolution: the NIH Human Biomolecular Atlas Program. Nature. 2019;574(7777):187.

[4] Trapnell C. Defining cell types and states with single-cell genomics. Genome Research. 2015;25(10):1491–1498.

[5] Wagner A, Regev A, Yosef N. Revealing the vectors of cellular identity with single-cell genomics. Nature Biotechnology. 2016;34(11):1145.

[6] Stuart T, Satija R. Integrative single-cell analysis. Nature Reviews Genetics. 2019;20(5):257–272.

[7] Lee JH, Daugharthy ER, Scheiman J, Kalhor R, Yang JL, Ferrante TC, et al. Highly multiplexed subcellular RNA sequencing in situ. Science. 2014;343(6177):1360–1363.

[8] Chen KH, Boettiger AN, Moffitt JR, Wang S, Zhuang X. Spatially resolved, highly multiplexed RNA profiling in single cells. Science. 2015;348(6233):aaa6090.

[9] Shah S, Lubeck E, Zhou W, Cai L. In situ transcription profiling of single cells reveals spatial organization of cells in the mouse hippocampus. Neuron. 2016;92(2):342–357.

[10] Ståhl PL, Salmén F, Vickovic S, Lundmark A, Navarro JF, Magnusson J, et al. Visualization and analysis of gene expression in tissue sections by spatial transcriptomics. Science. 2016;353(6294):78–82.

[11] Moffitt JR, Bambah-Mukku D, Eichhorn SW, Vaughn E, Shekhar K, Perez JD, et al. Molecular, spatial, and functional single-cell profiling of the hypothalamic preoptic region. Science. 2018;362(6416):eaau5324.

[12] Eng CHL, Lawson M, Zhu Q, Dries R, Koulena N, Takei Y, et al. Transcriptome-scale superresolved imaging in tissues by RNA seqFISH+. Nature. 2019;568:235–239.

[13] Wang X, Allen WE, Wright MA, Sylwestrak EL, Samusik N, Vesuna S, et al. Three-dimensional intact-tissue sequencing of single-cell transcriptional states. Science. 2018;341(6400):eaat5691.

[14] Rodriques SG, Stickels RR, Goeva A, Martin CA, Murray E, Vanderburg CR, et al. Slide-seq: A scalable technology for measuring genome-wide expression at high spatial resolution. Science. 2019;363(6434):1463–1467.

[15] Vickovic S, Eraslan G, Salmén F, Klughammer J, Stenbeck L, Schapiro D, et al. High-definition spatial transcriptomics for in situ tissue profiling. Nature Methods. 2019;16(10):987–990.

[16] Zhuang X. Spatially resolved single-cell genomics and transcriptomics by imaging. Nature Methods. 2021;18(1):18–22.

[17] Larsson L, Frisén J, Lundeberg J. Spatially resolved transcriptomics adds a new dimension to genomics. Nature Methods. 2021;18(1):15–18.

[18] Maynard KR, Collado-Torres L, Weber LM, Uytingco C, Barry BK, Williams SR, et al. Transcriptome-scale spatial gene expression in the human dorsolateral prefrontal cortex. Nature Neuroscience. 2021 Mar;24(3):425–436.

[19] Lein E, Borm LE, Linnarsson S. The promise of spatial transcriptomics for neuroscience in the era of molecular cell typing. Science. 2017;358(6359):64–69.

[20] Palla G, Fischer DS, Regev A, Theis FJ. Spatial components of molecular tissue biology. Nature Biotechnology. 2022:1–11.

[21] Schapiro D, Jackson HW, Raghuraman S, Fischer JR, Zanotelli VR, Schulz D, et al. histoCAT: analysis of cell phenotypes and interactions in multiplex image cytometry data. Nature Methods. 2017;14(9):873.

[22] Zhu Q, Shah S, Dries R, Cai L, Yuan GC. Identification of spatially associated subpopulations by combining scRNAseq and sequential fluorescence in situ hybridization data. Nature Biotechnology. 2018;36(12):1183.

[23] Hu J, Li X, Coleman K, Schroeder A, Ma N, Irwin DJ, et al. SpaGCN: Integrating gene expression, spatial location and histology to identify spatial domains and spatially variable genes by graph convolutional network. Nature Methods. 2021;18(11):1342–1351.

[24] Jerby L, Regev A. Mapping multicellular programs from single-cell profiles. bioRxiv. 2020.

[25] Zhao E, Stone MR, Ren X, Guenthoer J, Smythe KS, Pulliam T, et al. Spatial transcriptomics at subspot resolution with BayesSpace. Nature Biotechnology. 2021;39(11):1375–1384.

[26] Svensson V, Teichmann SA, Stegle O. SpatialDE: identification of spatially variable genes. Nature Methods. 2018;15(5):343.

[27] Arnol D, Schapiro D, Bodenmiller B, Saez-Rodriguez J, Stegle O. Modeling Cell-Cell Interactions from Spatial Molecular Data with Spatial Variance Component Analysis. Cell Reports. 2019;29(1):202–211.

[28] Nitzan M, Karaiskos N, Friedman N, Rajewsky N. Gene expression cartography. Nature. 2019;576(7785):132–137.

[29] Sun S, Zhu J, Zhou X. Statistical analysis of spatial expression patterns for spatially resolved transcriptomic studies. Nature Methods. 2020;17(2):193–200.

[30] Stuart T, Butler A, Hoffman P, Hafemeister C, Papalexi E, Mauck III WM, et al. Comprehensive Integration of Single-Cell Data. Cell. 2019;177(7):1888–1902.e21.

[31] Welch JD, Kozareva V, Ferreira A, Vanderburg C, Martin C, Macosko EZ. Single-Cell Multi-omic Integration Compares and Contrasts Features of Brain Cell Identity. Cell. 2019;177(7):1873–1887.e17.

[32] Elosua-Bayes M, Nieto P, Mereu E, Gut I, Heyn H. SPOTlight: seeded NMF regression to de-convolute spatial transcriptomics spots with single-cell transcriptomes. Nucleic Acids Research. 2021;49(9):e50–e50.

[33] Biancalani T, Scalia G, Buffoni L, Avasthi R, Lu Z, Sanger A, et al. Deep learning and alignment of spatially resolved single-cell transcriptomes with Tangram. Nature Methods. 2021;18(11):1352–1362.

[34] Lee DD, Seung HS. Algorithms for non-negative matrix factorization. In: Advances in Neural Information Processing Systems; 2001. p. 556–562.

[35] Sun T, Song D, Li WV, Li JJ. scDesign2: a transparent simulator that generates high-fidelity singlecell gene expression count data with gene correlations captured. Genome Biology. 2021;22(1):1–37.

[36] Tasic B, Menon V, Nguyen TN, Kim TK, Jarsky T, Yao Z, et al. Adult mouse cortical cell taxonomy revealed by single cell transcriptomics. Nature Neuroscience. 2016;19(2):335.

[37] Satija R, Farrell JA, Gennert D, Schier AF, Regev A. Spatial reconstruction of single-cell gene expression data. Nature Biotechnology. 2015;33(5):495.

[38] Marques S, Zeisel A, Codeluppi S, van Bruggen D, Falcão AM, Xiao L, et al. Oligodendrocyte heterogeneity in the mouse juvenile and adult central nervous system. Science. 2016;352(6291):1326–1329.

[39] Zhao C, Deng Y, Liu L, Yu K, Zhang L, Wang H, et al. Dual regulatory switch through interactions of Tcf7l2/Tcf4 with stage-specific partners propels oligodendroglial maturation. Nature Communications. 2016;7(1):1–15.

[40] Linington C, Bradl M, Lassmann H, Brunner C, Vass K. Augmentation of demyelination in rat acute allergic encephalomyelitis by circulating mouse monoclonal antibodies directed against a myelin/oligodendrocyte glycoprotein. The American Journal of Pathology. 1988;130(3):443.

[41] Tasic B, Yao Z, Graybuck LT, Smith KA, Nguyen TN, Bertagnolli D, et al. Shared and distinct transcriptomic cell types across neocortical areas. Nature. 2018;563(7729):72–78.

[42] Zeisel A, Muñoz-Manchado AB, Codeluppi S, Lönnerberg P, La Manno G, Juréus A, et al. Cell types in the mouse cortex and hippocampus revealed by single-cell RNA-seq. Science. 2015;347(6226):1138–1142.

[43] Qiu X, Mao Q, Tang Y, Wang L, Chawla R, Pliner HA, et al. Reversed graph embedding resolves complex single-cell trajectories. Nature Methods. 2017;14(10):979–982.

[44] Marques S, van Bruggen D, Vanichkina DP, Floriddia EM, Munguba H, Väremo L, et al. Transcriptional convergence of oligodendrocyte lineage progenitors during development. Developmental Cell. 2018;46(4):504–517.

[45] Beiter RM, Fernández-Castañeda A, Rivet-Noor C, Merchak A, Bai R, Slogar E, et al. Evidence for oligodendrocyte progenitor cell heterogeneity in the adult mouse brain. bioRxiv. 2020.

[46] Levitin HM, Yuan J, Cheng YL, Ruiz FJ, Bush EC, Bruce JN, et al. De novo gene signature identification from single-cell RNA-seq with hierarchical Poisson factorization. Molecular Systems Biology. 2019;15(2):e8557.

[47] Dataset: Allen Institute for Brain Science (2021). Allen Cell Types Database – Human Multiple Cortical Areas [dataset]. Available from: http://celltypes.brain-map.org/rnaseq;.

[48] Zhang M, Eichhorn SW, Zingg B, Yao Z, Cotter K, Zeng H, et al. Spatially resolved cell atlas of the mouse primary motor cortex by MERFISH. Nature. 2021;598(7879):137–143.

[49] Tan SS, Kalloniatis M, Truong HT, Binder MD, Cate HS, Kilpatrick TJ, et al. Oligodendrocyte positioning in cerebral cortex is independent of projection neuron layering. Glia. 2009;57(9):1024–1030.

[50] Korsunsky I, Millard N, Fan J, Slowikowski K, Zhang F, Wei K, et al. Fast, sensitive and accurate integration of single-cell data with Harmony. Nature Methods. 2019;16(12):1289–1296.

[51] Armingol E, Officer A, Harismendy O, Lewis NE. Deciphering cell–cell interactions and communication from gene expression. Nature Reviews Genetics. 2021;22(2):71–88.

[52] Liu Y, Yang M, Deng Y, Su G, Enninful A, Guo CC, et al. High-spatial-resolution multi-omics sequencing via deterministic barcoding in tissue. Cell. 2020;183(6):1665–1681.

[53] Palla G, Spitzer H, Klein M, Fischer D, Schaar AC, Kuemmerle LB, et al. Squidpy: a scalable framework for spatial omics analysis. Nature Methods. 2022;19(2):171–178.

[54] Brunet JP, Tamayo P, Golub TR, Mesirov JP. Metagenes and molecular pattern discovery using matrix factorization. Proceedings of the National Academy of Sciences. 2004;101(12):4164–4169.

[55] Zhang Y, Brady M, Smith S. Segmentation of brain MR images through a hidden Markov random field model and the expectation-maximization algorithm. IEEE Transactions on Medical Imaging. 2001;20(1):45–57.

[56] Murphy K. Machine learning: a probabilistic perspective. MIT Press; 2012.

[57] Besag J. On the statistical analysis of dirty pictures. Journal of the Royal Statistical Society: Series B (Methodological). 1986;48(3):259–279.

[58] Gurobi Optimization L. Gurobi Optimizer Reference Manual; 2020. Available from: http://www.gurobi.com.

[59] Kingma DP, Ba J. Adam: A method for stochastic optimization. arXiv preprint arXiv:14126980. 2014.

[60] Lein ES, Hawrylycz MJ, Ao N, Ayres M, Bensinger A, Bernard A, et al. Genome-wide atlas of gene expression in the adult mouse brain. Nature. 2007;445(7124):168–176.

[61] Gayoso A. Shor J. JonathanShor/DoubletDetection: Doubletdetection V3. 2018.

