## Supplemental Information for "SPICEMIX: Integrative single-cell spatial modeling of cell identity"

#### A Supplementary Methods

##### A.1 Additional method details of the algorithm

###### *A.1.1 Derivation of optimization of hidden states*

We rewrite and expand Eqn. 9,

$$\hat{X}_{\mathrm{MAP}} = \underset{X \in R_+^{K \times N}}{\operatorname{argmax}} \{P(X \mid Y, \Theta)\} = \underset{X \in R_+^{K \times N}}{\operatorname{argmax}} \left\{ \prod_{i \in \boldsymbol{\mathcal{V}}} \pi(x_i)\phi(y_i, x_i) \prod_{(i,j) \in \boldsymbol{\mathcal{E}}} \varphi(x_i, x_j) \right\} \tag{13}$$

$$= \underset{X \in R_+^{K \times N}}{\operatorname{argmax}} \left\{ \sum_{i \in \boldsymbol{\mathcal{V}}} [\ln \pi(x_i) + \ln \phi(y_i, x_i)] + \sum_{(i,j) \in \boldsymbol{\mathcal{E}}} \ln \varphi(x_i, x_j) \right\} \tag{14}$$

$$= \underset{X \in R_+^{K \times N}}{\operatorname{argmin}} \left\{ \sum_{i \in \boldsymbol{\mathcal{V}}} \left[ \frac{(y_i - M x_i)^2}{2\sigma_y^2} + \lambda_x^\top x_i \right] + \sum_{(i,j) \in \boldsymbol{\mathcal{E}}} \frac{x_i^\top}{\|x_i\|_1} \Sigma_x^{-1} \frac{x_j}{\|x_j\|_1} \right\}. \tag{15}$$

In the last equation, we substitute $\pi$, $\varphi$, and $\phi$ with their definition in Eqn. 2, 3, and 5.

For each $i$, we decompose $x_i$ into $s_i z_i$, where $s_i \in \mathbb{R}_+$ is a non-negative real number, representing the $L_1$ norm of $x_i$, and $z_i \in \mathbb{S}_{K-1}$ is a point on the $K$-dimensional simplex:

$$x_i = s_i z_i \quad \text{s.t.} \quad s_i \in \mathbb{R}_+, z_i \in \mathbb{S}_{K-1}. \tag{16}$$

Then the programming becomes:

$$\underset{\forall i,\, s_i \in \mathbb{R}_+, z_i \in \mathbb{S}_{K-1}}{\operatorname{argmin}} \left\{ \sum_{i \in \boldsymbol{\mathcal{V}}} \left[ \frac{(y_i - M z_i s_i)^2}{2\sigma_y^2} + \lambda_x^\top z_i s_i \right] + \sum_{(i,j) \in \boldsymbol{\mathcal{E}}} z_i^\top \Sigma_x^{-1} z_j \right\}, \tag{17}$$

which is a quadratic programming of either $\{z_i\}_i$ or $\{s_i\}_i$ when the other set of variables is fixed. In cases where the reconstruction error $\sigma_y$ is large and the pairwise potential matrix $\Sigma_x^{-1}$ is also large, the programming of $\{z_i\}_i$ given $\{s_i\}_i$ is not positive semi-definite. We therefore use the iterated conditional model (ICM) [1] to find a local minimum, starting from $X^{(t-1)}$. In the ICM, we do local minimization for one cell at a time and repeat this multiple times until convergence. To do that, for cell $i$, we solve for $z_i$ and $s_i$ alternatively with the other fixed. The quadratic programming of $z_i$ given $s_i$ and other cells is positive semi-definite and therefore we solve it by Gurobi [2].

##### A.1.2 Derivation of optimization of model parameters

Given $X = X^{(t)}$, we aim to maximize the posterior probability of $\Theta$:

$$\Theta^{(t)} = \underset{\Theta}{\operatorname{argmax}} \; P\left(\Theta \mid Y, X\right) = \underset{\Theta}{\operatorname{argmax}} \; P\left(Y, X \mid \Theta\right) P(\Theta) \tag{18}$$

$$= \underset{\Theta}{\operatorname{argmax}} \; \exp\left(-\lambda_\Sigma \|\Sigma_x^{-1}\|_F^2\right) \frac{1}{Z(\Theta)} \prod_{(i,j)\in\boldsymbol{\mathcal{E}}} \varphi(x_i, x_j) \prod_{i\in\boldsymbol{\mathcal{V}}} \phi(y_i, x_i)\pi(x_i) \tag{19}$$

$$= \underset{\Theta}{\operatorname{argmax}} \; -\lambda_\Sigma \|\Sigma_x^{-1}\|_F^2 + \sum_{(i,j)\in\boldsymbol{\mathcal{E}}} \log\varphi(x_i, x_j) + \sum_{i\in\boldsymbol{\mathcal{V}}} \left(\log\phi(y_i, x_i) + \log\pi(x_i)\right) - \log Z(\Theta). \tag{20}$$

To simplify the partition function, we use the mean-field assumption:

$$P(X, Y \mid \Theta) \approx \prod_{i\in\boldsymbol{\mathcal{V}}} P(x_i, y_i \mid \Theta, X_{-i}, Y_{-i}) = \prod_{i\in\boldsymbol{\mathcal{V}}} P(x_i, y_i \mid \Theta, X_{-i}), \tag{21}$$

where

$$P(x_i, y_i \mid \Theta, X_{-i}) = \frac{1}{Z_i(\Theta)} \pi(x_i)\phi(y_i, x_i) \prod_{j\in\eta(i)} \varphi(x_i, x_j). \tag{22}$$

Then the programming becomes:

$$\underset{\Theta}{\operatorname{argmax}} -\lambda_\Sigma \|\Sigma_x^{-1}\|_F^2 + \sum_{i\in\boldsymbol{\mathcal{V}}} \left( \log\phi(y_i, x_i) + \log\pi(x_i) + \sum_{j\in\eta(i)} \log\varphi(x_i, x_j) - \log Z_i(\Theta) \right). \tag{23}$$

*Derivation of partition function*: Based on the mean-field assumption, we can write the partition function for each individual cell:

$$Z_i(\Theta) = \int_{x_i\in\mathbb{R}_+^K, y_i\in\mathbb{R}^G} \pi(x_i)\phi(y_i, x_i) \prod_{j\in\eta(i)} \varphi(x_i, \hat{x}_j)\, \mathrm{d}x_i\, \mathrm{d}y_i \tag{24}$$

$$= \int_{x_i\in\mathbb{R}_+^K} \pi(x_i) \prod_{j\in\eta(i)} \varphi(x_i, \hat{x}_j)\, \mathrm{d}x_i \int_{y_i\in\mathbb{R}^G} \phi(y_i, x_i)\, \mathrm{d}y_i. \tag{25}$$

We first calculate the inner integral of $y_i$, which is a Gaussian integral with mean $Mx_i$ and variance $\sigma_y^2$:

$$Z_i^Y(\Theta) := \int_{y_i\in\mathbb{R}^G} \phi(y_i, x_i)\, \mathrm{d}y_i = \int_{y_i\in\mathbb{R}^G} \exp\left[-\frac{(y_i - Mx_i)^2}{2\sigma_y^2}\right] \mathrm{d}y_i = \left(2\pi\sigma_y^2\right)^{G/2}, \tag{26}$$

which is independent of $x_i$ and is a function of $\sigma_y$. Then the partition function is simplified to

$$Z_i(\Theta) = Z_i^Y(\Theta) \cdot \int_{x_i\in\mathbb{R}_K} \pi(x_i) \prod_{j\in\eta(i)} \varphi(x_i, \hat{x}_j)\, \mathrm{d}x_i. \tag{27}$$

To simplify the integral of $x_i$, we again write $x_i$ as $s_i z_i$, where $s_i \in \mathbb{R}_+$ is a non-negative real number, representing the $L_1$ norm of $x_i$, and $z_i \in \mathbb{S}_{K-1}$ is a point in the $K$-dimensional simplex:

$$x_i = s_i z_i \quad \text{s.t.} \quad s_i \in \mathbb{R}_+, z_i \in \mathbb{S}_{K-1}. \tag{28}$$

The Jacobian of this transformation is $s_i^{K-1}$. Then the integral of $x_i$ becomes:

$$Z^X(\Theta) := \int_{x_i \in \mathbb{R}_+^K} \pi(x_i) \prod_{j \in \eta(i)} \varphi(x_i, \hat{x}_j)\, \mathrm{d}x_i \tag{29}$$

$$= \int_{s_i \in \mathbb{R}_+} \int_{z_i \in \mathbb{S}_{K-1}} \mathrm{d}s_i\, \mathrm{d}z_{i,1} \cdots \mathrm{d}z_{i,K-1} s_i^{K-1} \exp\left[-\lambda_x s_i - \sum_{j \in \eta(i)} \hat{z}_j^\top \Sigma_x^{-1} z_i\right] \tag{30}$$

$$= \left(\int_{s_i \in \mathbb{R}_+} \mathrm{d}s_i s_i^{K-1} \exp\left[-\lambda_x s_i\right]\right) \left(\int_{z_i \in \mathbb{S}_{K-1}} \mathrm{d}z_{i,1} \cdots \mathrm{d}z_{i,K-1} \exp\left[-\sum_{j \in \eta(i)} \hat{z}_j^\top \Sigma_x^{-1} z_i\right]\right). \tag{31}$$

The integral of $s_i$ is easy to compute,

$$Z^s(\Theta) := \int_{s_i \in \mathbb{R}_+} \mathrm{d}s_i s_i^{K-1} \exp\left[-\lambda_x s_i\right] = \lambda_x^{-K}(K-1)!. \tag{32}$$

The integral in $Z_i^z(\Theta)$ has analytic form:

$$Z_i^z(\Theta) := \int_{z_i \in \mathbb{S}_{K-1}} \mathrm{d}z_{i,1} \cdots \mathrm{d}z_{i,K-1} \exp\left[-\eta_i^\top z_i\right] = \sum_{k=1}^K \frac{\exp(-\eta_{ik})}{\prod_{j \neq k}(\eta_{ij} - \eta_{ik})}, \tag{33}$$

where $\beta_i = \Sigma_x^{-1} \sum_{j \in \eta(i)} \hat{z}_j$. To derive Eqn. 33, we first notice that the integral $Z_i^z(\Theta)$ is the value of the convolution of $\{h_{ik}\}_{k=1}^K$ evaluated at 1, denoted as $(*_{k=1}^K h_{ik})(1)$, where $h_{ik} : z_{ik} \mapsto \exp(-\eta_{ik} z_{ik})$. Because of the convolution theorem for the Laplace transform, the Laplace transform of $(*_{k=1}^K h_{ik})$ is $\boldsymbol{\mathcal{L}}[*_{k=1}^K h_{ik}] = s \mapsto \prod_{k=1}^K \frac{1}{\eta_{ik}+s}$. By partial fractions, it is easy to see that $\prod_{k=1}^K \frac{1}{\eta_{ik}+s}$ is equal to $\sum_{k=1}^K \frac{1}{(\eta_{ik}+s)\prod_{j \neq k}(\eta_{ij}-\eta_{ik})}$. Because the inverse Laplace transform $\boldsymbol{\mathcal{L}}^{-1}$ is linear, and because $\boldsymbol{\mathcal{L}}^{-1}[s \mapsto 1/(\alpha+s)] = t \mapsto e^{-\alpha t}$, we know $\boldsymbol{\mathcal{L}}^{-1}\left[s \mapsto \sum_{k=1}^K \frac{1}{(\eta_{ik}+s)\prod_{j \neq k}(\eta_{ij}-\eta_{ik})}\right] = t \mapsto \sum_{k=1}^K \frac{\exp(-\eta_{ik} t)}{(\eta_{ik}+s)\prod_{j \neq k}(\eta_{ij}-\eta_{ik})}$. Therefore, Eq. 33 holds. To avoid numerical instability, we use the first $n$ terms in Taylor's series of the exponentials, where $n$ is approximately the range of $\Sigma_x^{-1}$ times 1.1 plus $K-1$. This is more stable, because the first $K-1$ terms cancel out with each other. Then the formula can be rewritten as an $(n-K+1)$-D polynomial and can be computed in $O(nK)$ time and $O(K)$ space.

*Optimization of $M$*: The parameter $M$ appears in the $\log \phi(y_i, x_i)$ terms only. Therefore, to maximize the posterior probability, we solve the following positive semi-definite quadratic programming for $M$ by Gurobi [2],

$$M^{(t)} = \operatorname*{argmax}_{M \in \mathbb{S}_{G-1}^K} \sum_{i \in \boldsymbol{\mathcal{V}}} \log \phi(y_i, x_i) = \operatorname*{argmin}_{M \in \mathbb{S}_{G-1}^K} \sum_{i \in \boldsymbol{\mathcal{V}}} (y_i - Mx_i)^2. \tag{34}$$

*Optimization of $\sigma_y$*: The partial derivatives of related terms with respect to $\sigma_y^{-1}$ are:

$$\frac{\partial}{\partial \sigma_y^{-1}} \log \phi(y_i, x_i) = -\sigma_y^{-1}(y_i - Mx_i)^2 \qquad \frac{\partial}{\partial \sigma_y^{-1}} \log Z_i^Y(\Theta) = -\sigma_y G. \tag{35}$$

This implies that the objective function is convex and the global optimal point is:

$$\sigma_y^{(t)} = \sqrt{\frac{1}{NG} \sum_{i \in \boldsymbol{\mathcal{V}}} (y_i - Mx_i)^2}. \tag{36}$$

*Optimization of* $\Sigma_x^{-1}$: In the objective function, $\Sigma_x^{-1}$ appears in $\varphi(x_i, x_j)$ and $Z_i^z(\Theta)$, more specifically, $Z^z(\Theta)$. Therefore, we minimize:

$$-\sum_{i \in \boldsymbol{\mathcal{V}}} \sum_{j \in \boldsymbol{\mathcal{N}}(i)} \log \varphi(x_i, x_j^{\mathrm{MAP}}) + \sum_{i \in \boldsymbol{\mathcal{V}}} \log Z_i^z(\Theta) \tag{37}$$

by PyTorch [3] using Adam [4] as the optimizer. The learning rate is initialized to $10^{-2}$ and is multiplied by $0.98$ every 331 iterations.

#### A.2 Additional method details of simulation data generation and analysis

##### *A.2.1 Simulation methodology using metagenes*

For the metagene-based simulation methodology, the identity of a cell belonging to a specific type was defined by a specific mixture of metagenes associated with major-type, subtype, or layers, along with three noise metagenes (see **Fig.** 2b for the average proportions for each cell type). This pattern followed the observed trends of real mouse cortex data [5, 6]. For excitatory neurons, the layer-specific metagene defined the subtype. The two glial types had different major-type metagenes and no subtype metagenes. Given the class-specific metagene proportions, which we denote by the $K$-dimensional vector $b_c$ for cell type $c$, the proportions for an individual cell are generated as follows,

$$v_i = \frac{\tilde{v}_i}{\sum_k \tilde{v}_{i,k}} \qquad\qquad \tilde{v}_i = b_c + \eta_i,$$

where $\eta_i \sim \boldsymbol{\mathcal{N}}(0, \sigma_x \Sigma_c)$ is a $K$-dimensional Gaussian random variable that controls the cell-specific variation of metagene proportions. The diagonal matrix $\Sigma_c$ defines the variance for each metagene for cell type $c$. The parameter $\sigma_x$ is a scaling constant that controls the overall variance of metagene proportions for a given simulation. To simulate cell-specific variation of the number of total gene counts, we then scaled $v_i$ by a random scaling factor $s_i$ drawn from the Gamma distribution, with a shape parameter of $K$ and a scale parameter of 1. This yields the final hidden state

$$x_i = s_i v_i$$

for each cell $i$.

The procedure to generate metagenes is as follows. We randomly generated the value $M_{g,k}$ for the expression of gene $g$ for metagene $k$, but linked a significant percentage of the genes of subtype metagenes, such that the expression of those genes was the same across linked metagenes. For the two inhibitory neuron subtype-specific metagenes, 75% of the genes were linked. The four layer-specific metagenes had 90% of their genes linked together. We also linked 75% of the genes of the excitatory and inhibitory major-type metagenes, since they are both neural types. This pattern follows what was observed of metagenes learned from real spatial transcriptomic data in this work. We generated the value for each gene for each metagene from the Gamma distribution with a scale parameter of 1. The shape parameter for noise metagenes was 8, and the parameter for all other metagenes was 4. This achieved a desired level of sparsity among gene expression values. After the initial values for each gene were drawn, we normalized each metagene to sum to one. In this way, any scaling of the total number of counts per cell is captured entirely by the hidden variables $x_i$.

Given the randomized metagenes and hidden states, the observed expression $Y_{g,i}$ for cell $i$ for gene $g$ is a linear combination of the metagenes, with weights $x_i$, and with added Gaussian noise, $y_i = Mx_i + e_i$, where $e_i \sim \boldsymbol{\mathcal{N}}(0, \sigma_y^2 I_G)$ and $I_G$ is the identity matrix of dimension $G$. In our experiments, we set $G = 100$ genes.

To test the robustness of SpiceMix, we designed four different simulation scenarios following the above description, in which the values of $\sigma_y$ and $\sigma_x$ were varied (see **Fig.** 2d). For each scenario, we generated 20 replicates and reported the mean and confidence intervals of the adjusted Rand index (ARI) score across replicates.

##### *A.2.2 Simulation methodology using scDesign2*

The simulated gene expression of the second set of simulated data was generated by scDesign2 [7], in place of the metagene-based formulation described above. We trained the model of scDesign2 using scRNA-seq data of the mouse cortex from Tasic et al. [8]. We limited the gene set to the intersection of those measured in [8] and those measured by STARmap in [6], yielding a set of 969 genes. We used the following seven cell types for our simulation: L2/3, L4, L5, and L6 excitatory neurons; Pvalb and Vip inhibitory neurons; and oligodendrocytes (from both Opalin and 96*Rik subtypes). We then added a second Vip inhibitory neuron type that used the same generative model trained by scDesign2, but we added a slight constant offset to the expression of each cell to make the dataset more challenging. This cell type replaced the second glial type from our metagene-based experiment. To mimic the low counts of spatial transcriptomics data generated by STARmap or seqFISH+, we randomly discarded reads until the median total count per cell was 600, which is comparable to that of STARmap.

For this dataset, we devised two additional steps of introducing pairwise spatial dependency between cells. First, we added several diffusive spatial noise patterns, each with a unique associated noise vector that is added to the expression of each cell. The contribution of this noise term to the expression of a cell is scaled based on the distance of the cell from a defined center for each pattern and a defined spread for each pattern. Mathematically, this is expressed as follows:

$$y_i = y_i^{(sc)} + \sum_j \frac{1}{\sigma_{p_j}\sqrt{2\pi}} \exp\left\{-\frac{1}{2}\left(\frac{d(c_i, \mu_{p_j})}{\sigma_{p_j}}\right)^2\right\},$$

where $y^{(sc)}$ is the downsampled expression generated from scDesign2, $\sigma_{p_j}$ and $\mu_{p_j}$ are the spread and mean associated with noise pattern $p_j$, respectively, $c_i$ is the spatial coordinate of cell $i$, and $d(c_i, \mu_{p_j})$ is the Euclidean distance between $c_i$ and $\mu_{p_j}$. To ensure that the resulting counts are integers, the added noise is rounded. The noise for each cell for each gene is rounded up or down based on a Bernoulli distribution with probability equal to the fractional part of the noise. This ensures that the general trend of the noise pattern is preserved after rounding.

Second, we changed the association of some reads to a neighboring cell, to model the problem of leakage of counts in spatial transcriptomics data, which happens when cells overlap in the image plane or when there are errors in the segmentation of cell boundaries. For each read, the probability of being leaked is modeled by a Bernoulli random variable and the likelihood of any neighboring cell receiving the count is uniform.

We created 20 replicates and reported the mean and confidence intervals of the ARI score across replicates. We varied the ratio of the average amplitude of the signal (the downsampled counts from scDesign2) to the average amplitude of the noise to be between 2:1 to 1:1. For each experiment, we used either five or seven spatial noise terms. We varied the probability of a count leaking into a neighboring cell to be between 10% to 20%.

##### A.2.3 Additional method details for analyzing simulation data

For SPICEMIX and NMF, we tried at least 5 different random seeds for initialization and chose one based on the highest value of the $Q$ function obtained. To reduce the dimension of the input to Seurat, we used 10, 15, and 20 principal components and chose the value for each replicate that resulted in the highest ARI score. To identify cell types in the latent spaces of NMF and SPICEMIX, we used Louvain clustering. For SPICEMIX, NMF, and Seurat, we set the number of neighbors for Louvain clustering to be eight. To determine the resolution parameter for Louvain clustering, we used a binary search, with a range of $[0.5, 1.5]$, and stopped once the resulting number of clusters was eight. For NMF and SPICEMIX, since we desired each metagene to be given equal weight when defining cell identity, we z-score normalized the latent states along the cell dimension before clustering.

In order to select the hyperparameters for SpaGCN, we followed the tutorial provided by the authors in their GitHub repository. The tutorial includes methods for deciding the characteristic length scale, deciding a suitable resolution for initial clustering, and for determining which cells should be adjacent to each other in the neighborhood graph. Importantly, the SpaGCN pipeline requires the user to specify the number of clusters, which we set to eight.

To analyze the simulated data generated by the metagene-based approach, we tested values of $\lambda_\Sigma = 2|\boldsymbol{\mathcal{E}}| \times 10^{-4}$ and $\lambda_\Sigma = 2|\boldsymbol{\mathcal{E}}| \times 10^{-2}$ for the regularization parameter for SPICEMIX. We experimented with the number of metagenes $K$, which we ranged from 10 to 13, with a step size of 1. The results shown in **Fig.** 2 are for $K = 10$ for both SPICEMIX and NMF, which yielded the best performance for both, though we found the improvement in performance by SPICEMIX was similar for all values of $K$. We ran both NMF and SPICEMIX for 50 iterations. For the HMRF, we tried values between 0 to 10, with a step size of 1, for the spatial regularization term $\beta$. For each value, we tried 5 different random seeds. We reported the results for the value of $\beta$ and the random seed that yielded the best ARI for each replicate. The algorithm chooses the optimal number of hidden states based on a provided maximum number, which we set to be 10.

Since SpaGCN and Seurat assume the input data to be count-normalized and log-normal, we took the exponent of the expression in each cell, normalized the resulting expression of each cell to be 10,000, and then applied the log transform of the expression. For SpaGCN, we reported the 75th percentile of the ARI score obtained across random seeds for each replicate.

To analyze simulation data generated by scDesign2, we set $\lambda_\Sigma = 2|\boldsymbol{\mathcal{E}}| \times 10^{-4}$ for SPICEMIX. For SPICEMIX and NMF, we tried $K = 12$ and $K = 15$ metagenes and chose the value that resulted in the highest ARI for each replicate. This is comparable to our method for choosing the number of principal components for Seurat. We ran NMF for 100 iterations and SPICEMIX for 500 iterations. For SpaGCN, we reported the 50th percentile and 75th percentile of the ARI score obtained across random seeds for each replicate. Before running any method, we normalized the counts of each cell to sum to 10,000 and then applied the log transform.

#### A.3 Additional method details for the analysis of the seqFISH+ dataset

##### A.3.1 Preprocessing of seqFISH+ data

We first removed genes which had non-zero expression in less than 40% of cells, which yielded an unbiased set of 2,470 genes. We then normalized the expression of these genes by scaling the total counts to 10,000 per cell, adding one, and applying the log transform: $E'_{ig} := \log\left(1 + \left(10^4 \frac{E_{ig}}{\sum_{g'} E_{ig'}}\right)\right)$. To generate a graphical representation of the cells, we applied Delaunay triangulation to physical coordinates of cells, and then removed edges of length larger than 300 pixels (30.9 $\mu$m).

##### A.3.2 Selection of hyperparameters of algorithms

For the regularization parameter of the spatial pairwise dependency, $\lambda_\Sigma$, we considered possible values in the set $\{2|\boldsymbol{\mathcal{E}}| \times 10^{-2}, 2|\boldsymbol{\mathcal{E}}| \times 10^{-4}, 2|\boldsymbol{\mathcal{E}}| \times 10^{-6}\}$. We found $2|\boldsymbol{\mathcal{E}}| \times 10^{-4}$ to yield the desired balance of spatial regularization based upon visual inspection. We experimented with the number of metagenes, $K$, and chose the highest value before the expression of metagenes became too sparse. We also examined the UMAP plots of latent states, without annotations from the original analysis, to guide our selection. This led us to use $K = 20$ metagenes for both SpiceMix and NMF. For each hyperparameter configuration, we ran several iterations of the algorithm with different initial random seeds and chose the random seed that resulted in the highest value of the objective function, $Q$. After learning the latent states, we z-score normalized the latent states along the cell dimension and performed hierarchical clustering on the normalized latent states to define cell type assignment using Ward's method and the Euclidean distance [9]. We used the Calinski-Harabasz (CH) index [10] as the criterion for determining the optimal number of clusters. Before downstream analysis, we repeatedly merged the two clusters with the lowest threshold form hierarchical clustering until the last 3 splits did not create any cluster with less than five cells. We then eliminated outlier SpiceMix cell types that had less than five cells. This led to 15 cell types for SpiceMix and 13 cell types for NMF.

##### A.3.3 Selection of Louvain clusters used in comparison

In the analysis of the seqFISH+ dataset using Louvain clustering, reported in [5], cells from two regions (VISp and SVZ) were collected and clustered together. Since we studied the VISp only, we removed Louvain clusters identified in [5] that were enriched in SVZ. Specifically, for each Louvain cluster, we performed the Fisher's exact test and found that 10 Louvain clusters were enriched in the SVZ (clusters 1, 8, 12, 14, 15, 17, 21, 22, 24, and 26) with a $P$<0.05. We note that all of these clusters, except for cluster 26, had a $P < 10^{-6}$, so the choice of the threshold did not significantly affect the analysis. Moreover, only 13 cells from the VISp belonged to one of these clusters and meaningful association between these clusters and SpiceMix clusters was not possible. Thus, we omitted these 10 clusters from our comparative analysis.

##### A.3.4 Additional justification of SpiceMix excitatory subtypes in seqFISH+

The five excitatory subtypes in the cell type assignments of SpiceMix were further verified by comparing DEGs and the expression patterns of known marker gene to those in [8] (**Fig.** S5). All differential expressions were tested using MAST [11] via the interface provided in Seurat [12].

When finding DEGs, only genes that appeared in both [8] and [5] and that were expressed in at least 1% of cells in [8] were tested, which results in $n = 8,295$ genes. When finding DEGs for each cluster, only genes that were expressed at a higher level in that cluster were tested, but all genes were used to correct the $P$-values by FDR. DEGs were called in a one-versus-rest manner with a threshold on the FDR adjusted $P$-value less than 0.01 in either dataset separately. Since more subtypes were identified in [8], finer subtypes were merged according to their layers to form comparable clusters. To further justify the identify of the major excitatory type eL6, cells in the eL6a and eL6b subtypes of SpiceMix were merged, and all L6 subtypes in [8] were also merged. One-sided Fisher exact test was used to test the significance of the overlapping of DEGs.

When testing the differential expressions of individual known marker genes, the expression levels of all genes were fed into MAST and only the uncorrected $P$-values of target genes were extracted. In any such test, the results of at most two genes would be collected, so $P$-value correction is unnecessary.

#### A.4 Additional method details for the analysis of the STARmap dataset

##### A.4.1 Preprocessing of STARmap data

We normalized the data by scaling the total counts to 10,000 per cell, adding one, and applying the log transform: $E'_{ig} := \log\left(1 + \left(10^4 \frac{E_{ig}}{\sum_{g'} E_{ig'}}\right)\right)$. To generate a graphical representation of the cells, we applied Delaunay triangulation to physical coordinates of cells, and then removed edges of length larger than 600 pixels.

##### A.4.2 Selection of hyperparameters of algorithms

For the regularization parameter of the spatial pairwise dependency, $\lambda_\Sigma$, we considered possible values in the set $\{2|\boldsymbol{\mathcal{E}}| \times 10^{-2}, 2|\boldsymbol{\mathcal{E}}| \times 10^{-4}, 2|\boldsymbol{\mathcal{E}}| \times 10^{-6}\}$. We found $2|\boldsymbol{\mathcal{E}}| \times 10^{-4}$ to yield the desired balance of spatial regularization based upon visual inspection. We experimented with the number of metagenes, $K$, and chose the highest value for each algorithm before the expression of metagenes became too sparse. We also examined the UMAP plots of latent states, without annotations from the original analysis, to guide our selection. This led us to use $K = 20$ metagenes for SPICEMIX and $K = 15$ metagenes for NMF. For each hyperparameter configuration, we ran several iterations of the algorithm with different initial random seeds and chose the random seed that resulted in the highest value of the objective function, $Q$. After learning the latent states, we z-score normalized the latent states along the cell dimension and performed hierarchical clustering on the normalized latent states to define cell type assignment using Ward's method and the Euclidean distance [9]. We used the CH index as the criterion for determining the optimal number of clusters. Before downstream analysis, we removed an outlier SPICEMIX cell type that had only one cell. This led to 16 cell types for SPICEMIX and 11 cell types for NMF.

We used the implementation of HMRF from [13]. We ran HMRF on the same input file with a limit of 12∼16 cell types and scaling factors ranging from 0 to 30. For scHPF, we manually set its two hyperparameters to the default value and varied the number of latent factors to be between 15 and 20. We found that 16 latent factors produced the most clusters. For each number of factors, the number of clusters was chosen to maximize the CH index. We also performed analysis with scHPF setting the hyperparameter values automatically and again using 16 latent factors.

##### A.4.3 Monocle2 trajectory analysis

To investigate the expression of oligodendrocytes for evidence of a trajectory among such cells, we first separated out the labeled oligodendrocytes and OPCs, according to SPICEMIX cell-type assignments. We used the expression values of these cells as input to Monocle2 [14]. We filtered out genes that were not expressed above a minimum threshold of 1 count in at least 30% of cells. We used PCA to reduce the dimension of the remaining genes. We clustered cells in the dimension-reduced space using the implementation of the algorithm of Rodriguez and Laio in Monocle2 [14] and then found differentially-expressed genes across the clusters. These genes were then used by Monocle2 to order the cells and infer trajectory.

##### A.4.4 GO enrichment analysis of myelin sheath formation

We determined the genes related to myelin sheath formation by the gene ontology (GO) database [15, 16]. First, we searched for myelin sheath-related GO terms by looking for the occurrences of either 'myelin' or 'sheath' in the description of GO terms with GOATOOLS [17]. We then filtered out irrelevant GO terms, such as cellular components in neural cells or sperm, biological processes of periphery neural systems, and all parents and ancestors of any GO terms discovered in the previous step. Third, we queried the genes associated with these GO terms and dropped GO terms without any genes included in the STARmap dataset. The final list of GO terms was narrowed down to: GO:0014003, GO:0021779, GO:0048709, GO:0021529, GO:0032288, GO:0030913, GO:1990227, GO:0045163, GO:0022010, GO: 0031642, GO:0031643, GO:0019911, GO:0035749, GO:0097456, GO:0043209, GO:0043218. Fourth, we kept genes that (1) were associated with any of the GO terms, (2) were included in the STARmap dataset, and (3) were expressed in at least 40% of OPCs and 40% of Oligo-1 cells. We then used linear regression to analyze the relation between expression levels of these myelin sheath-related genes and those of SPICEMIX metagenes in individual cells. Specifically, we regressed the expression level of each myelin sheath-related gene against the difference between the proportions of metagenes 12 and 13: $(X_{13,i} - X_{12,i})/(\sum_{k=1}^{20} X_{k,i})$ for cell $i$, with the Python package scikit-learn [9]. The *P*-value of the hypothesis that the slope was nonzero was also calculated by the scikit-learn function. Then, we corrected *P*-values using the Benjamini-Hochberg method with the Python package statsmodel [18].

#### A.5 Additional method details for the analysis of the Visium dataset

##### A.5.1 Preprocessing of Visium data

For analysis with SPICEMIX, we removed genes which had non-zero expression in less than 10% of spots, which yielded an unbiased set of 3,194 genes. We did not apply this filtering when using SpaGCN or BayesSpace. We then normalized the expression of these genes by scaling the total counts to 10,000 per spot, adding one, and applying the log transform: $E'_{ig} := \log\left(1 + \left(10^4 \frac{E_{ig}}{\sum_{g'} E_{ig'}}\right)\right)$. To generate a graphical representation of the spots, we defined the neighborhood of a spot to be the set of directly adjacent spots in the hexagonal grid, since the spots in each FOV form a hexagonal grid. Therefore, except for spots on the edge of the grid, each spot has exactly 6 neighbors.

##### A.5.2 Sample selection for ARI score comparison

Since the four FOVs from brain sample Br5595 lacked layer L1 and included ambiguous spots, we excluded these FOVs from our evaluation. The four FOVs from brain sample Br5292 were also excluded from the ARI score comparison because SPICEMIX identified finer structures in them (as we demonstrated), which is an improvement but has negative impact to the ARI score which assesses the similarity. Therefore, we focused only on the 4 FOVs from brain sample Br8100. Additionally, the other methods in our comparison featured FOV 151673 in their analysis, which is from sample Br8100.

##### A.5.3 ARI score comparison on brain sample Br8100

To benchmark SPICEMIX's performance on the Visium dataset, we evaluated its performance for identifying layers using the ARI score. In this task, the desired labels are smooth layers whose widths are generally 3$\sim$10 or more spots, whereas SPICEMIX, as an unsupervised method, identified spot-to-spot variability within layers, providing insights at a much higher resolution than layers. To mitigate the variability, we performed smoothing on the spot embeddings and cluster labels. The complete pipeline of clustering on the Visium dataset can be summarized as follows:

1. Run SPICEMIX jointly on the four FOVs 151673-151676 from brain sample Br8100 with $K = 20$ metagenes and $\lambda_\Sigma = 10^{-12}$.
2. Define an expression score for each metagene as the average expression in the top 1% spots, and

filter out metagenes with low expression scores by the elbow plot.

3. Divide the expression levels of each metagene by the sum of its expressions in the top 1% spots per FOV.
4. Smooth metagene expression levels to the average of itself and its neighbors:

$$X'_k := \frac{\sum_{j \in \boldsymbol{\mathcal{N}}(k) \cup \{k\}} X_j}{|\boldsymbol{\mathcal{N}}(k)| + 1}$$

where $X_k$ is the vector-valued embedding of spot $k$, and $\boldsymbol{\mathcal{N}}(k)$ represents the neighbors of spot $k$ in a hexagonal grid.

5. Run Louvain clustering using the `scanpy` package's `louvain` API, with `n_neighbors=20` and `resolution=0.35`.
6. Smooth cluster labels until convergence. Specifically, let $L_k^0$ be the raw cluster label of spot $k$ produced in the previous step, and repeat the following process iteratively starting with $t = 1$:

$$L_k^{t+1} := \begin{cases} \ell & \frac{1}{|\boldsymbol{\mathcal{N}}(k)|+1} \sum_{j \in \boldsymbol{\mathcal{N}}(k) \cup \{k\}} \mathbb{1}(L_j^t = \ell) > 0.5 \\ L_k^t & \text{no such label } \ell \text{ exists} \end{cases}$$

We compared the ARI score attained by SPICEMIX with that of BayesSpace [19] and SpaGCN [20] based on our own experiments using the provided code. For SpaCGN, we ran the method two ways: using only one FOV at a time for the input or using all four FOVs simultaneously as input. We chose the SpaGCN hyperparameters using the methods in the authors' public GitHub repository. For BayesSpace, since there was no provided method to run on multiple FOVs at once, we only ran the method individually on each FOV. We loaded and preprocessed the Visium dataset using the methods provided in the BayesSpace R package. For both methods, we replicated each experiment ten times and took the average ARI score as the comparison benchmark. For all methods, the number of clusters was set to be seven, the number of annotated layers in the ground truth.

##### A.5.4 Detailed analysis procedure on brain sample Br8100

To interpret the metagenes learned from the 4 FOVs from brain sample Br8100, we identified differentially expressed genes (DEGs) from a snRNA-seq study [21] and examined these genes' ranks in the metagenes. Specifically, we downloaded the gene expression aggregated per cell type, calculated as trimmed means. For astrocytes, since three astrocyte clusters were identified in this study, we defined a gene as a DEG of astrocytes if the lowest expression of that gene among the three astrocyte clusters was greater than its expression in any other clusters. We used a similar definition for oligodendrotyces. We selected cluster 'Exc L2-3 LINC00507 RPL9P17' to represent the excitatory neurons residing in superficial layers (labeled as Exc (S) in **Fig.** 5d), because this cluster is the most superficial one according to the annotations. We then defined a gene as a DEG of Exc (S) if its expression in cluster 'Exc L2-3 LINC00507 RPL9P17' is greater than the 0.8th quantile of its expression in the other clusters. For deep excitatory neurons (labeled as Exc (D) in **Fig.** 5d), we selected the 12 excitatory clusters that were annotated to be located in layer L6. We defined a gene as a DEG of Exc (D) if the 0.2th quantile of its expression among these 12 clusters is greater than the 0.8th quantile of its expression in the other clusters. To compare the ranks of these DEGs in learned metagenes, we used the Wilcoxon signed-rank test between every pair of metagenes.

To confirm that the sulcal and gyric regions have different expression patterns, we first divided the cortex into two halves which approximate the sulcal region and gyric regions and then called DEGs between the two regions. The detailed process is as follows. (1) In each of the 4 FOVs, we found the spot with coordinate $(x, y)$ that maximizes the quantity $x + y$ among the spots annotated to the white matter (colored in cyan in **Fig.** S21), and the one among those annotated to the cortex (colored in green in **Fig.** S21). (2) In each FOV, we connected the two spots found in step (1) and used this line as the boundary to split layers L3 and L4 into two parts. We defined the lower-right part as the gyric region, and the upper-left part as the sulcal region. (3) We collected spots from layers L3 and L4 of all FOVs and called DEGs between the spots in the sulcal regions and the spots in the gyric regions, using MAST [11] via Seurat [22], using $10^{-40}$ as the *P*-value threshold for significance.

##### *A.5.5 Detailed analysis procedure on FOV 151507 of brain sample Br5292*

We ran SPICEMIX with $K = 20$ metagenes and $\lambda_\Sigma = 10^{-12}$ on FOV 151507. We estimated the intensity at each spot and divided spots into four regions by thresholding. In the dataset, one spot was expected to have a diameter of 50$\mu$m and the physical distance between neighboring spots was 100$\mu$m. Since the distance between neighboring spots was ~6 pixels on the histological image, we approximated the intensity of each spot by (1) converting the histological image to gray scale, (2) finding pixels whose distances to that spot were at most 1.5 pixels, and (3) averaging the intensities of these pixels weighted by the reciprocal distance to the spot. This process was implemented via the `RadiusNeighborsRegressor` class of the scikit-learn package [9]. Next, we defined the dark strip as the set of spots with an intensity less than 0.52, the bright gap as the set of spots with an intensity greater than 0.627, and the cortex to be the remaining spots. After the initial thresholding, we observed that a few (less than five) spots were far away from the dark strip and the bright gap, but were classified into the dark strip or the bright gap. Therefore, we further refined the region assignments by reassigning spots that were 20 pixels away from the dark strip and the bright gap to the cortex. Since the dark strip and the bright gap were very thin, a significant portion of spots lay on the boundaries between the dark strip, the bright gap, and the flanking cortex. These spots were mixtures of these three regions with high probability and their estimated intensities may not reflect their identities. Hence, we drew one boundary parallel to the fitted layer boundaries between cortical layers on each side of the dark strip and the bright gap and reassigned cortical spots between these two boundaries as mixtures.

To identify differential expressions within layer L1 and the white matter, we filtered out genes that were expressed in less than 10% of cells in the corresponding region. For the identification of DEGs in the white matter, we corrected *P*-values by the Benjamini/Hochberg procedure. No multi-testing correction was applied to the *P*-values of marker genes of mural cells, since we were querying for particular genes.

#### A.6 Additional method details

##### *A.6.1 Doublet detection*

To confirm that our results were not artifacts derived from doublets, we used DoubletDetect [23] to identify potential doublets. Specifically, we trained boost classifiers on the raw read count matrix with parameters `use_phenograph=True` and `standard_scaling=False`, and then used them to infer doublets with parameters `p_thresh=1e-8` and `voter_thresh=0.5`. To avoid the effect of randomness in this algorithm, we trained 10 boost classifiers with different random seeds on each dataset, and we define a cell as a potential doublet if any of the 10 classifiers identified it.

##### A.6.2 Calculation of cell-type affinity

In the neighbor graph, each vertex represents one cell and every edge connecting two vertices indicates that the two corresponding cells are neighbors. To quantify the affinity between two cell types, i.e., two sets of vertices in the neighbor graph, we constructed a random graph by shuffling edges in the original neighbor graph while keeping the number of neighbors of every cell unchanged, and compared the actual number of edges across these two cell types with the expected number.

Specifically, we cut every edge into two halves which we call stubs, and then we rewired each stub into any other stub in this network uniformly at random to create an edge. In this procedure, we allow an edge connecting a vertex to itself and multiple edges between the same pair of vertices. In a graph of $n$ vertices and $m$ undirected edges, we create $2m$ stubs, and any vertex $u$ with degree $k_u$ is associated with $k_u$ stubs. Now we consider one stub of vertex $u$ and pair it with any of the remaining $2m-1$ stubs uniformly at random. This stub is wired to another stub of vertex $u$, i.e., creating a self-loop, with probability $(k_u-1)/(2m-1)$, and it is wired to a stub of another vertex $v \neq u$ with probability $k_v/(2m-1)$. Hence, the expected number of self-loops of vertex $u$ is:

$$\frac{1}{\binom{2m}{k_u}\binom{2m-1}{k_u-1}} = \frac{k_u(k_u-1)}{2m-1}.$$

There are $2k_uk_v/(2m-1)$ edges between vertices $u$ and $v$ in expectation. Moreover, let $K_U$ denote the total number of degrees of vertices in vertex set $U$, and then, the expected number of edges connecting two vertices in $U$, including self-loops, is $K_U(K_U-1)/(2m-1)$. For two different sets of vertices $U$ and $V$, the expected number of edges connecting $u \in U$ and $v \in V$ is $2K_UK_V/(2m-1)$.

To quantify the affinity between two cell types, we subtract the expected number of edges between the two corresponding vertex sets in the random graph from the total number of edges between the two cell types in the original neighbor graph. We denote the difference for cell types $i$ and $j$ by $A_{i,j}$, and define the affinity between the two cell types as $A_{i,j}/\sqrt{K_iK_j}$. In other words, we normalize the deviation of the number of edges from the expected number by the square root of the product of the sizes of the two cell types, in order to eliminate the effect of cell-type sizes. We chose the normalization factor as $\sqrt{K_i}$ instead of $K_i$ because the average vertex degree is a constant.

### B Supplementary Results

#### B.1 Additional details of scDesign2 simulation results

SpiceMix outperformed other methods across five of the six scenarios, and tied with NMF in one scenario (**Fig.** S2a). NMF and Seurat performed comparably (**Fig.** S2a). SpaGCN was highly sensitive to both the spatial noise and leakage (**Fig.** S2a). The metagenes of SpiceMix captured both sparse and layer-specific patterns of expression (**Fig.** S2b). They were also able to disentangle the the effects of spatial noise and leakage (**Fig.** S2b). The metagene capturing oligodendrocyte expression was expressed at low levels in cells neighboring oligodendrocytes, which is likely capturing the leakage of oligodendrocyte expression into those cells (**Fig.** S2b). This demonstrates an advantage of the metagene formulation for this particular type of noise. SpiceMix was also able to learn metagenes that capture the expression patterns of the spatial noise terms (**Fig.** S2b). As a result of not modeling spatial information, NMF and Seurat incorrectly mixed the two similar inhibitory neuron types (i1 and i3), which had distinct layer enrichment, whereas SpiceMix missed only a few stray cells (**Fig.** S2c).

#### B.2 Additional details of seqFISH+ results

The excitatory neuron types of SpiceMix exhibit strong layer-specificity (**Fig.** 3d), which is consistent with findings from scRNA-seq studies [8]. In contrast, several of the excitatory Louvain clusters reported by [5] were dispersed across as many as three layers (see Fig. 3h in [5]). Within a layer, some excitatory neuronal types can be further separated into subtypes [8]. SpiceMix identified two L6 subtypes (eL6a and eL6b), which were mixed together across several clusters by [5], in addition to L2/3, L4, and L5 types (**Fig.** 3c (middle), **Fig.** S5). The expression of marker genes *Col6a1* and *Ctgf* [8] confirmed the labelling of eL6 subtypes by SpiceMix (**Fig.** 3c (left), **Fig.** S5). Eng et al. [5] identified 10 excitatory neuronal subtypes, in contrast to the five total excitatory types of SpiceMix, however, based on the expression of known marker genes identified in [8], no more than four of these clusters could be confirmed to match known excitatory subtypes. The auxiliary analysis of spatial domains using the HMRF-based method of Zhu et al. [13], as reported in [5], also could not separate the eL6a and eL6b subtypes, despite explicitly modeling spatial relationships. NMF also identified strong layer-specificity of excitatory neurons (**Fig.** S4), which indicate that this improvement is not an artifact of spatial smoothing and may rather be due in large part to the matrix factorization formulation of both NMF and SpiceMix. Among glial cells, both SpiceMix and NMF separated a Louvain cluster of [5] into SMC and Endo cells, which was confirmed by the expression of their respective marker genes (i.e., *Bgn* highly expressed in SMC but not Endo cells, and *Flt1* highly expressed in both SMC and Endo cells [8]) (**Fig.** 3c). The subsequent analysis of the method of Zhu et al. [13], as reported in [5], also could not clearly separate this Louvain cluster into Endo and SMC types (**Fig.** S3b).

We found that increasing $\lambda_x$ to 10 and/or increasing the sparsity threshold to $80\%$ in NMF had minimal effect on the cell type assignments (**Fig.** S7a,b,d,f), except that Micro was merged into Endo when $\lambda_x = 10$ without using thresholding (**Fig.** S7d). However, further increasing sparsity by either method actually had a negative impact on the cell type assignments of NMF. Importantly, when the threshold was $0.8$, the eL5 subtype disappeared and eL5 neurons were mixed into eL4, eL6, and inhibitory clusters (**Fig.** S7c,f,i). Also, the Micro subtype was merged to Endo when $\lambda_x = 10$ and to Oligo-1 when $\lambda_x = 100$. When the value of $\lambda_x$ was further increased to 100 (**Fig.** S7g,h,i), eL5 neurons were further split into multiple clusters that were merged into eL4, eL6, and/or inhibitory clusters. The only setting where inhibitory neurons were separated to two clusters was with $\lambda_x = 100$ and the sparsity threshold set to $0.8$, but one of the two clusters was incorrectly mixed with eL5 neurons. Hence, increasing sparsity by either or both methods cannot improve the performance of NMF and the advantage of SPICEMIX remained.

##### B.3 Additional details of STARmap results

We included the complete results of NMF, HMRF, and scHPF to compare to SPICEMIX (**Fig.** S8, **Fig.** S11, and **Fig.** S17, respectively). NMF revealed four excitatory neuron subtypes, two inhibitory neuron subtypes, and five glial subtypes. Both SPICEMIX and NMF revealed a clearer delineation of eL2/3 and eL4 neurons than the assignments of [6]. SPICEMIX reassiged 36 cells in excitatory subtypes eL2/3 and eL4, especially near the layer boundary (**Fig.** S9d). We found that at least 15 eL2/3* or eL4* cells of [6] resided not in layers L2-L4 but in layers L5 and L6, and SPICEMIX did not label them as eL2/3 or eL4 neurons, accordingly. We examined the expression of marker genes among the eL2/3 and eL4 cells for which SPICEMIX and [6] disagreed, which showed a clearer agreement of SPICEMIX assignments with a scRNA-seq study [8] than the assignments of [6] (**Fig.** S9a). Specifically, all of the eL4 markers (*Rorb*, *Whn*, and *Rspo1*) were expressed at higher levels among the eL2/3* cells, whereas for SPICEMIX, these markers were expressed at higher levels in the eL4 cells, following the trends of [8]. NMF changed 26 eL2/3 and eL4 neurons. However, the eL2/3 and eL4 types of the NMF assignment are not as constrained in one spatial layer as the assignments of SPICEMIX (**Fig.** S8b). Both SPICEMIX and NMF reassigned a large set of astrocytes, as labeled by [6], to eL5 neurons (**Fig.** 4c (middle) and **Fig.** S8a (middle), respectively). Astrocytes are known to be dispersed throughout the primary visual cortex, unlike the highly-localized Astro-1* cluster of [6] in the L5 layer (**Fig.** S10). Differential expression analysis showed that many of the marker genes of astrocytes (identified by [8]), such as *F3* and *Sox9*, are expressed in Astro-1* at a level significantly lower than those in Astro-2* and comparable to those in eL5 neurons (**Fig.** S10), confirming this assignment. Furthermore, the Astro-1* cells express marker genes of excitatory neurons, such as *Slc17a7*, *Tcerg1l*, *Stac*, and *Parm1*, at much higher levels than Astro-2* cells (**Fig.** S10).

We found that increasing $\lambda_x$ to 10 and/or increasing the sparsity threshold to $80\%$ did not improve the cell type assignments of NMF. Importantly, when $\lambda_x$ was 1 or 10, (1) NMF always produced a cluster that was a mixture of excitatory neurons and endothelial cells (Endo in **Fig.** S15a and Exc/Endo in **Fig.** S15b,c,d,e,f); (2) NMF could not further separate PVALB and SST; and (3) NMF could not identify finer oligodendrocyte clusters. When $\lambda_x$ was further increased to 100, the latent states became overly sparse and NMF produced many small and meaningless clusters. We also note that we ran NMF with $K = 20$ metagenes and the results were similar to that of $K = 15$ metagenes. NMF was still unable to discover finer cell types such as the oligodendrocyte types. This confirmed that the difference between the results from SPICEMIX and NMF was not because of the different numbers of metagenes.

The recent latent factor method scHPF [24] was unable to identify several spatial subtypes that were discovered by SPICEMIX. Specifically, when using automatic parameter selection, scHPF did not identify Astro-2, OPC, and the split of eL6 into eL6a, eL6b, and eL6c (**Fig.** S17a). Using manual parameter selection yielded more subtypes, namely the Reln inhibitory subtype and a split of eL4 neurons (which we did not validate), but conversely, the method did not identify Astro-2, OPC, and the spatially-distinct Oligo-1 and Oligo-2 split, nor still did it identify the split of eL6 into eL6a, eL6b, and eL6c (**Fig.** S17a). This also resulted in the mixing of some SST neurons into the PVALB subtype. Moreover, the *in situ* maps of the cell assignments of scHPF revealed significant intermixing of excitatory neurons between layers (**Fig.** S17b). The *in situ* maps of the expression of the latent factors of scHPF revealed that the latent factors were not as layer-specific as those of SPICEMIX (**Fig.** S17c), though they were less diffuse than those of NMF.

HMRF almost always output a degenerative case where cells are clustered into at most 11 cell types, despite the choice of the initial number of cell types or the smoothing parameter, which performs poorly as compared to the assignments of SPICEMIX or of [6].

###### B.3.1 Metagene mixture of OPC and Oligo-1 in STARmap data

Because Astro-2/OPC is a mixture of OPCs in the deep layer and astrocytes in the most superficial layer, we kept only the cells of this cluster that are located in the superficial layer in the following analysis. These astrocytes correspond to the thin sliver of magenta-colored cells at the superficial extremity of the tissue, whereas the remainder of Astro-2/OPC cells, which we posit are oligodendrocytes, are located in a deep-tissue layer near Oligo-1 cells (**Fig.** 4c). Additionally, the expression of astrocyte and oligodendrocyte marker genes among these astrocytes clearly resembles that of the STARmap Astro-2* class, whereas the expression of such genes among the oligodendrocytes of the Astro-2/OPC class resembles that of the Oligo* class (**Fig.** S14a,c,d). We expect that these astrocytes and oligodendrocytes were grouped together due to their overall similar gene expression profiles, as seen in a UMAP plot of the cells in gene expression space (**Fig.** S14c), despite their distinct spatial enrichment patterns. We also note that the metagenes primarily expressed by cells within both groups of the Astro-2/OPC class exhibited strong spatial affinity for themselves (**Fig.** 4e), which increased the likelihood for SPICEMIX to group them together.

Out of the 11 genes, 7 were significantly correlated with the difference of metagene proportions. We reasoned that the expression levels of the other 4 genes either were supposed to be irrelevant to the myelin sheath formation or were inaccurate because of potential incorrect image segmentation and assignments of mRNAs to cells. The mouse *Cd9* gene encodes a protein which was reported to participate in paranodal junction assembly [25] that happened in mature oligodendrocytes but not in OPCs, and was observed in the mouse central nervous system during development, but not in mature mouse [26]. However, in a recent scRNA-seq study [8], *Cd9* mRNA was observed in both OPCs and mature oligodendrocytes following extremely similar distributions, which is consistent with our finding on the STARmap dataset and explains why the expression level of *Cd9* mRNA is not correlated with differentiation from OPCs to oligodendrocytes. We therefore hypothesize that *Cd9* mRNAs are present in OPCs in mature mice, but *Cd9* proteins are absent or are maintained at a low level. The mouse *Gfap* gene encodes the glial fibrillary acidic protein and is a widely used biomarker of astrocytes in the central nervous system and an indicator of many activities of astrocytes [27]. The participation of *Gfap* in myelin sheath formation was reported by [28]. However, as mentioned in [28], the presence of *Gfap* was more likely to be a false positive due to the occasional labeling of astrocytes.

In the high-quality scRNA-seq data [8], all genes but *Atp1a2* were expressed in most mature oligodendrocytes, but only a few OPCs, and the expression levels in oligodendrocytes were significantly higher than those in OPCs, whereas *Atp1a2* exhibited the opposite pattern. In other words, most OPCs and oligodendrocytes could be undoubtedly delineated only by one gene. In contrast, the OPCs and oligodendrocytes cannot be clearly distinguished from each other even based on expression levels of multiple genes. This suggests that the STARmap dataset sacrificed accuracy of mRNA levels to cells' spatial locations, and that SPICEMIX is able to characterize cells under a large noise level.

#### References


[1] Besag J. On the statistical analysis of dirty pictures. Journal of the Royal Statistical Society: Series B (Methodological). 1986;48(3):259–279.

[2] Gurobi Optimization L. Gurobi Optimizer Reference Manual; 2020. Available from: http://www.gurobi.com.

[3] Paszke A, Gross S, Chintala S, Chanan G, Yang E, DeVito Z, et al. Automatic Differentiation in PyTorch. In: NIPS Autodiff Workshop; 2017. .

[4] Kingma DP, Ba J. Adam: A method for stochastic optimization. arXiv preprint arXiv:14126980. 2014.

[5] Eng CHL, Lawson M, Zhu Q, Dries R, Koulena N, Takei Y, et al. Transcriptome-scale super-resolved imaging in tissues by RNA seqFISH+. Nature. 2019;568:235–239.

[6] Wang X, Allen WE, Wright MA, Sylwestrak EL, Samusik N, Vesuna S, et al. Three-dimensional intact-tissue sequencing of single-cell transcriptional states. Science. 2018;341(6400):eaat5691.

[7] Sun T, Song D, Li WV, Li JJ. scDesign2: a transparent simulator that generates high-fidelity single-cell gene expression count data with gene correlations captured. Genome Biology. 2021;22(1):1–37.

[8] Tasic B, Menon V, Nguyen TN, Kim TK, Jarsky T, Yao Z, et al. Adult mouse cortical cell taxonomy revealed by single cell transcriptomics. Nature Neuroscience. 2016;19(2):335.

[9] Pedregosa F, Varoquaux G, Gramfort A, Michel V, Thirion B, Grisel O, et al. Scikit-learn: Machine Learning in Python. Journal of Machine Learning Research. 2011;12:2825–2830.

[10] Caliński T, Harabasz J. A dendrite method for cluster analysis. Commun Stat Simul Comput. 1974 Jan;3(1):1–27.

[11] Finak G, McDavid A, Yajima M, Deng J, Gersuk V, Shalek AK, et al. MAST: a flexible statistical framework for assessing transcriptional changes and characterizing heterogeneity in single-cell RNA sequencing data. Genome Biology. 2015;16(1):1–13.

[12] Butler A, Hoffman P, Smibert P, Papalexi E, Satija R. Integrating single-cell transcriptomic data across different conditions, technologies, and species. Nature Biotechnology. 2018;36(5):411–420.

[13] Zhu Q, Shah S, Dries R, Cai L, Yuan GC. Identification of spatially associated subpopulations by combining scRNAseq and sequential fluorescence in situ hybridization data. Nature Biotechnology. 2018;36(12):1183.

[14] Qiu X, Mao Q, Tang Y, Wang L, Chawla R, Pliner HA, et al. Reversed graph embedding resolves complex single-cell trajectories. Nature Methods. 2017;14(10):979–982.

[15] Ashburner M, Ball CA, Blake JA, Botstein D, Butler H, Cherry JM, et al. Gene ontology: tool for the unification of biology. Nature Genetics. 2000;25(1):25–29.

[16] Consortium GO. The gene ontology resource: 20 years and still GOing strong. Nucleic Acids Research. 2019;47(D1):D330–D338.

[17] Klopfenstein D, Zhang L, Pedersen BS, Ramírez F, Vesztrocy AW, Naldi A, et al. GOATOOLS: A Python library for Gene Ontology analyses. Scientific Reports. 2018;8(1):1–17.

[18] Seabold S, Perktold J. statsmodels: Econometric and statistical modeling with python. In: 9th Python in Science Conference; 2010. .

[19] Zhao E, Stone MR, Ren X, Guenthoer J, Smythe KS, Pulliam T, et al. Spatial transcriptomics at subspot resolution with BayesSpace. Nature Biotechnology. 2021;39(11):1375–1384.

[20] Hu J, Li X, Coleman K, Schroeder A, Ma N, Irwin DJ, et al. SpaGCN: Integrating gene expression,

spatial location and histology to identify spatial domains and spatially variable genes by graph convolutional network. Nature Methods. 2021;18(11):1342–1351.

[21] Dataset: Allen Institute for Brain Science (2021). Allen Cell Types Database – Human Multiple Cortical Areas [dataset]. Available from: http://celltypes.brain-map.org/rnaseq;.

[22] Satija R, Farrell JA, Gennert D, Schier AF, Regev A. Spatial reconstruction of single-cell gene expression data. Nature Biotechnology. 2015;33(5):495.

[23] Gayoso A. Shor J. JonathanShor/DoubletDetection: Doubletdetection V3. 2018.

[24] Levitin HM, Yuan J, Cheng YL, Ruiz FJ, Bush EC, Bruce JN, et al. De novo gene signature identification from single-cell RNA-seq with hierarchical Poisson factorization. Molecular Systems Biology. 2019;15(2):e8557.

[25] Ishibashi T, Ding L, Ikenaka K, Inoue Y, Miyado K, Mekada E, et al. Tetraspanin protein CD9 is a novel paranodal component regulating paranodal junctional formation. Journal of Neuroscience. 2004;24(1):96–102.

[26] Terada N, Baracskay K, Kinter M, Melrose S, Brophy PJ, Boucheix C, et al. The tetraspanin protein, CD9, is expressed by progenitor cells committed to oligodendrogenesis and is linked to $\beta 1$ integrin, CD81, and Tspan-2. Glia. 2002;40(3):350–359.

[27] Eng LF, Ghirnikar RS, Lee YL. Glial fibrillary acidic protein: GFAP-thirty-one years (1969–2000). Neurochemical Research. 2000;25(9-10):1439–1451.

[28] Werner HB, Kuhlmann K, Shen S, Uecker M, Schardt A, Dimova K, et al. Proteolipid protein is required for transport of sirtuin 2 into CNS myelin. Journal of Neuroscience. 2007;27(29):7717–7730.

[29] Marques S, Zeisel A, Codeluppi S, van Bruggen D, Falcão AM, Xiao L, et al. Oligodendrocyte heterogeneity in the mouse juvenile and adult central nervous system. Science. 2016;352(6291):1326–1329.

[30] Maynard KR, Collado-Torres L, Weber LM, Uytingco C, Barry BK, Williams SR, et al. Transcriptome-scale spatial gene expression in the human dorsolateral prefrontal cortex. Nature Neuroscience. 2021 Mar;24(3):425–436.

#### Supplementary Figures

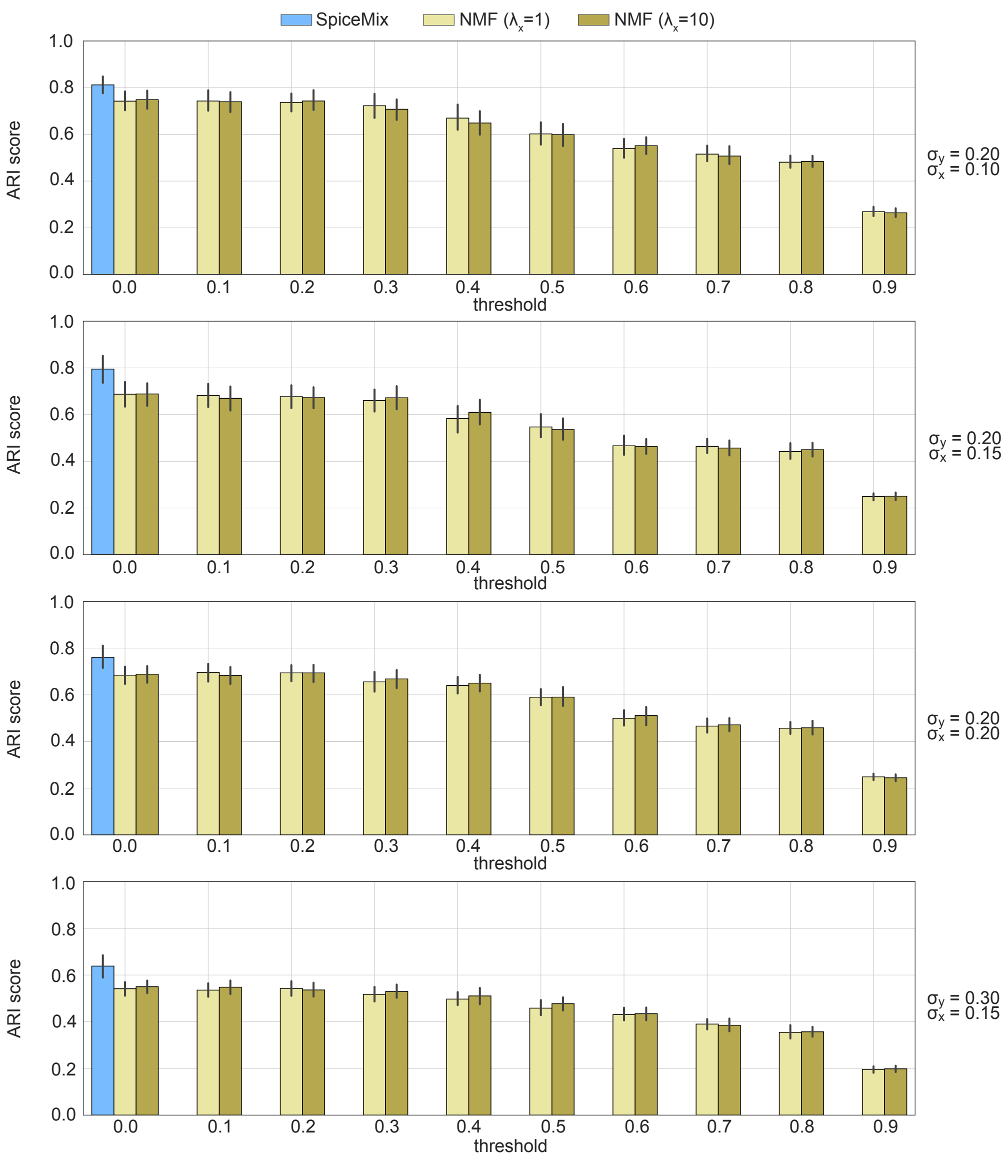


**Figure S1:** Comparison of SpiceMix with NMF variants on metagene-based synthetic datasets with different noise levels. NMF with different regularization strengths are distinguished by color. Boxes in each panel are grouped by the sparsity threshold which is labeled along the X-axis.

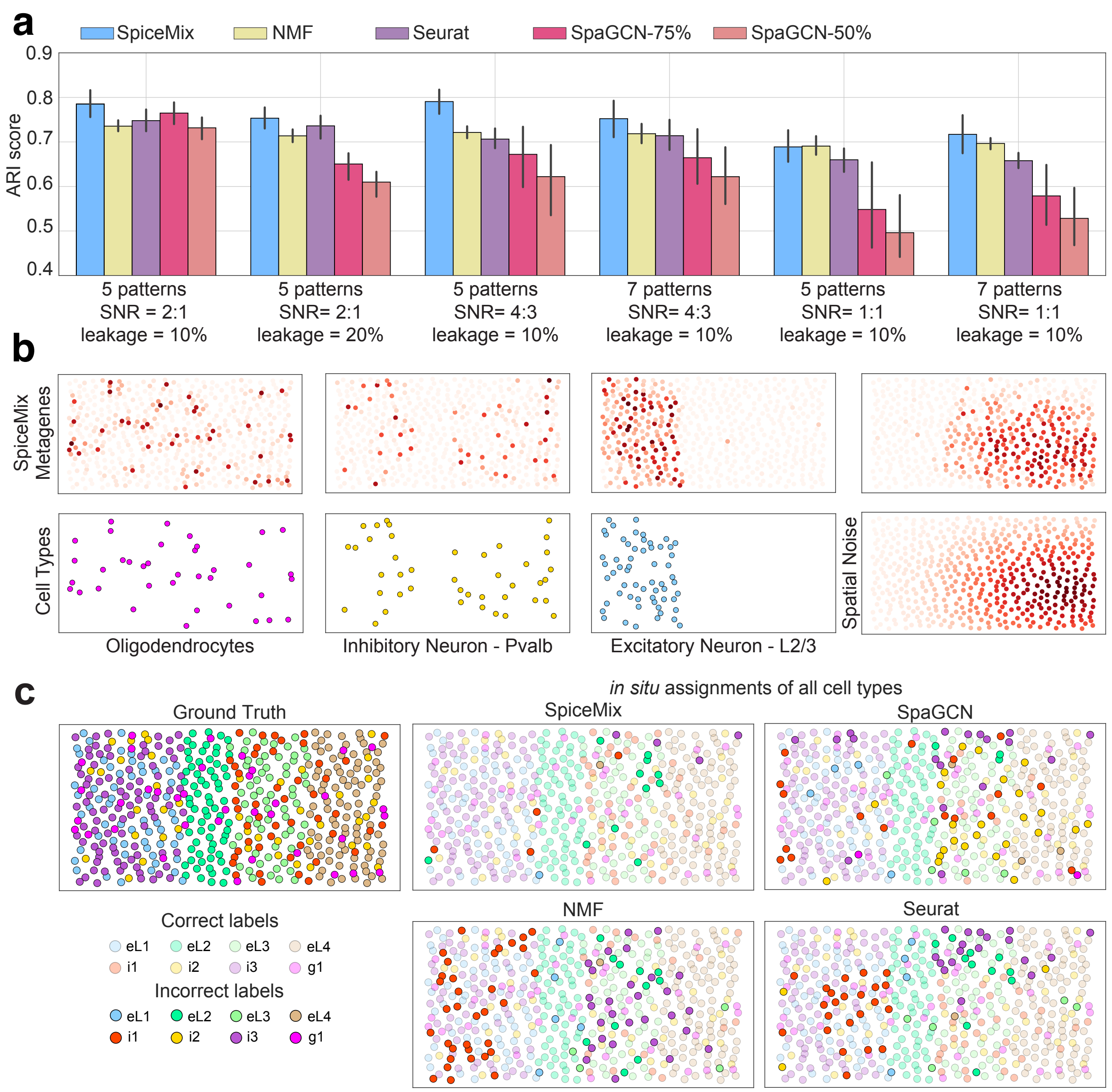


**Figure S2:** Overview of the simulated spatial transcriptome data of the mouse cortex using scDesign2, and performance comparison between SpiceMix, NMF, Seurat, and SpaGCN. **a.** Bar plots of the adjusted Rand index (ARI) score that measures the quality of the matching between the identified cell types for each method and the true simulated cell types. Results are reported across four simulation scenarios with varying degrees of randomness. The leakage percentage is the probability of a transcript being incorrectly assigned to a neighboring cell. The signal-to-noise ratio (SNR) is the ratio of the average true counts to the average counts generated by the spatial noise terms. The black error bars show +/- one standard deviation. **b.** Comparison of *in situ* expression of the learned metagenes of SpiceMix to *in situ* maps of cell types and an example spatial noise pattern. **c.** Assignments of excitatory neurons for each method in their spatial context. Colors were assigned to cells by the closest matching simulated type. Cells assigned to the incorrect cell types have bright colors. Cells assigned to the correct cell types have faint colors.

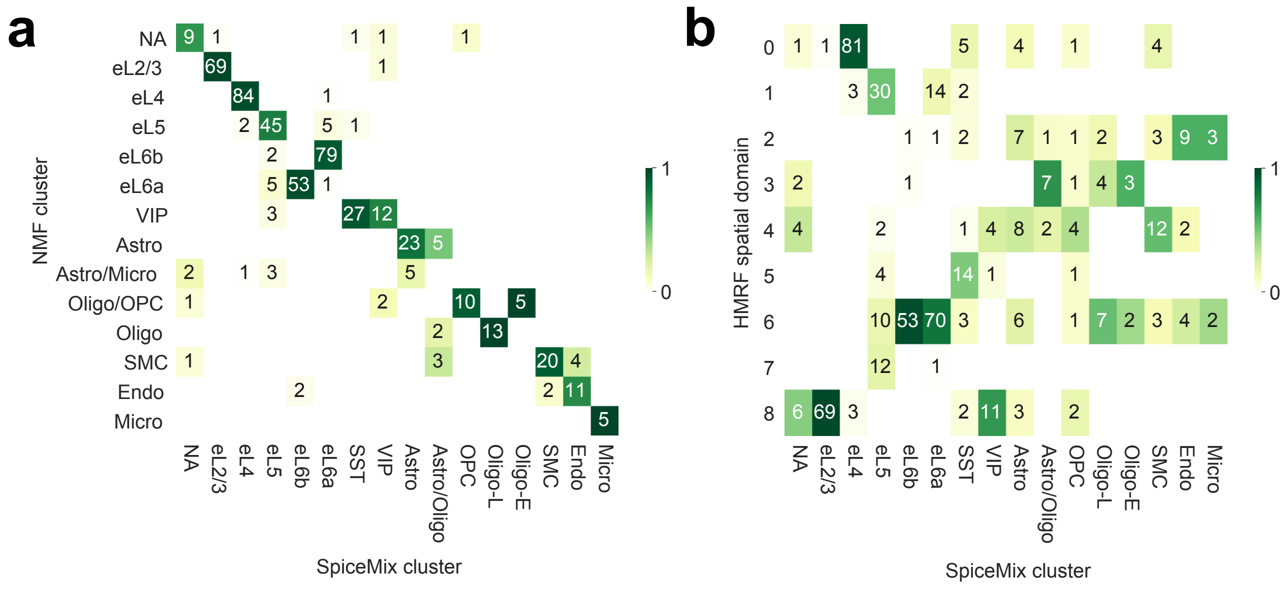


**Figure S3:** Additional details of the comparison of SpiceMix to NMF and to the HMRF-based method of Zhu et al. [13] on the seqFISH+ data [5]. **a.** The confusion matrix between SpiceMix and NMF cell-type assignments. **b.** The confusion matrix between SpiceMix and the spatial domains learned by the HMRF-based method of Zhu et al. [13], as reported in [5].

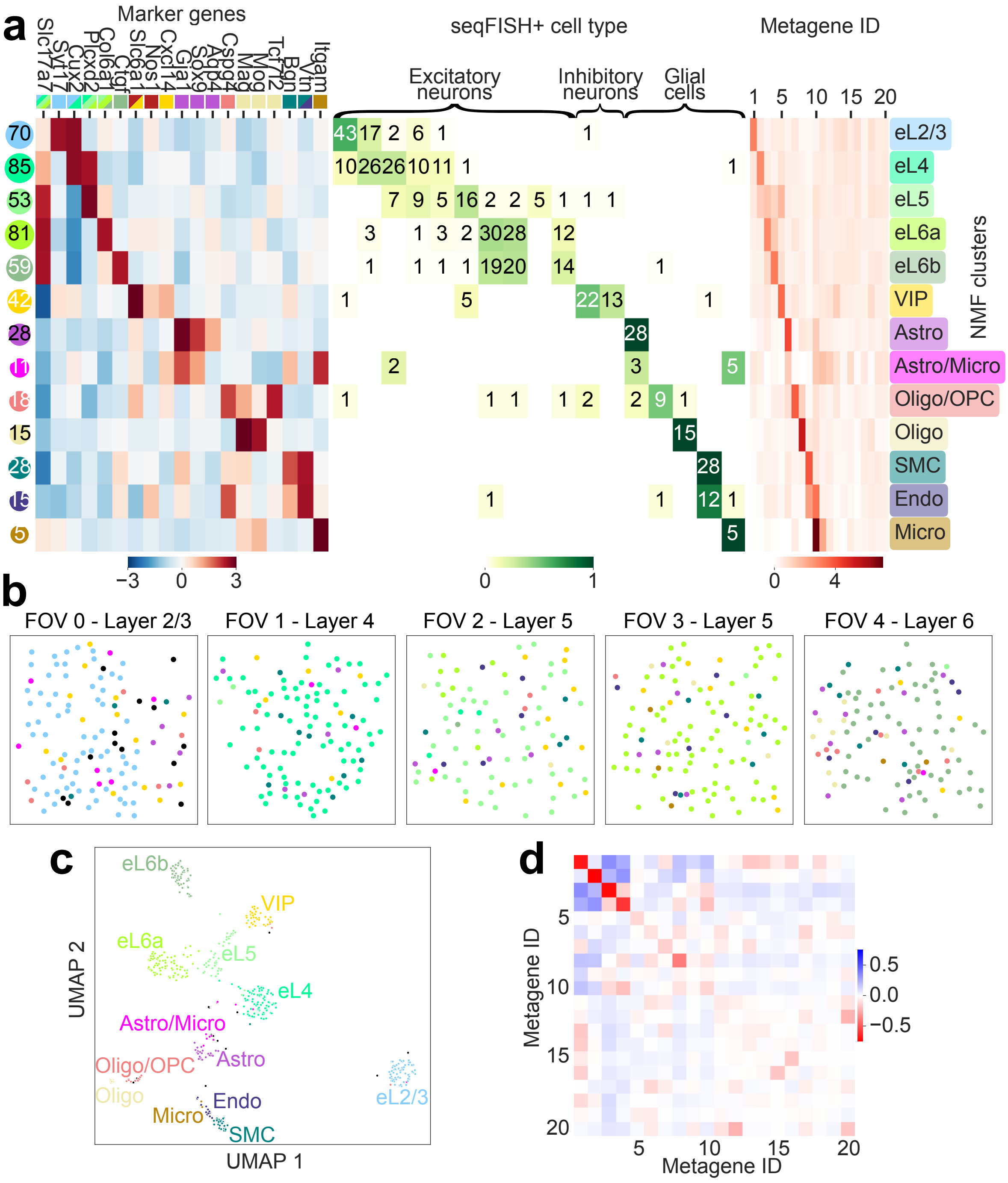

**Figure S4:** Characteristics of NMF cell types on the seqFISH+ data [5]. **a.** (Left) Average expression of known marker genes within NMF cell types. (Middle) Agreement of NMF cell types with clusters identified in [5]. (Right) Expression of metagenes within each cell type. **b.** Cell-type assignments of NMF *in situ*. **c.** UMAP plot of the cells in the latent representation learned by NMF. **d.** The pair-wise empirical correlation between the expression of NMF metagenes of neighboring cells. For all plots, the color correspond to those associated with the label names in panel **(a)**.

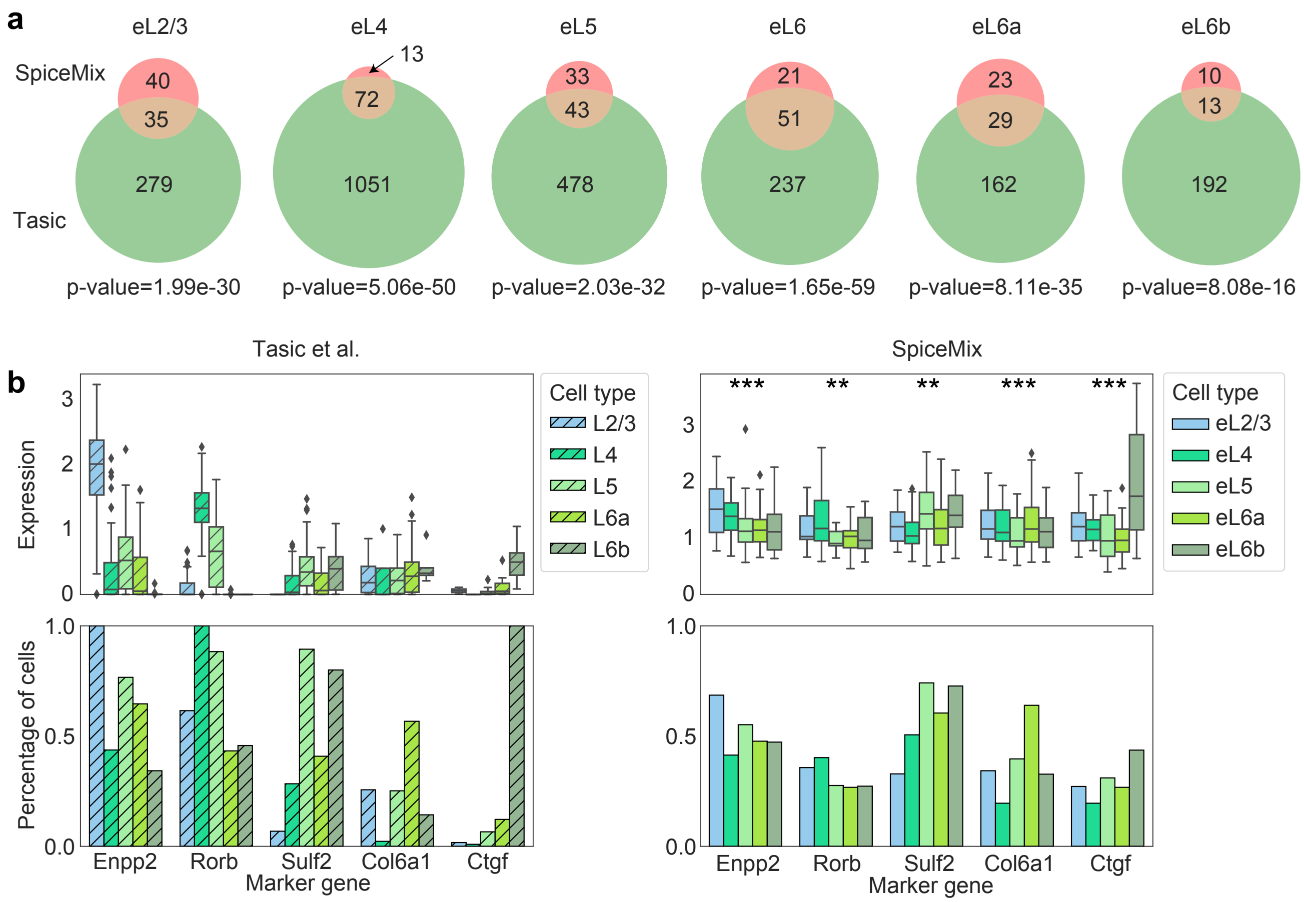


**Figure S5:** Additional justification of excitatory subtypes discovered by SpiceMix from the seqFISH+ dataset [5]. **a.** The overlap between DEGs of excitatory subtypes in SpiceMix and the counterparts in Tasic et al. [8]. The pink, orange, and green areas denote DEGs that are exclusively found between SpiceMix subtypes, that are found in both datasets, and that are exclusively found between subtypes in [8], respectively. **b.** A comparison of the expression of marker genes of excitatory neurons, reported in [8], between the scRNA-seq dataset of [8] and SpiceMix on the seqFISH+ dataset (top), and a comparison of the percentage of cells expressing those marker genes (bottom). $^{**}$ denotes one-vs-rest $P < 5 \times 10^{-3}$, and $^{***}$ denotes one-vs-rest $P < 5 \times 10^{-6}$.

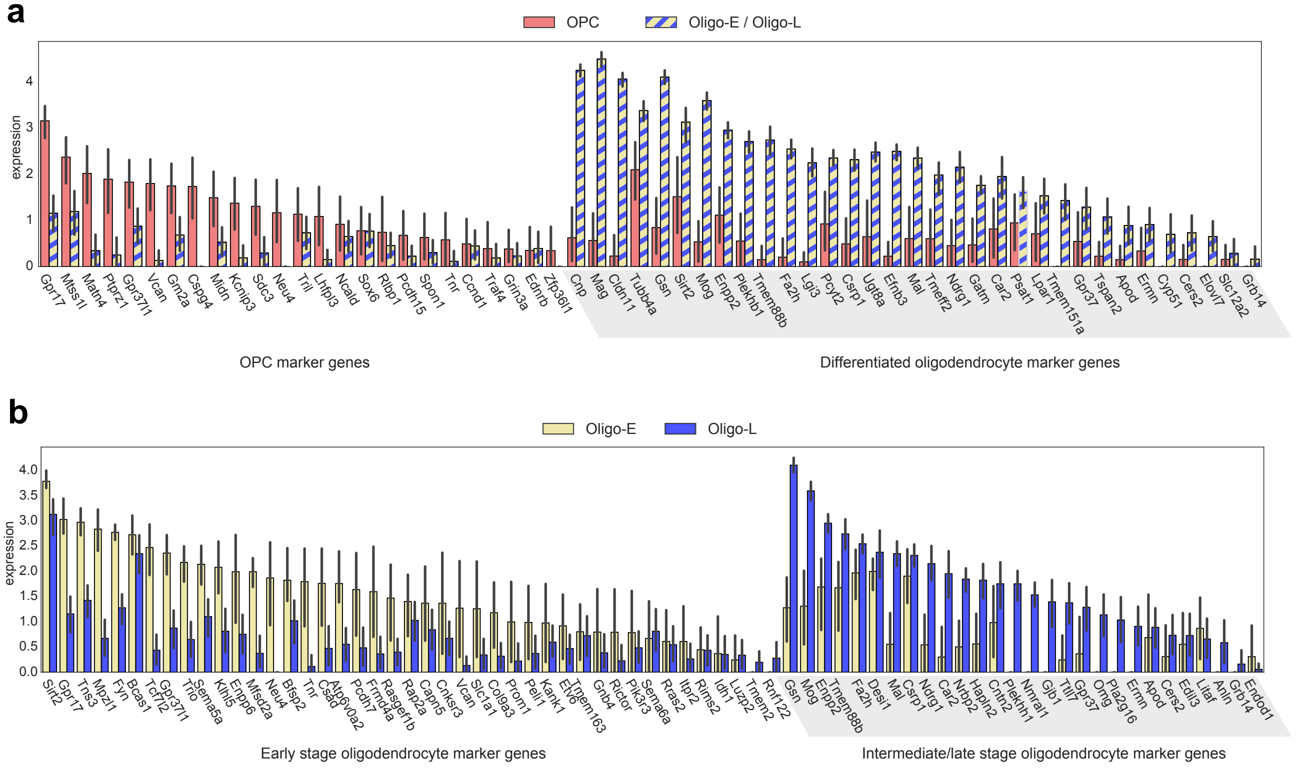


**Figure S6:** Additional justification of oligodendrocyte subtypes discovered by SpiceMix from the seqFISH+ dataset [5]. The average expression, after normalizing cell counts to 10,000 and applying the log-transform, are shown, with error bars showing +/- one standard deviation. **a.** The average expression of marker genes for OPCs and for differentiated oligodendrocytes, discovered by [29], in SpiceMix oligodendrocyte clusters based on the seqFISH+ data. The plots for "Oligo-E/Oligo-L" are the combined expression of cells from these two types. **b.** The average expression of marker genes for early stage and for intermediate/late stage oligodendrocytes, discovered by [29], in SpiceMix oligodendrocyte clusters based on the seqFISH+ data.

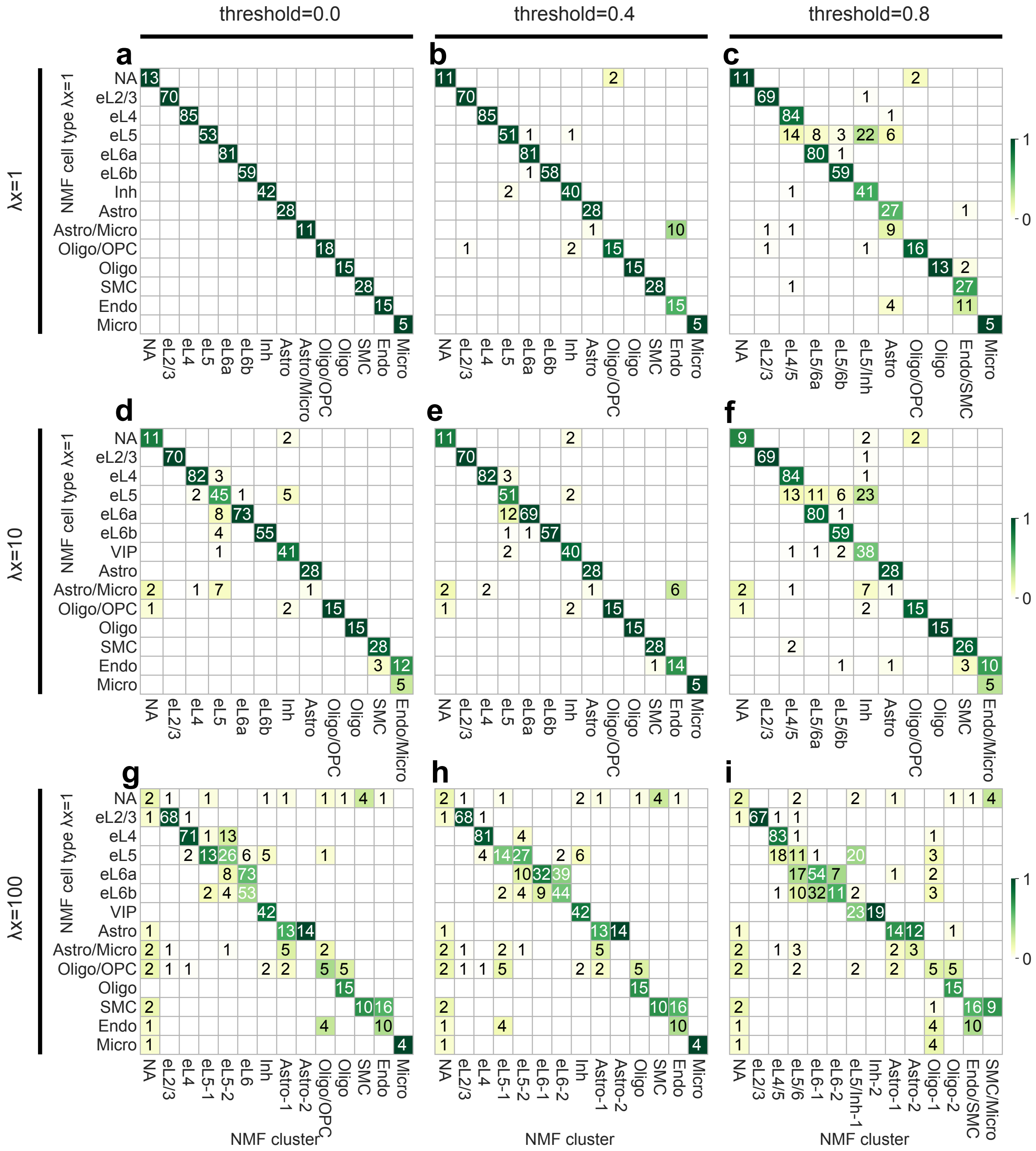

**Figure S7:** Comparison of cell type assignments on seqFISH+ between NMF variants with different regularization strengths $\lambda_x$ (rows) and different sparsity thresholds (columns). In each of the 9 panels, every row denotes one cluster of NMF with $\lambda_x = 1$ and without sparsity thresholding on the latent states, and columns represent clusters of NMF with the corresponding setting.

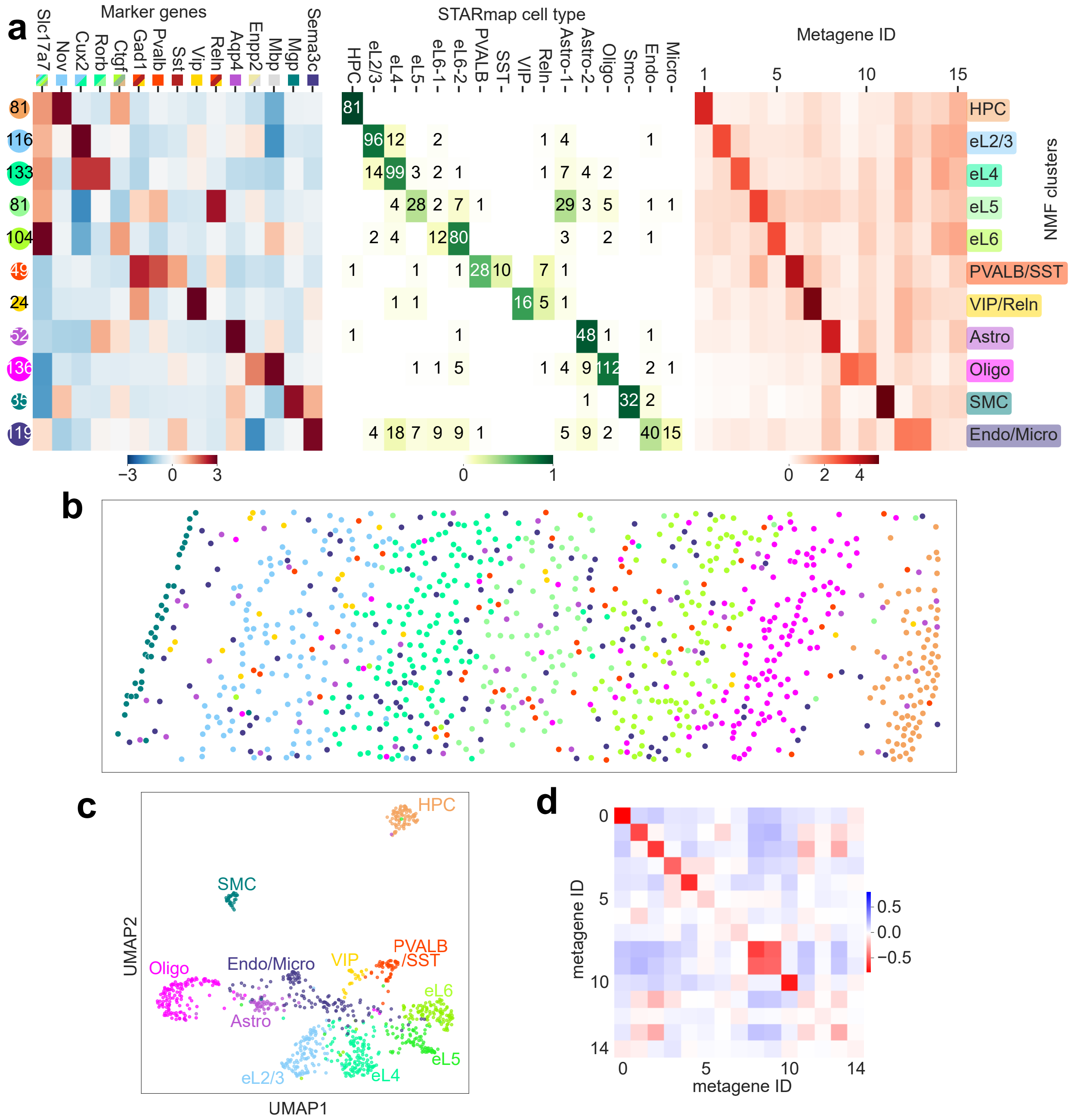

**Figure S8:** Characteristics of NMF cell types on the STARmap data [6]. **a.** (Left) Average expression of known marker genes within NMF cell types. (Middle) Agreement of NMF cell-type assignments with those from [6]. (Right) Expression of metagenes within each cell type. **b.** Cell-type assignments of NMF *in situ*. **c.** UMAP plot of the cells in the latent representation learned by NMF. **d.** The pair-wise empirical correlation between the expression of NMF metagenes of neighboring cells. For all plots, the color correspond to those associated with the label names in panel **(a)** (left).

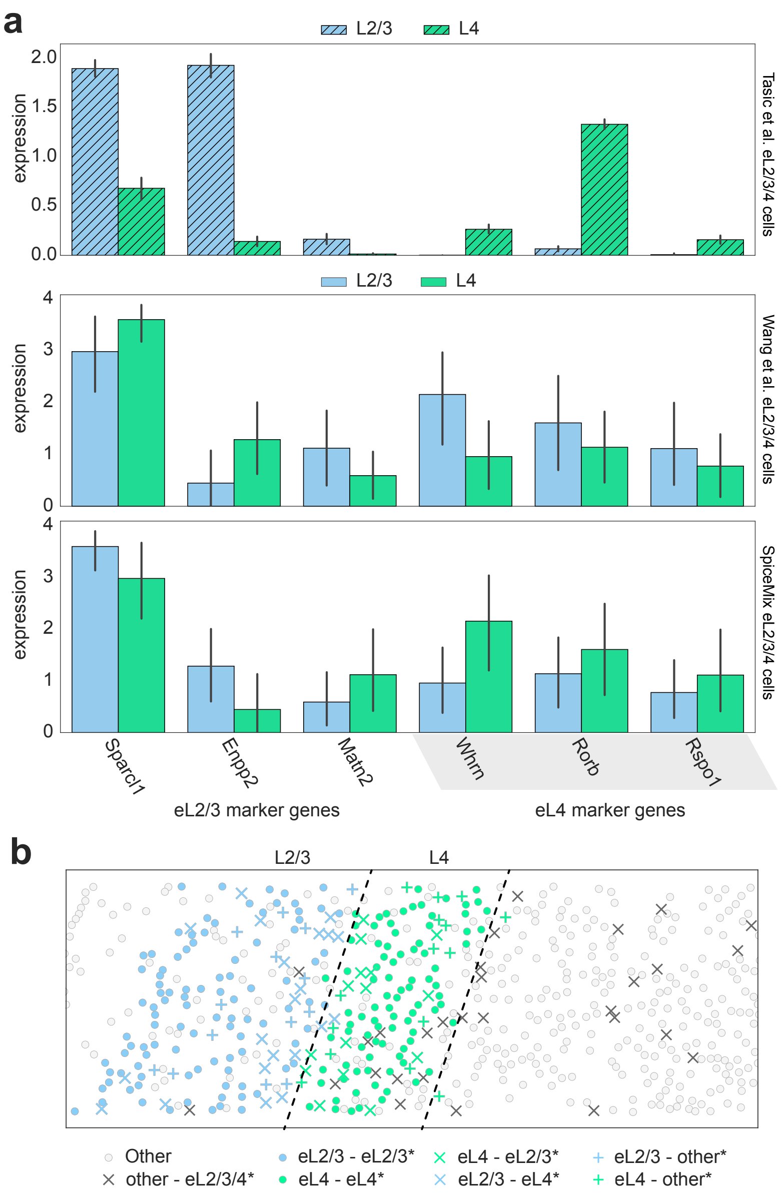


**Figure S9:** Comparison of excitatory neurons of Tasic et al. [8] with excitatory neuron assignments of Wang et al. [6] and SpiceMix on STARmap data. The average expression, after normalizing cell counts to 10,000 and applying the log-transform, are shown, with error bars showing +/- one standard deviation. **a.** (Top) The average expression of marker genes of eL2/3 and eL4 neurons, from the scRNA-seq dataset of Tasic et al. [8], as identified in their paper. (Middle) The average expression of eL2/3 and eL4 neurons assignments by SpiceMix from STARmap data. These are cells which Wang et al. [6] labeled as eL4* and eL2/3* neurons, respectively. (Bottom) The average expression of eL2/3* and eL4* neurons assignments by Wang et al. [6] from STARmap data. These are cells which SpiceMix labeled as eL2/3 and eL4 neurons, respectively. We considered only those marker genes from [8] that were expressed in at least 20% of the cells considered from the STARmap dataset. **b.** Comparison of *in situ* patterns of eL2/3 and eL4 neuron assignments of Wang et al. [6] and SpiceMix. The blue '×' indicates cells labeled as eL4* neurons by Wang et al. [6] but changed to eL2/3 neurons by SpiceMix. The green '×' indicates cells labeled as eL2/3* neurons by Wang et al. [6] but changed to eL4 neurons by SpiceMix. The blue '+' indicates cells labeled as other cell types by Wang et al. [6] but changed to eL2/3 neurons by SpiceMix. The green '+' indicates cells labeled as other cell types by Wang et al. [6] but changed to eL4 neurons by SpiceMix. The black '×' indicates cells labeled by Wang et al. [6] as eL2/3* or eL4* neurons, but labeled as an other cell type by SpiceMix. All circles indicate consistent eL2/3, eL4, or other cell type labels between Wang et al. [6] and SpiceMix.

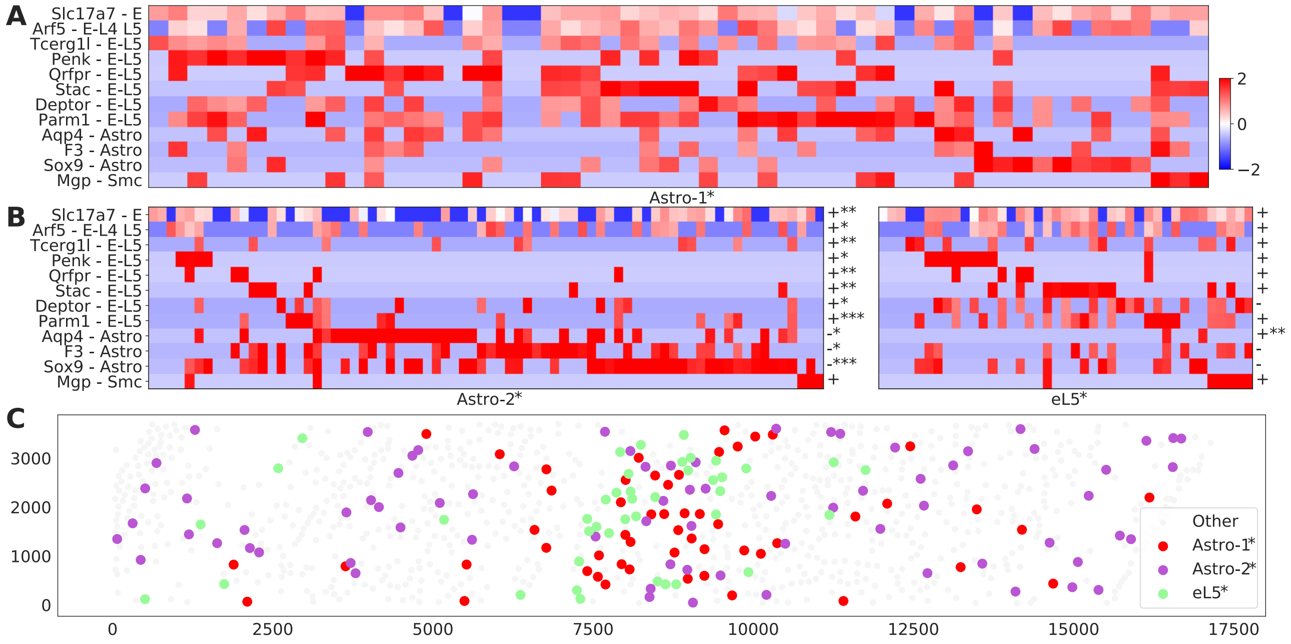


**Figure S10:** Characteristics of Astro-1*, Astro-2*, and eL5* from the original analysis of the STARmap data in [6]. **a.** Expression profiles of cells in Astro-1*, z-score normalized per gene. **b.** Expression profiles of cells in Astro-2* and eL5*, z-score normalized per gene. The symbols tn the right of each subplot indicate the difference of gene expressions between Astro-1* and the cell type showed below the subplot, and their $P$-values. Specifically, a plus/minus sign indicates Astro-1* cells have a higher/lower expression, and the number of asterisks represents the $P$-value: * for $P<0.01$, ** for $P<0.001$, *** for $P<0.0001$. **c.** Spatial distribution of Astro-1*, Astro-2*, and eL5* cells.

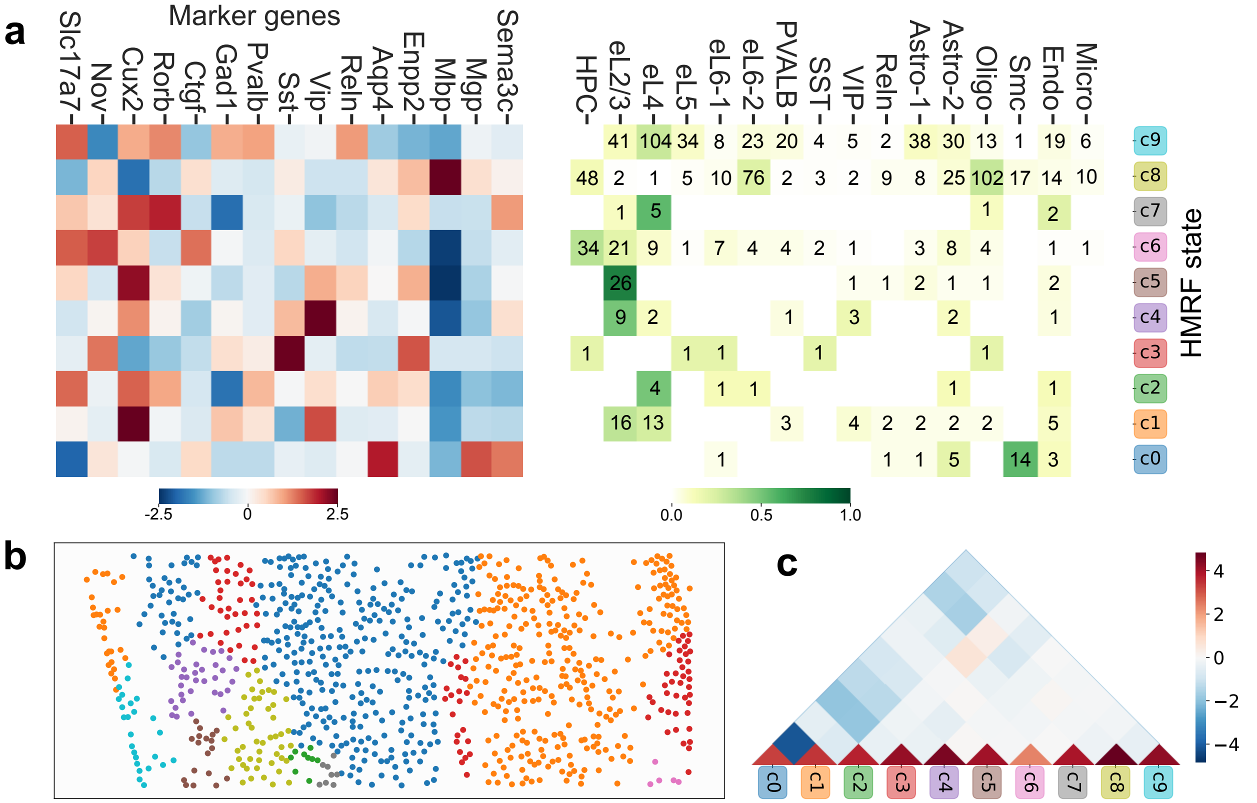


**Figure S11:** Characteristics of HMRF spatial patterns based on the STARmap data [6]. **a.** The numbers of cells in HMRF clusters. **b.** Selected marker gene profiles of HMRF clusters. **c.** Confusion matrix between STARmap cell types (rows) and HMRF clusters (columns). **d.** Spatial distributions of cells colored by HMRF clusters. **e.** Affinities between HMRF clusters.

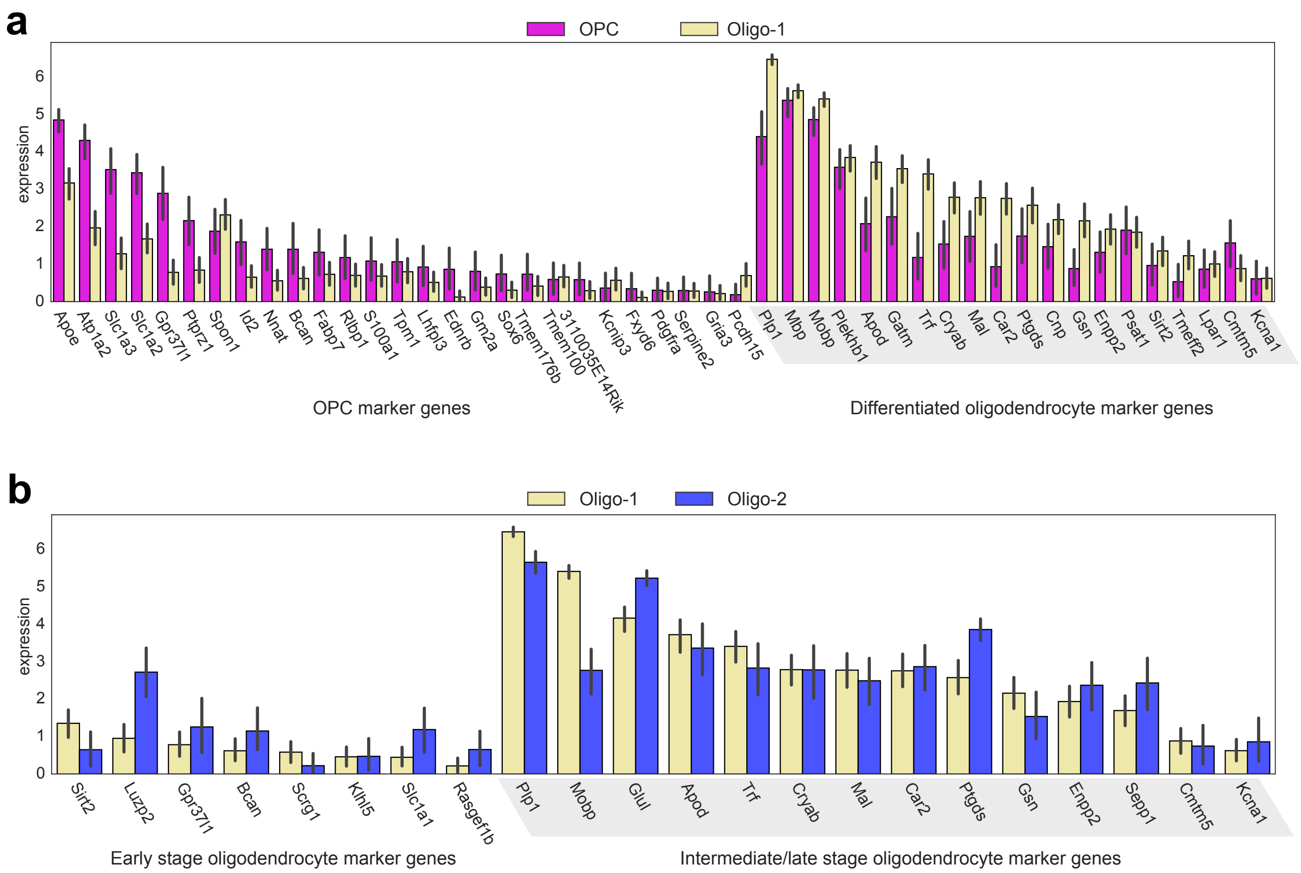


**Figure S12:** Characteristics of the OPC type and two oligodendrocyte subtypes identified by SPICEMIX on the STARmap data [6]. The average expression, after normalizing cell counts to 10,000 and applying the log-transform, are shown, with error bars showing +/- one standard deviation. **a.** The average expression of DEGs that split OPCs and differentiated oligodendrocytes, discovered by [29], in SPICEMIX OPC and Oligo-1 clusters based on the STARmap data. **b.** The average expression of DEGs that split early and intermediate/late stage oligodendrocytes, discovered by [29], in SPICEMIX oligodendrocyte clusters based on the STARmap data.

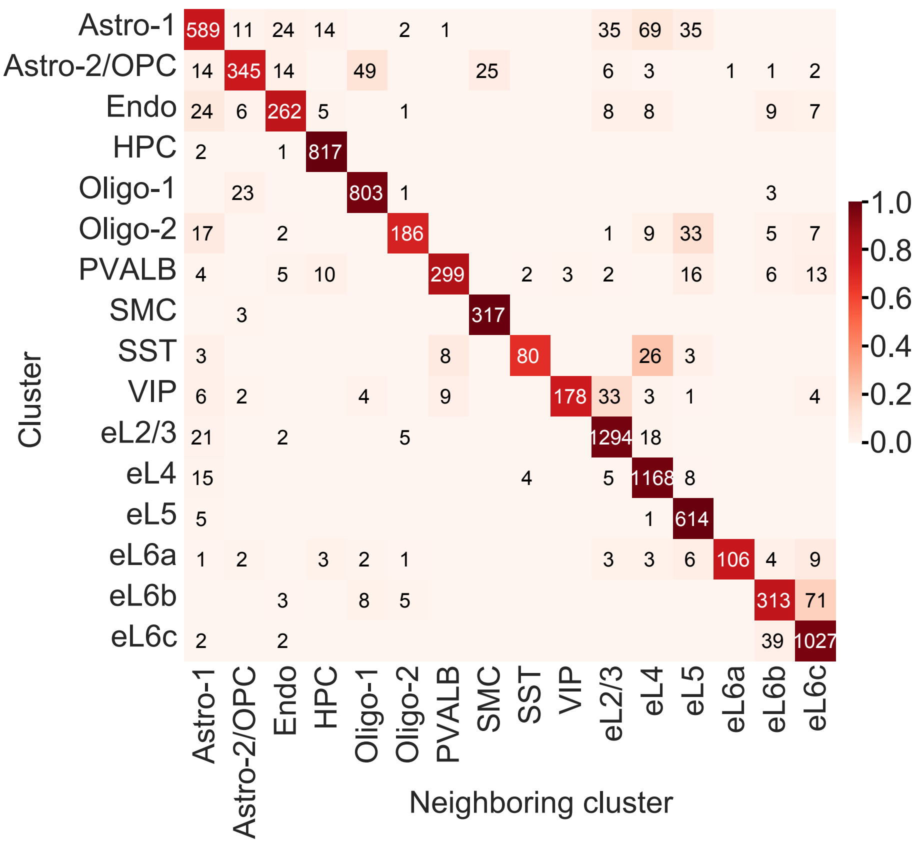


**Figure S13:** Additional justification of the separation of cell types in the latent space learned by SpiceMix on the STARmap dataset [6]. The figure shows the adjacency matrix for the 10-nearest neighbor graph of SpiceMix cell embeddings, aggregated by cell types. Each row shows the cell type distribution of the neighboring cells of one particular cell type, in the embedding space. Numeric annotations are the number of directed edges in the 10-nearest neighbor graph. The color represents the adjacency matrix normalized per row.

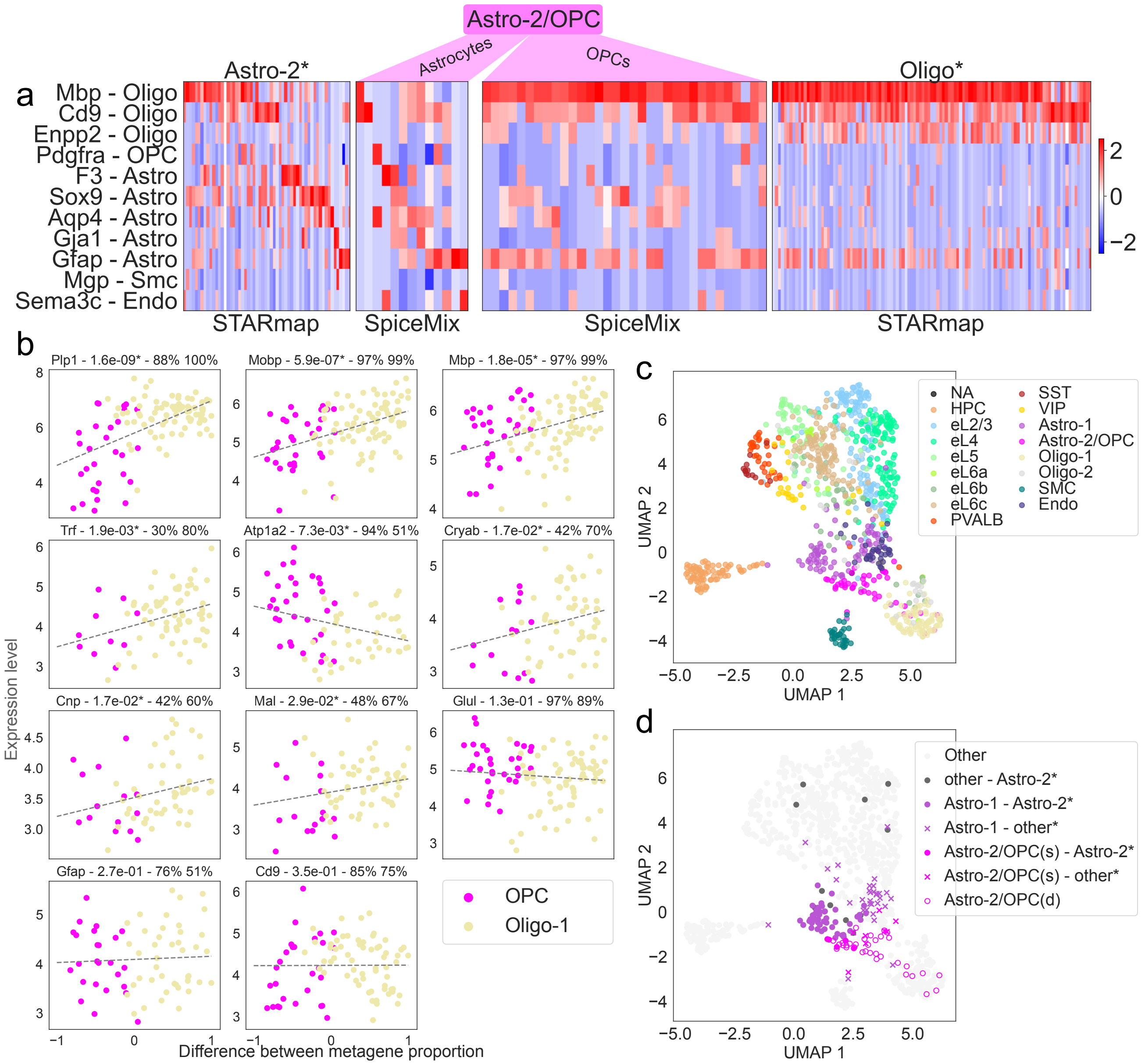


**Figure S14:** Extended analysis of the Astro-2/OPC cluster revealed by SPICEMIX from the STARmap data [6]. **a.** Comparison of the expression of oligodendrocyte, astrocyte, SMC, and endothelial cell marker genes among the Astro-2* and Oligo* clusters of [6] and the Astro-2/OPC cluster of SPICEMIX. **b.** Relation between the expression of 11 myelin sheath-related genes from the STARmap data and proportions of expression of metagenes learned by SPICEMIX. Each plot shows the expression levels against the metagene proportion differences in individual cells colored according to SPICEMIX cell type assignment. A cell with x-coordinate -1 expresses metagene 12 exclusively, while a cell with x-coordinate +1 expresses metagene 13 exclusively. The dashed lines are the fitted linear regression model. The title of each plot consists of the gene symbol, the corrected $p$-value of having a nonzero slope, and the fractions of OPCs and oligodendrocytes with nonzero expression levels, respectively. An asterisk after the $p$-value means that the result is significant under the threshold of 0.05. **c.** UMAP plot of the STARmap cells based on their expression profiles, colored according to SPICEMIX cell labels, showing that the Astro-2/OPC cells lie between the Astro-1 and Oligo-1 cells. **d.** UMAP plot of the STARmap cells based on their expression profiles, with labels showing the agreement or disagreement between SPICEMIX labels and the labels of [6]. The Astro-2/OPC cells are separated into astrocytes and oligodendrocytes based on their spatial location. They are labeled as Astro-2/OPC(s) and Astro-2/OPC(d), respectively, because astrocytes are in superfacial layers, while oligodendrocytes are in deeper layers.

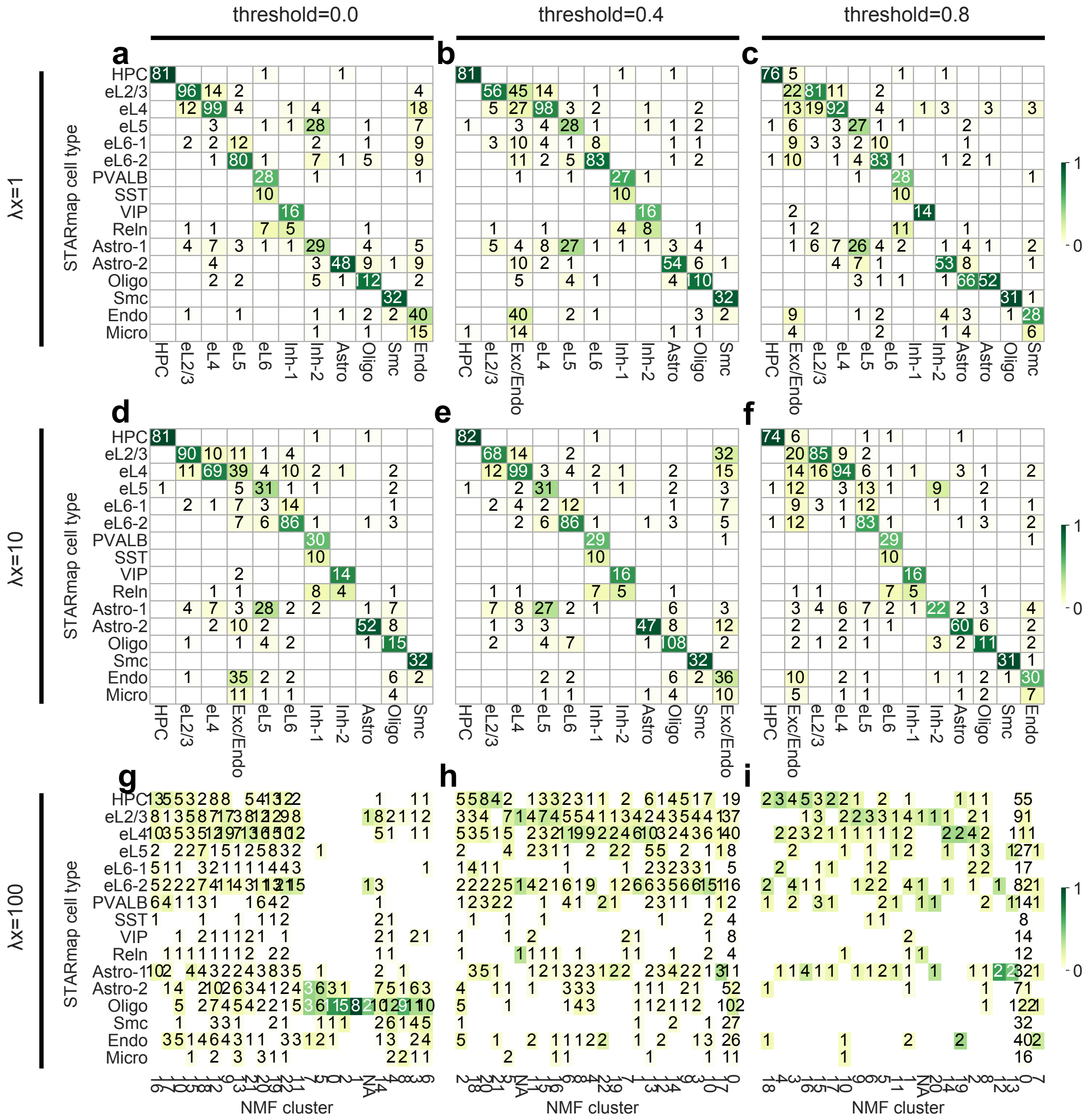

**Figure S15:** Comparison of cell type assignments on STARmap between the Wang et al. [6] and NMF variants with different regularization strengths $\lambda_x$ (rows) and different sparsity thresholds (columns). In each of the 9 panels, every row denotes one cell type in Wang et al. [6], and columns represent clusters of NMF with the corresponding setting.

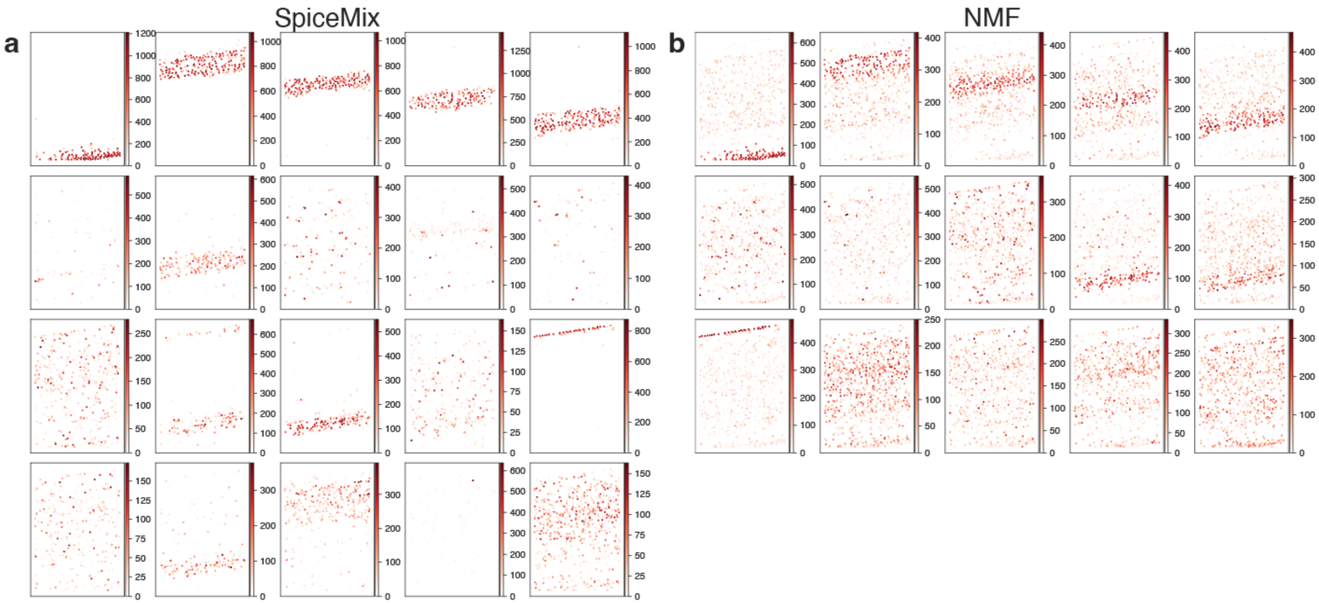


**Figure S16:** The spatial distribution of metagenes from SPICEMIX (**a**) and from NMF **(b)** on the STARmap dataset [6].

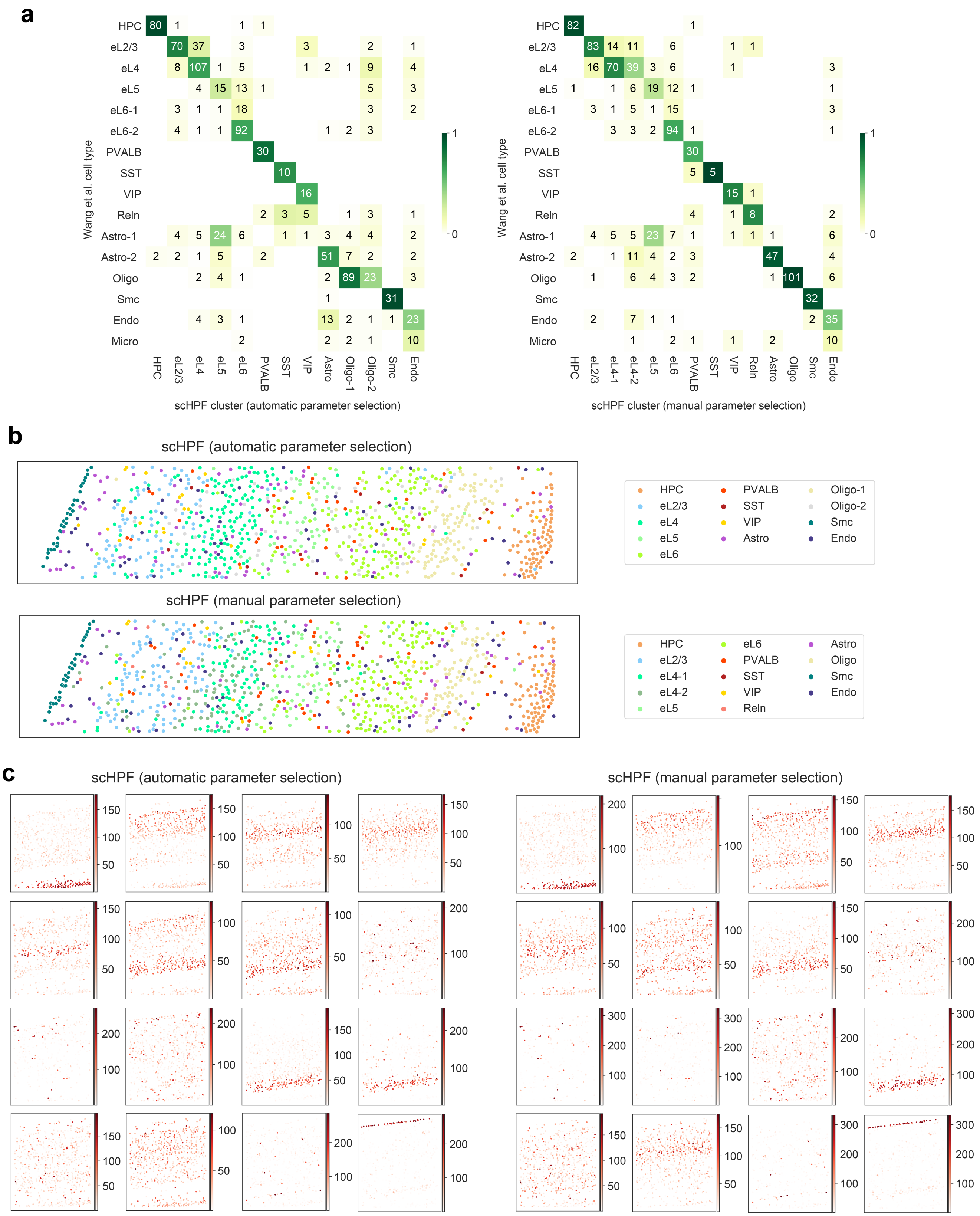


**Figure S17:** Results of scHPF [24] on the STARmap data [6]. **a.** Confusion matrix showing agreement between scHPF clusters and cell-type assignments of [6]. **b.** *In situ* map of cells and imputed cluster assignments. **c.** *In situ* expression of inferred latent factors.

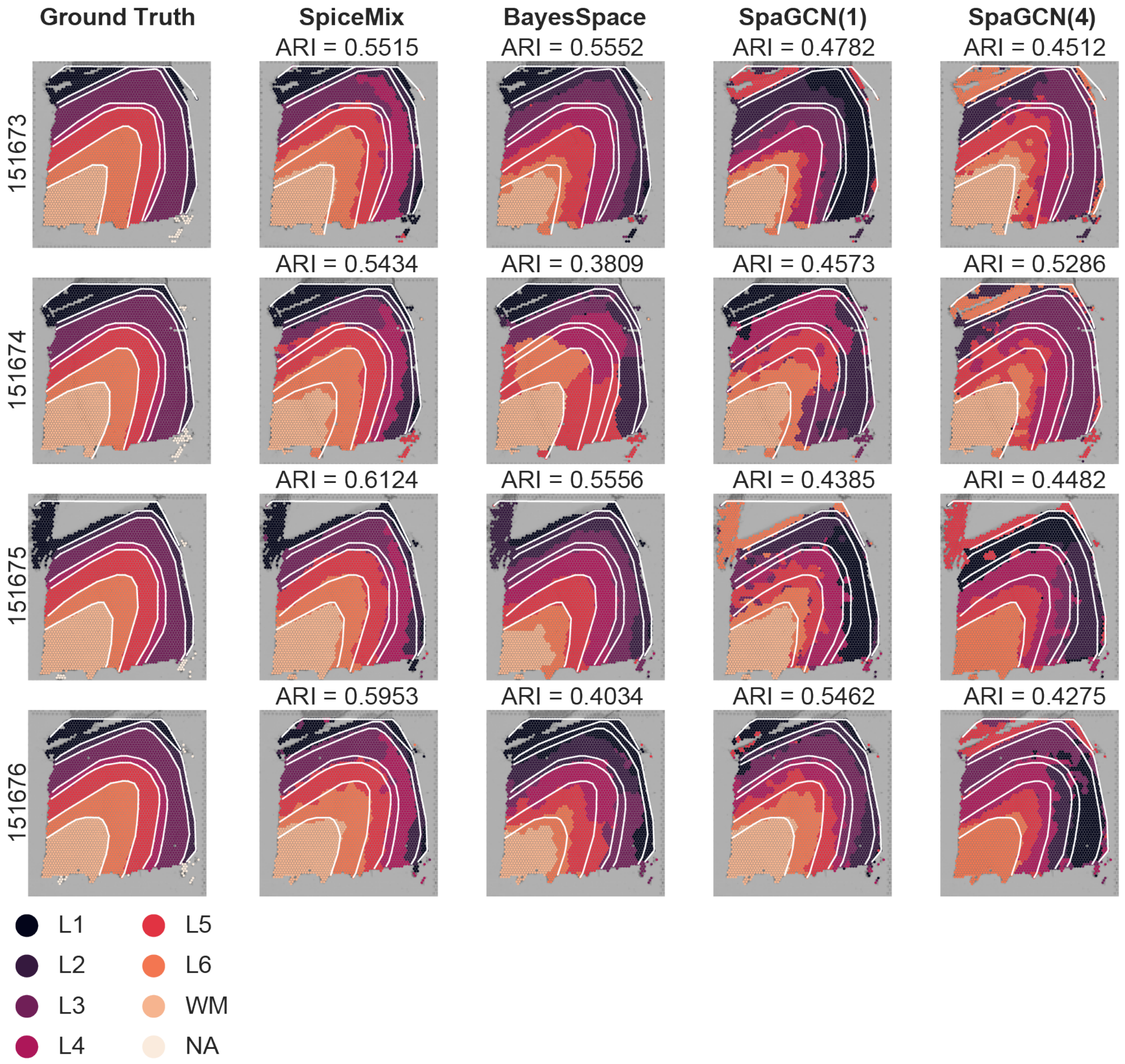


**Figure S18:** *In situ* plots of layer annotations of *ground truth*, SPICEMIX, BayesSpace, SpaGCN(1), and SpaGCN(4) (one column per method), on FOVs 151673-151676 (one row per FOV) of the human DLPFC Visium dataset [30]. 20 different random seeds were tried for BayesSpace and SpaGCN(1), and 10 random seeds were tried for SpaGCN(4). For each of BayesSpace, SpaGCN(1), and SpaGCN(4), the random seed that attained the best ARI scores is shown. White lines illustrate layer boundaries that were derived from the ground truth. The colors in the first column represent the *ground truth* annotations, and the legend is at the bottom. The colors in the rest three columns do not have the same meaning as in the first column. The 4 maps in the second column (SPICEMIX) share the same color scheme, as we trained SPICEMIX and did clustering on all 4 FOVs at the same time. The maps in the third column (BayesSpace) and in the fourth column (SpaGCN(1)) do not share the same color scheme, because BayesSpace and SpaGCN(1) were trained on each FOV separately. The 4 maps in the last column (SpaGCN(4)) share the same color scheme, as we applied SpaGCN() on all 4 FOVs at the same time.

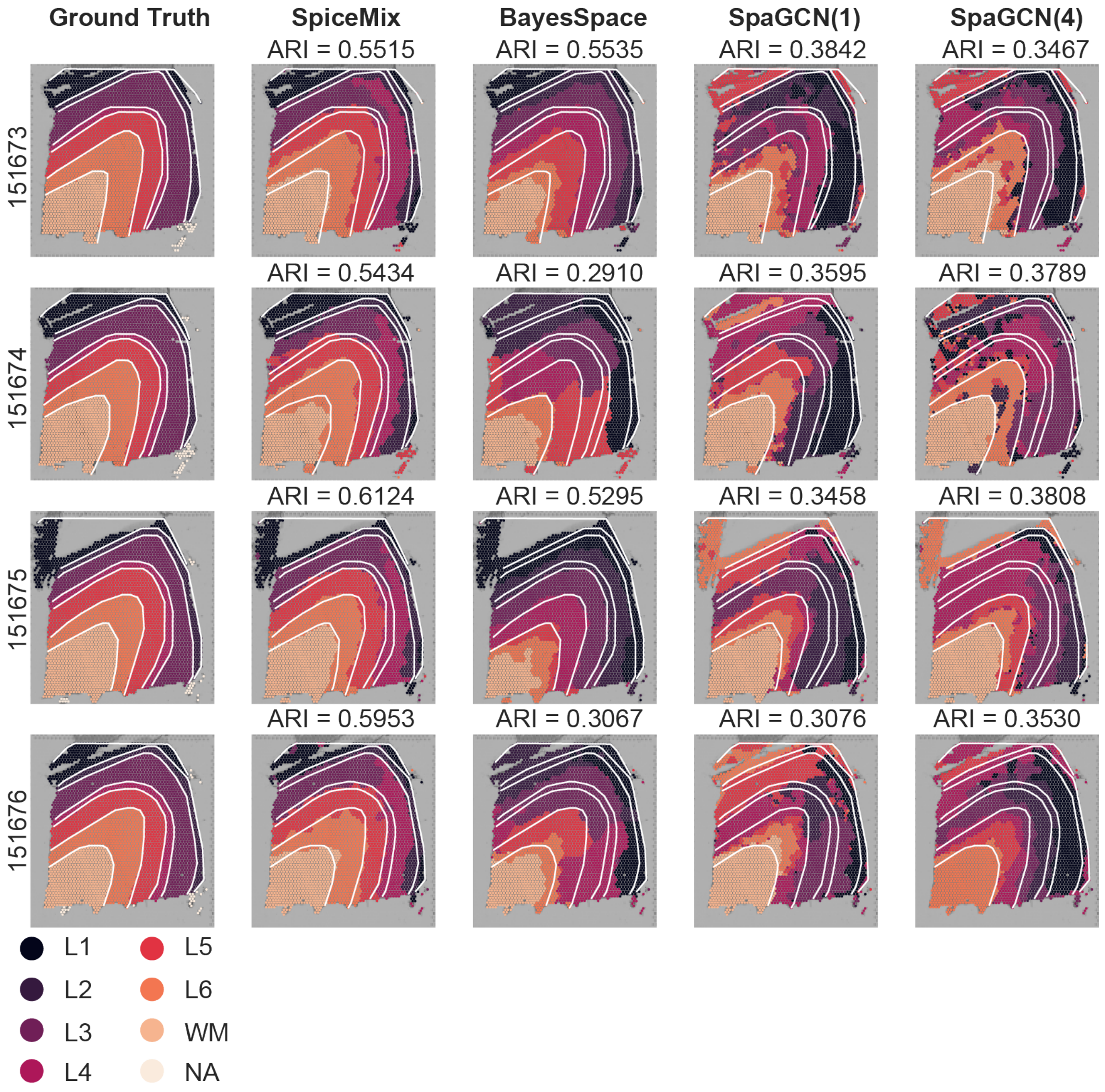


**Figure S19:** See the caption of **Fig.** S18 with the same explanation except that the *in situ* maps that attained the median ARI scores are shown for BayesSpace, SpaGCN(1), and SpaGCN(4). Specifically, the map of the 11-th best random seed was shown for BayesSpace and SpaGCN(1), and the map of the 6-th best random seed was shown for SpaGCN(4).

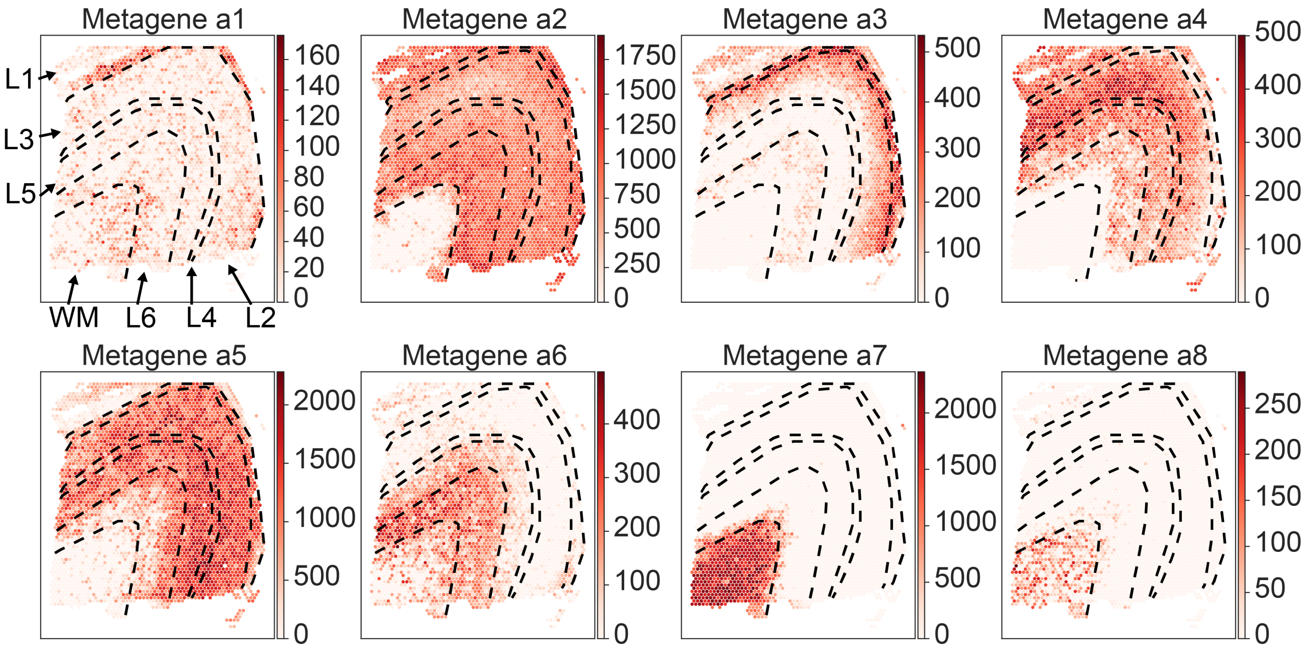


**Figure S20:** *In situ* plots of the expression of metagenes from SpiceMix for FOV 151673 of the human DLPFC Visium dataset [30]. The 8 metagenes are those with highest expression levels.

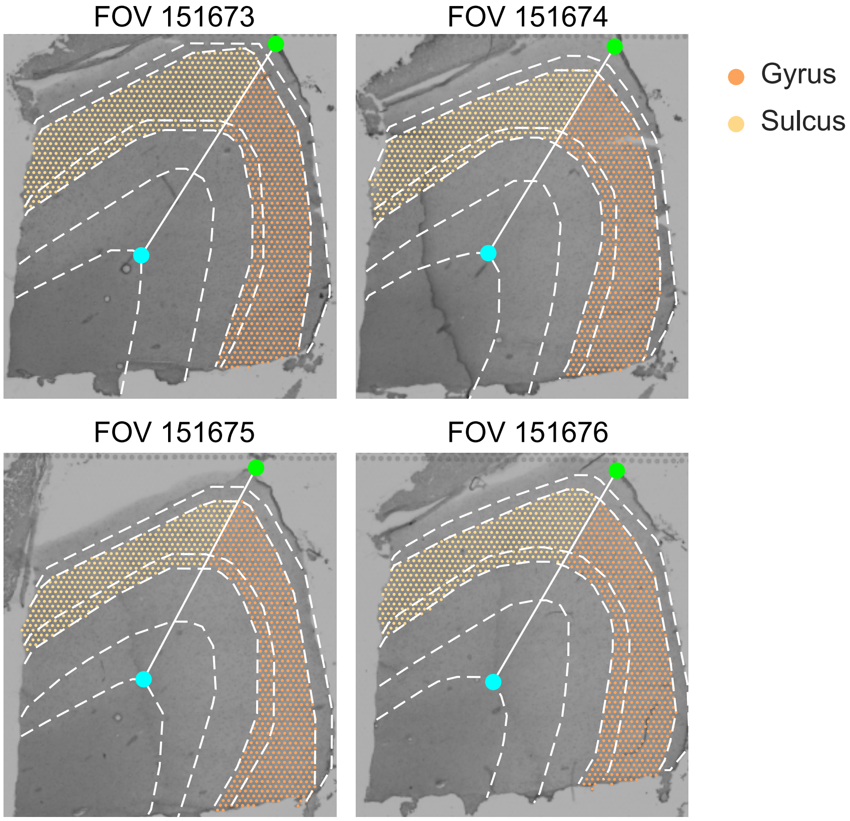


**Figure S21:** Illustration of the approximate separations of layers 3 and 4 to the gyric side and the sulcal side of the human DLPFC Visium dataset [30]. The color scheme matches the one used in **Fig.** 5b.

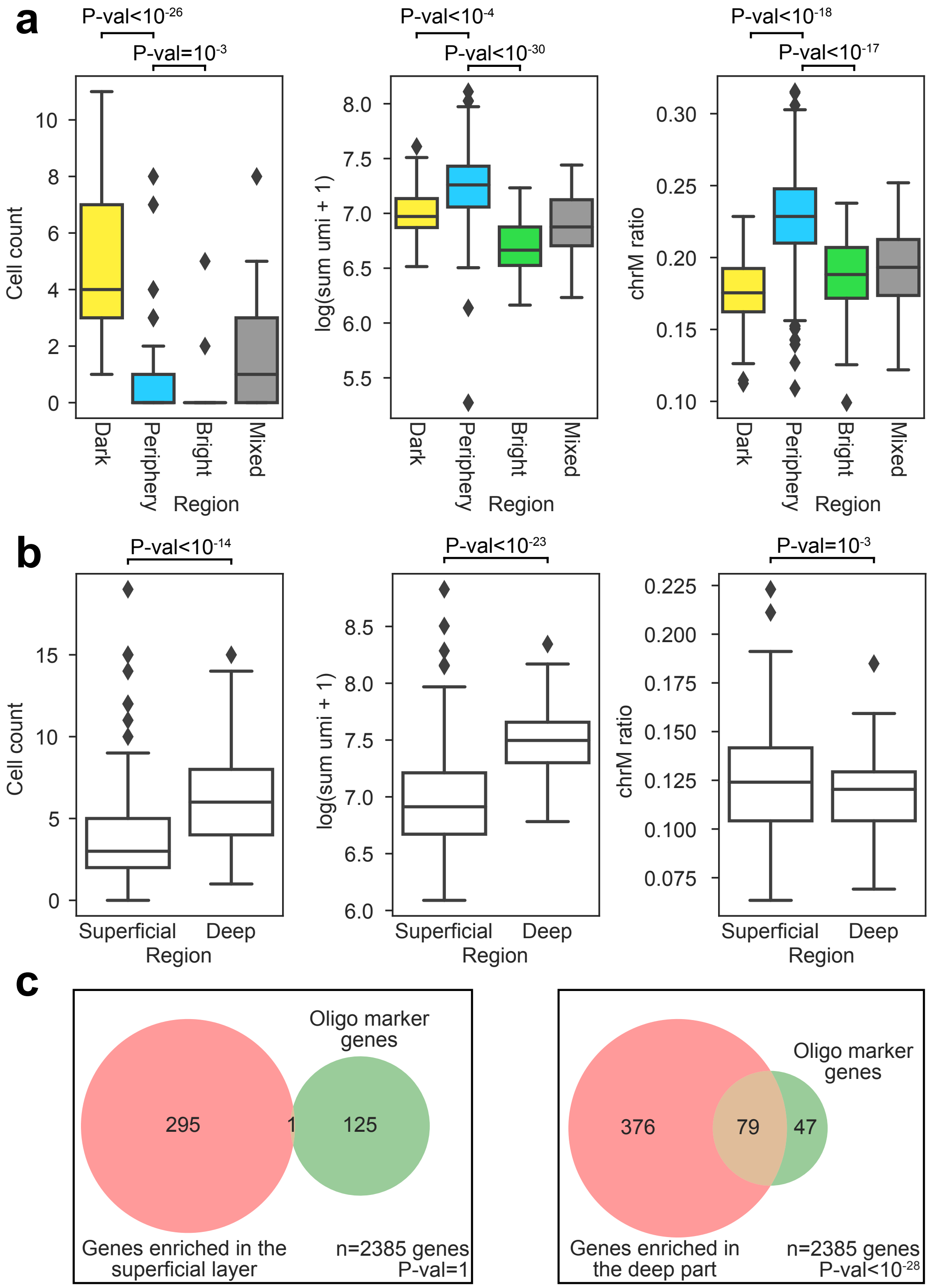


**Figure S22:** Additional evidence of the biological differences between the finer structures in L1 (panel **a**), and between the superficial layer and the deep part of the white matter (panels **b** and **c**), in sample FOV 151507 of the human DLPFC Visium dataset [30]. **a.** The overlap between the marker genes of oligodendrocytes and genes enriched in the superficial layer (left) and in the deep part (right) of the white matter in FOV 151507. **b.** Comparison of the distributions of 3 phenotypes in the superficial layer and the deep region of the white matter in FOV 151507. **c.** Comparison of the distributions of 3 phenotypes in the dark stripe, the bright gap, the flanking tissue, and the ambiguous spots of layer 1 in FOV 151507.
